# Transcription Factor Subtype Governs Response and Resistance to DLL3-Directed T-Cell Engagement in Small Cell Lung Cancer

**DOI:** 10.64898/2026.04.02.715020

**Authors:** Damien Vasseur, Shin Saito, Gunsagar S. Gulati, Garyoung Gary Lee, Yasmin N. Laimon, Berkay Simsek, Maddie Lerner, Hyeonseo Cho, Yixiang Li, Tianchu Wang, Ji-Heui Seo, Hunter Savignano, Brady James, Ze Zhang, Karl Semaan, Zhenjie Jin, Wassim Daoud Khatoun, Gaelle Nafeh, Rashad Nawfal, Alissa J. Cooper, Kathryn Miller, Maxwell D. Seager, Elliott J. Brea, Eric Smith, Jonathan Chang, Marc Pelletier, Carlotta Costa, Toni K. Choueiri, Sabina Signoretti, Jacob Sands, Sylvan C. Baca, Matthew L. Freedman, Matthew G. Oser

## Abstract

Although small cell lung cancer (SCLC) comprises transcription factor (TF)-defined molecular subtypes (ASCL1, NEUROD1, POU2F3), the extent to which these subtypes predict response to clinically effective therapy in patients—and whether therapy can select for subtype switching—remains unknown. The recent approval of the DLL3×CD3 bispecific T-cell engager tarlatamab represents one of the first meaningful advances in relapsed small cell lung cancer (SCLC) in decades, yet responses remain heterogeneous and resistance is inevitable. Here, we inferred SCLC gene expression from circulating chromatin in prospectively collected patient plasma (46 patients; 167 samples), enabling interrogation of response and acquired resistance to tarlatamab. Parallel development of the first immunocompetent syngeneic mouse model to study tarlatamab response and resistance enabled functional validation. Across species, findings converged on a central principle: TF subtype governs both initial response and acquired resistance. Therapeutic response was significantly associated with ASCL1-subtype tumors, whereas NEUROD1-subtype tumors exhibited inferior responses and POU2F3-subtype tumors were uniformly resistant, consistent with DLL3 being a direct ASCL1 transcriptional target and most highly expressed in ASCL1-positive tumors. Strikingly, one mode of acquired resistance revealed therapeutic selection for a NEUROD1-high state with concomitant DLL3 downregulation. Other resistant tumors exhibited enrichment of regulatory and exhausted T-cell programs, highlighting tarlatamab’s dual-targeting mechanism of action. Together, these results reveal that tarlatamab exerts selective pressure against ASCL1-driven lineages, facilitating resistance through loss of an antigen intrinsically linked to that state. These findings underscore the clinical relevance of TF-defined molecular subtypes in human SCLC. More broadly, they highlight the power of integrating longitudinal *in vivo* plasma transcriptional profiling from patient plasma with functional mouse modeling to uncover clinical and biological mechanisms of response and resistance to cell-surface-targeted therapies.

## INTRODUCTION

Small-cell lung cancer (SCLC) is a high-grade neuroendocrine cancer accounting for ∼13% of lung cancers and remains one of the most aggressive malignancies characterized by rapid growth, early dissemination, and relapse in over 90% of patients following initial combined chemo-immunotherapy^1^. Recently, the development of DLL3-directed immunotherapy has meaningfully altered the treatment landscape^2–4^. Tarlatamab, a bispecific T-cell engager targeting delta-like ligand 3 (DLL3) on tumor cells and CD3 on T cells, promotes T-cell-mediated cytotoxicity and has demonstrated clinically significant activity in relapsed SCLC^2–4^. In the phase II DELLPHI-301 trial, tarlatamab achieved objective responses in approximately 40% of patients, with over half of responders maintaining benefit beyond six months^2^. Subsequent phase III data (DELLPHI-304) demonstrated improved overall survival compared with second-line chemotherapy^3^, establishing tarlatamab as the standard second-line therapy. However, most patients do not experience durable responses^2^, highlighting an urgent need to define biomarkers of response and mechanisms of resistance.

DLL3 can act as an inhibitory ligand in the NOTCH pathway^5–8^ and is a direct downstream transcriptional target of the lineage transcription factor ASCL1^9,10^. In SCLC, DLL3 is aberrantly expressed on the tumor cell surface, whereas DLL3 expression in normal tissues is typically low and rarely localized on the cell surface^11^. These properties have made DLL3 an attractive therapeutic target in SCLC and across high-grade neuroendocrine malignancies^12^.

SCLC is increasingly recognized as a heterogeneous disease comprised of molecular subtypes defined by dominant transcription factor programs, including the ASCL1-dominant subtype (∼60%), NEUROD1-dominant subtype (∼20%), POU2F3-dominant subtype (∼10%), and an inflammatory subtype lacking these lineage drivers^13,14^. DLL3 expression is highest in ASCL1-positive SCLC tumors, variably present in NEUROD1-positive tumors, and typically low or absent in POU2F3-positive tumors^13^. While DLL3 is expressed in over 80% of SCLCs^2,13,14^, only a subset of patients with DLL3-positive SCLC derived clinical benefit from tarlatamab^2^, suggesting that DLL3 expression alone is insufficient for response and that a more comprehensive, unbiased interrogation may identify biomarkers more strongly correlated with clinical benefit.

To date, unbiased longitudinal analyses of response and resistance mechanisms in patients receiving tarlatamab have been lacking, owing both to its recent clinical approval and to the limited availability of serial tumor biopsies in this disease. Liquid biopsy approaches offer a powerful alternative for studying tumor evolution in SCLC. Recent advances in cell-free chromatin profiling have enabled genome-wide epigenomic interrogation of tumors using plasma^15–19^, providing access to transcriptional and regulatory states without the need for invasive tissue sampling. In particular, cell-free ChIP-sequencing (cfChIP-seq) has been shown to capture histone modification patterns associated with gene activation, enabling tumor subtyping and longitudinal monitoring of epigenomic evolution^15,16^. Recently, we developed a computational method to predict gene expression in cancer from cfChIP-seq data called APEX (Associating Plasma Epigenomic features with eXpression)^20^. APEX infers tumor gene expression from circulating chromatin with enhanced accuracy over existing methods^15,18,21–26^, offering a noninvasive window into dynamic transcriptional changes associated with resistance.

## RESULTS

### Liquid Biopsy Epigenomic Profiling Reveals the ASCL1 Subtype Predicts Response to Tarlatamab in SCLC

To identify genes associated with response to tarlatamab, plasma samples were prospectively collected from 46 patients receiving tarlatamab. The cohort comprised 40 patients (87.0%) with *de novo* SCLC, two (4.3%) poorly differentiated neuroendocrine carcinoma (NEC), two (4.3%) with SCLC transformation from *EGFR*-mutant lung adenocarcinoma, and two (4.3%) with metastatic carcinoid histology. In total 167 plasma samples were collected longitudinally during tarlatamab treatment and analyzed using our cell free epigenomic profiling assay^15^. Based on the first imaging evaluation, 18 patients derived “clinical benefit”, including 15 partial responses (PR) and 3 cases of stable disease (SD), whereas 22 derived “no clinical benefit” including 20 cases of progressive disease (PD) and 2 cases of oligoprogression (Oligo-PD) (see **METHODS**), which is consistent with response rates from clinical trials^2,4^. Response status could not be determined for 6 patients due to insufficient follow-up at the time of analysis (**Figure 1A**). As of January 2026, after a median follow-up of 2.1 months, the median progression-free-survival (PFS) for the cohort was 3.4 months (95% CI: 2.0 to not reached). The median duration of tarlatamab treatment was significantly longer among patients who derived clinical benefit (3.7 months; range, 1.3 to 15.7) compared to those who did not (1.9 months; range, 1.0 to 3.4) (p = 0.007, two-sided Wilcoxon rank-sum test) (**Figure 1B**). Baseline demographic and clinical characteristics are summarized in **Table 1**.

**Figure 1.**
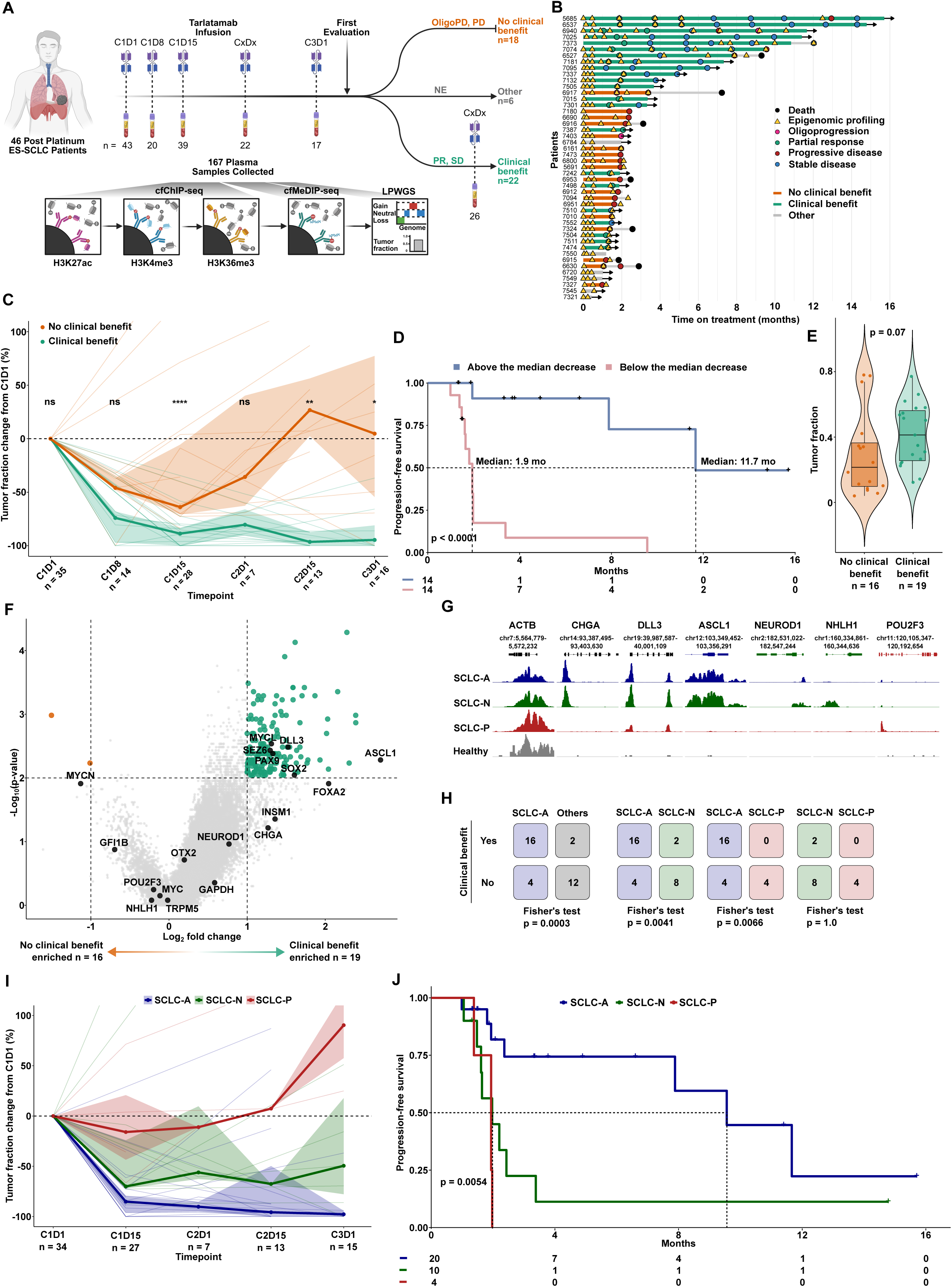
Liquid Biopsy Epigenomic Profiling Reveals the ASCL1 Subtype Predicts Response to Tarlatamab in SCLC. **(A)** Overview of the study cohort. Plasma samples were profiled using serial cell-free chromatin immunoprecipitation followed by sequencing (cfChIP-seq) targeting H3K27ac, H3K4me3, and H3K36me3; cell-free methylated DNA immunoprecipitation sequencing (cfMeDIP-seq); and low-pass whole-genome sequencing (LPWGS) (**Methods**). Number of patients and plasma samples collected is shown, along with best response at first radiographic assessment: partial response (PR) and stable disease (SD) together comprise “clinical benefit”, oligoprogression (Oligo-PD) and progressive disease (PD) together comprise “no clinical benefit”. Patients who received tarlatamab but had not yet undergone their first CT scan at the time of analysis are grouped under “Other”. (**B**) Swimmer plot showing the clinical course and treatment outcomes of the 46 patients treated with tarlatamab. (**C**) Line plot showing longitudinal tumor fraction dynamics stratified by clinical benefit (n=35 patients; 19 patients with clinical benefit and 16 without). Percent change in tumor fraction relative to C1D1 is shown across treatment timepoints (C1D1, C1D8, C1D15, C2D1, C2D15, C3D1) for clinical benefit and no clinical benefit. Thin lines represent individual patients. Solid lines indicate the group median at each timepoint, and shaded ribbons denote the interquartile range (IQR). The dashed horizontal line marks no change from baseline (0%). P values were calculated using two-sided Wilcoxon rank-sum tests at each timepoint; significance is indicated as ns (not significant), * (p < 0.05), ** (p < 0.01), and **** (p < 0.0001). (**D**) Kaplan-Meier curves showing progression-free survival (PFS) according to early tumor fraction decrease (n=28 patients with paired C1D1 and C1D15 samples). Patients were stratified by the median percent change in tumor fraction from C1D1 to C1D15 into above the median decrease (≥ 81.0%) or below the median decrease (< 81.0%). Median PFS was 11.7 months (95% CI, 7.9–NR) for ≥81.0% decrease and 1.9 months (95% CI, 1.6–NR) for <81.0%. PFS was calculated from treatment initiation to radiographic progression or censoring. Numbers at risk are shown. (**E**) Violin plot of baseline tumor fraction stratified by clinical benefit (n=35 patients ;19 patients w/ clinical benefit and 16 without clinical benefit). (**F**) Volcano plot of baseline APEX Log_2_TPM gene expression differences between patients with clinical benefit (n = 19 patients) and those without (n = 16 patients). The x-axis represents log₂ fold change (clinical benefit vs. no clinical benefit), and the y-axis represents –log₁₀(p-value). Dashed vertical lines indicate ±1 log₂ fold change, and the dashed horizontal line represents the significance threshold (p = 0.01). Gene labels highlight selected genes of interest in black. Genes enriched in patients with clinical benefit (p < 0.01, unadjusted; log₂ fold change > 1) are in green, whereas genes enriched in patients without clinical benefit (p < 0.01, unadjusted; log₂ fold change < −1) are in orange. (**G**) Genome browser view of H3K4me3 cfChIP-seq signal across representative patients and a healthy control. Shown are individual patients exemplifying each transcriptional subtype: SCLC-A (patient 5685), SCLC-N (patient 6916), and SCLC-P (patient 7324). Signal tracks are normalized to ACTB (housekeeping gene). (**H**) 2 × 2 contingency tables were constructed to compare patients with vs. without clinical benefit between subtypes. Corresponding Fisher’s exact test p-values are shown. Subtypes were grouped as SCLC-A (including Likely-A), SCLC-N (including Likely-N), and SCLC-P. (**I**) Longitudinal tumor fraction dynamics across early timepoints (C1D1, C1D15, C2D1, C2D15, C3D1) for all 34 patients in (**H**) grouped by subtype. The C1D8 timepoint was excluded because it was available for only one SCLC-N patient. Thin lines represent individual patients, colored by subtype. Thick lines indicate medians and shaded ribbons indicate IQR. The dashed horizontal line marks no change (0%). The decrease in tumor fraction from C1D1 to C1D15 was greatest in SCLC-A (median -85.1%), followed by SCLC-N (−70.0%) and SCLC-P (−15.9%). (**J**) Kaplan–Meier analysis of PFS stratified by subtype (SCLC-A, SCLC-N, SCLC-P, including Likely classifications). Time-to-event was defined as days from baseline to documented progressive disease. Median PFS is indicated by reference lines. Numbers at risk are shown below the plot. p-value is indicated.

**Table 1.**
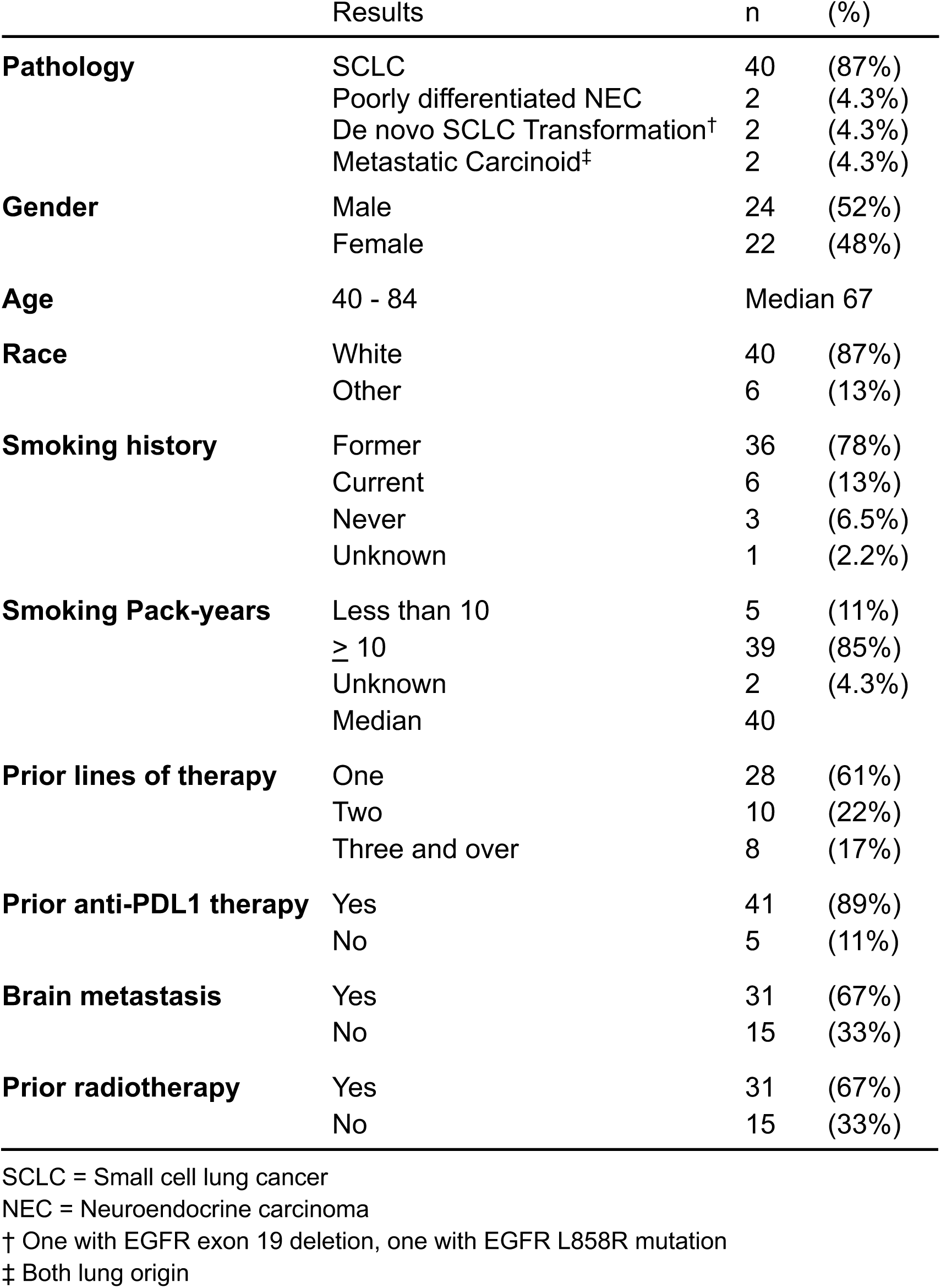
Summary of Baseline Demographic and Clinical Characteristics of Patient Cohort.

|  | Results | n | (%) |
| --- | --- | --- | --- |
| **Pathology** | SCLC | 40 | (87%) |
|  | Poorly differentiated NEC | 2 | (4.3%) |
|  | De novo SCLC Transformation<sup>†</sup> | 2 | (4.3%) |
|  | Metastatic Carcinoid<sup>‡</sup> | 2 | (4.3%) |
| **Gender** | Male | 24 | (52%) |
|  | Female | 22 | (48%) |
| **Age** | 40 - 84 | Median 67 |  |
| **Race** | White | 40 | (87%) |
|  | Other | 6 | (13%) |
| **Smoking history** | Former | 36 | (78%) |
|  | Current | 6 | (13%) |
|  | Never | 3 | (6.5%) |
|  | Unknown | 1 | (2.2%) |
| **Smoking Pack-years** | Less than 10 | 5 | (11%) |
|  | ≥ 10 | 39 | (85%) |
|  | Unknown | 2 | (4.3%) |
|  | Median | 40 |  |
| **Prior lines of therapy** | One | 28 | (61%) |
|  | Two | 10 | (22%) |
|  | Three and over | 8 | (17%) |
| **Prior anti-PDL1 therapy** | Yes | 41 | (89%) |
|  | No | 5 | (11%) |
| **Brain metastasis** | Yes | 31 | (67%) |
|  | No | 15 | (33%) |
| **Prior radiotherapy** | Yes | 31 | (67%) |
|  | No | 15 | (33%) |

SCLC = Small cell lung cancer
NEC = Neuroendocrine carcinoma
<sup>†</sup> One with EGFR exon 19 deletion, one with EGFR L858R mutation
<sup>‡</sup> Both lung origin

We next assessed whether early tumor fraction dynamics correlated with therapeutic efficacy of tarlatamab. Strikingly, we found that tumor fraction precipitously declined in patients with clinical benefit, exhibiting a near-complete clearance of ctDNA by C1D15 compared to more modest reductions in patients that did not (**Figure 1C, Figure S1A**). This two-week measurement translated to improved clinical outcomes: patients with a ctDNA decrease greater than the median value had a median PFS of 11.7 months vs. 1.9 months in patients with a less profound ctDNA response (**Figure 1D**). Baseline tumor fraction did not differ significantly between patients with clinical benefit vs. no clinical benefit (**Figure 1E**). These results demonstrate that early tumor fraction dynamics predict both treatment response and prognosis in SCLC.

To identify transcriptional correlates of tarlatamab response, we leveraged Associating Plasma Epigenomic features with eXpression (APEX), a computational method for inferring tumor RNA-seq level expression data from circulating cell-free chromatin (**Figure S1B-S1H**^20^). Inferred expression using APEX is referred to hereafter as “expression”. Differential expression analysis of baseline samples revealed 176 genes with enriched expression between patients with and without clinical benefit (**Figure 1F & Figure S1I**). Reassuringly, we observed significantly higher *DLL3* expression in patients deriving clinical benefit. Notably, many of the top genes enriched in patients deriving clinical benefit were components of the ASCL1 transcriptional program. These included ASCL1 itself as well as multiple direct ASCL1 target genes including the tarlatamab antigen DLL3, MYCL, SOX2, and FOXA2^9^, as well as PAX9, an ASCL1 lineage dependency^27^. We also observed enrichment of SEZ6, a promising antibody-drug conjugate target highly expressed in ASCL1-positive SCLC^28,29^. In contrast, no highly significant individual genes were strongly associated with lack of clinical benefit, although we observed modest enrichment of MYCN and GFI1B, genes previously associated with the POU2F3 SCLC subtype^30–32^. Collectively, these findings show that the ASCL1 transcriptional program is significantly correlated with response to tarlatamab.

Given that the ASCL1 transcriptional program is enriched in responders, we next asked whether transcriptionally defined SCLC molecular subtypes^13,14^ were associated with tarlatamab response. Using APEX-derived RNA-seq predictions and a random forest classifier trained on IMpower133 RNA-seq^33^ and subtype-specific gene signatures, we predicted baseline SCLC subtypes and evaluated their association with clinical outcomes in all SCLC patients with available baseline samples and known clinical outcomes (n=34 patients) (see **METHODS**). 20 (58.8%) were classified as the ASCL1 subtype (SCLC-A), 10 (29.4%) as the NEUROD1 subtype (SCLC-N), and 4 (11.8%) as the POU2F3 subtype (SCLC-P) **(Figure 1G**, **Figure S2A-S2B)**, mirroring previously reported subtype estimates^13^. To confirm these predictions, IHC for DLL3, ASCL1, NEUROD1, and POU2F3 was performed on tumor tissue from 6 available biopsies and was consistent with APEX-derived subtyping except for one sample (6917). This discrepancy may be due to tissue being obtained six months prior to the plasma sample as this patient received a line of therapy prior to tarlatamab, or to intra-tumor subtype heterogeneity being captured by APEX-derived RNA data but not by the tumor biopsy (**Figure S2C-S2D**).

We next examined whether clinical benefit differed by subtype: 16 of 20 (80.0%) SCLC-A patients had clinical benefit, compared with 2 of 10 (20.0%) SCLC-N and 0 of 4 (0%) SCLC-P patients. Notably, the SCLC-A subtype performed at least as well as, and trended toward improved predictive performance for clinical benefit relative to DLL3 expression alone (**Figure S2E**). Clinical benefit was significantly enriched in SCLC-A tumors compared with both SCLC-N and SCLC-P subtypes, whereas no significant difference was observed between SCLC-N and SCLC-P tumors **(Figure 1H).** Consistently, at C1D15, tumor fraction decreased most sharply in SCLC-A, followed by SCLC-N, and least in SCLC-P (**Figure 1I)**. Median PFS was longest in SCLC-A patients (9.7 months) compared with SCLC-N (2.0 months) and SCLC-P (2.0 months), underscoring the ASCL1 subtype as a robust predictor of response and longer-term clinical benefit to tarlatamab in SCLC (**Figure 1J**).

### An Immunocompetent ASCL1-Positive SCLC Mouse Model Exhibits Exquisite Sensitivity to Tarlatamab

We next sought to establish a preclinical immunocompetent mouse model to functionally study mechanisms of sensitivity and resistance to tarlatamab. Given the strong association between the ASCL1 subtype and tarlatamab sensitivity (**Figure 1**), we leveraged a previously described CRISPR-based genetically engineered *RPR2* SCLC mouse model (GEMM) of the ASCL1 subtype that is dependent on ASCL1^34–36^. From this GEMM model generated in a pure BL6J background, we previously derived transplantable syngeneic ASCL1-positive, DLL3-high tumor cell lines including the 1014P2 cell line^34–39^. 1014P2 cells highly express ASCL1 and DLL3 without expression of NEUROD1 or POU2F3 (**Figure 2A-2B**). Flow cytometric analysis confirmed DLL3 surface expression in 1014P2 cells, which was abrogated following CRISPR-mediated DLL3 inactivation in a polyclonal pool (**Figure 2B-2C**). We next assessed whether tarlatamab could directly bind mouse DLL3 using 1014P2 DLL3-isogenic [DLL3 wild-type (WT) vs. DLL3 knockout (KO)] cells generated using CRISPR/Cas9. Flow cytometric analysis demonstrated robust tarlatamab binding to DLL3 in DLL3 WT cells, which was markedly reduced following DLL3 KO, demonstrating that tarlatamab binds mouse cell surface DLL3 (**Figure 2D**). To determine whether tarlatamab could induce DLL3-dependent T-cell killing, we performed co-culture assays using 1014P2 cells and primary human T cells isolated from peripheral blood mononuclear cells (PBMCs) (**Figure 2E**). In the absence of T cells, tarlatamab had no effect on tumor cell viability (**Figure 2F**). In contrast, in the presence of T cells, tarlatamab induced dose-dependent tumor cell killing with half-maximal effective concentrations (EC₅₀) of approximately 1 nM, within the range of EC50’s reported in co-culture experiments with human SCLC cell lines^40^, with greater cytotoxicity observed at higher effector-to-target ratios (**Figure 2F**). T cell killing was nearly completely abolished in DLL3 knockout cells (**Figure 2G**), demonstrating that tarlatamab induces DLL3-dependent T cell-mediated killing of ASCL1-positive SCLC cells *in vitro*. Although tarlatamab recognizes mouse DLL3, it does not bind mouse CD3, necessitating the use of humanized CD3 mouse models to study its activity *in vivo*. We therefore employed BL6 mice in which the endogenous mouse CD3 locus was replaced with a human CD3 εδγ transgene. T cells isolated from the spleens of these mice bound tarlatamab as assessed by flow cytometry using an anti-human Fc antibody, whereas T cells from wild-type C57BL/6 mice showed no binding, confirming specificity for human CD3 (**Figure 2H**).

**Figure 2.**
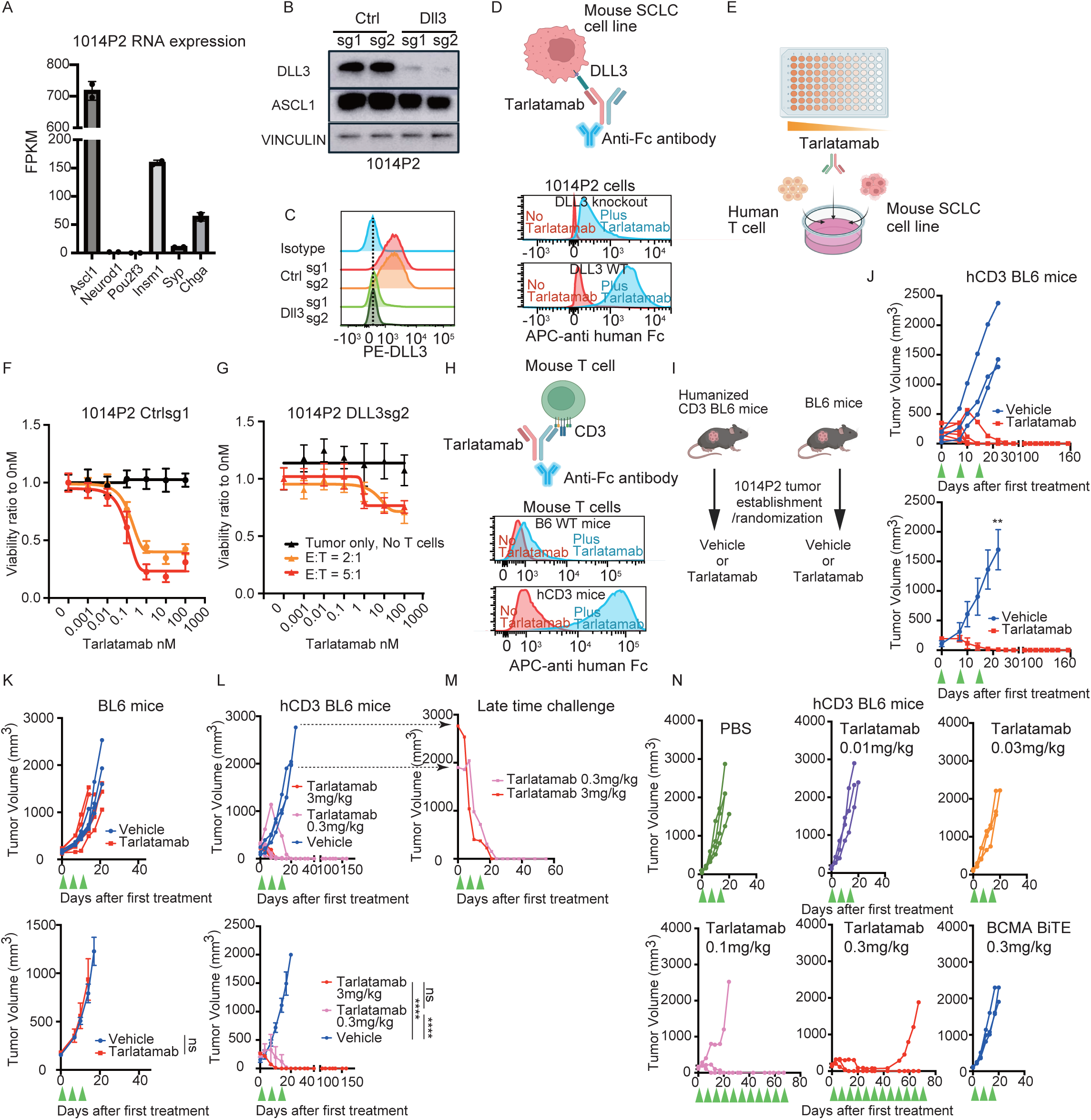
An Immunocompetent ASCL1-Positive SCLC Mouse Model Exhibits Exquisite Sensitivity to Tarlatamab. (**A**) Fragments per kilobase of transcript per million mapped reads (FPKM) values for representative lineage and neuroendocrine marker genes from RNA-sequencing of the 1014P2 mouse small-cell lung cancer cell line. (**B**) Immunoblot analysis of 1014P2 cells following CRISPR inactivation with either non-targeting control sgRNAs (Ctrl sg1 and sg2) or Dll3-targeting sgRNAs (Dll3 sg1 and sg2) for each indicated antibody. (**C**) Histogram from flow cytometric analysis for cell-surface DLL3 expression in 1014P2 transduced with either non-targeting control sgRNAs (Ctrl sg1 and sg2) or Dll3-targeting sgRNAs (Dll3 sg1 and sg2). y-axis is normalized to mode. (**D**) Schematic of the tarlatamab binding assay in a mouse SCLC cell line. 1014P2 cells transduced with either Dll3 sgRNA (DLL3 knockout) or control sgRNA (DLL3 WT) were incubated with (blue = Plus Tarlatamab) or without (red = No Tarlatamab) tarlatamab and subsequently stained with an APC-conjugated anti-Fc secondary antibody and analyzed by flow cytometry. Representative histograms of APC anti-Fc fluorescence are shown. y-axis is normalized to mode. (**E**) Schematic of *in vitro* cytotoxicity assay where healthy donor human T cells were co-cultured with mouse SCLC cell lines in the presence of tarlatamab at increasing concentrations.(**F-G**) Tumor cell viability following 48-hour co-culture in the absence of T cells (black) or in the presence of human T cells at effector-to-target (E:T) ratios of 2:1 (orange) or 5:1 (red) in 1014P2 control cells (**F**) or Dll3 knockout cells (**G**) at increasing concentrations of tarlatamab. Cell viability was quantified by flow cytometry, and values were normalized to the 0 nM tarlatamab condition. All conditions were performed in triplicates and mean +/- SD are shown. Representative result from three independent experiments. (**H**) Schematic of the tarlatamab binding assay in mouse T cells. T cells isolated from C57BL/6 wild-type (B6 WT) mice or humanized CD3 (hCD3) mice were incubated with (blue = Plus Tarlatamab) or without (red = No Tarlatamab) tarlatamab and subsequently stained with an APC-conjugated anti-Fc secondary antibody and analyzed by flow cytometry. Representative histograms of APC anti-Fc fluorescence are shown. y-axis is normalized to mode. (**I**) Schematic of *in vivo* tumor treatmentefficacy experiments in which humanized CD3 BL6 mice or C57BL/6 wild-type (BL6) mice were inoculated with 1014P2 cells subcutaneously and treated with either vehicle (PBS) or Tarlatamab. (**J**) Tumor growth of 1014P2 syngeneic mouse tumor subcutaneously injected into humanized CD3 BL6 mice (Biocytogen) treated with vehicle (n = 3, blue) or tarlatamab (3 mg/kg; n = 5, red). ** p < 0.01, Student’s unpaired two-tailed t test of tumor volume on day 18. (**K**) Tumor growth of 1014P2 syngeneic mouse tumor subcutaneously injected into in C57BL/6 wild-type (BL6) mice treated with vehicle (n = 4, blue) or tarlatamab (3 mg/kg; n = 6, red). ns = not significant, Two way ANOVA followed by Sidak’s multiple comparison test. (**L**) Tumor growth of 1014P2 syngeneic mouse tumor subcutaneously injected into humanized CD3 BL6 mice (genOway) treated with vehicle (n = 3, blue), tarlatamab (3 mg/kg; n = 5, red), or tarlatamab (0.3mg/kg; n= 4, pink). Ns = not significant, **** p < 0.0001, Two way ANOVA followed by Sidak’s multiple comparison test. For (**J-L**), Tarlatamab was administered intraperitoneally on days 0, 7, and 14 (green arrowhead). Top panel shows individual tumors and bottom panel shows mean +/- SEM. (**M**) Near endpoint 1014P2 subcutaneous syngeneic mouse tumor in humanized CD3 BL6 mice from Figure 2L (black broken arrow) were treated with tarlatamab 3mg/kg (red) or 0.3mg/kg (pink), intraperitoneally on days 0, 7, and 14 (green arrowhead) showing complete responses even in tumors with exceedingly large initial volumes close to 2000 mm^3^. (**N**) Tumor growth of 1014P2 syngeneic mouse cell line subcutaneously injected into humanized CD3 BL6 mice (genOway) treated with PBS (n = 4, green), tarlatamab 0.01 mg/kg (n = 3, purple), 0.03 mg/kg (n = 3, orange), 0.1 mg/kg (n = 3, pink), 0.3 mg/kg (n = 3, red), or a BCMA-targeting bispecific T cell engager Pavurutamab (BiTE; n = 3, blue) as a specificity control for human CD3 engagement. All treatments were administered intraperitoneally once weekly at the doses indicated starting on day 0 for the duration of the study (green arrowhead) and the tumor that developed acquired resistance at tarlatamab 0.3 mg/kg is the 1014P2-TR#2 model (see Figure 4).

We then used these humanized CD3 BL6 mice to evaluate *in vivo* antitumor efficacy. 1014P2 cells were implanted subcutaneously into either human CD3 transgenic mice (from Biocytogen) or wild-type BL6 as controls. Once tumors were established, mice were treated with tarlatamab (3 mg/kg)^40^ or vehicle intraperitoneally (IP), once weekly for three weeks (**Figure 2I**). In humanized transgenic CD3 BL6 mice, tarlatamab induced striking complete responses in all animals, with durable responses and no evidence of recurrence over several months (**Figure 2J**). In contrast, no antitumor activity was observed in BL6 wild-type mice (that express mouse CD3) (**Figure 2K**) demonstrating that efficacy was dependent on human CD3-mediated T cell engagement. To assess generalizability, we repeated these experiments in an independent humanized CD3 transgenic BL6 mouse strain (genOway), in which mouse CD3 is replaced by a pan-human CD3εδγ transgene. Tarlatamab (3 or 0.3 mg/kg) again induced complete and durable tumor regressions in all mice (**Figure 2L**). These doses approximate the clinically relevant human dosing range, with 3 mg/kg corresponding to higher clinical doses used in early trials and 0.3 mg/kg which is close to mouse equivalent for the 10 mg FDA-approved human dose. Strikingly, the antitumor activity of tarlatamab was sufficiently potent that even mice bearing large tumors nearing their endpoint (close to 2,000 mm³) at the time of treatment initiation achieved complete responses following delayed administration of tarlatamab (**Figure 2M**), underscoring the profound sensitivity of this ASCL1-positive, DLL3-high model. Dose de-escalation studies showed that tarlatamab induced complete responses at doses as low as 0.1 mg/kg, with no efficacy at ≤0.03 mg/kg, in humanized transgenic CD3 BL6 mice bearing established 1014P2 tumors (**Figure 2N**). As a specificity control, a bispecific T cell engager targeting B-cell maturation antigen (BCMA) was administered at 0.3 mg/kg and showed no antitumor activity (**Figure 2N**), confirming that tumor regression was not due to nonspecific T cell activation. Collectively, these data establish a syngeneic, immunocompetent mouse model of tarlatamab sensitivity, in which an ASCL1-positive, DLL3-high SCLC syngeneic model exhibits profound and durable responses to DLL3-directed T cell engagement. These findings are consistent with our plasma-derived predictors of response observed in human patients and provide a platform for mechanistic studies of response and resistance to tarlatamab in immunocompetent mice.

### Mechanisms of Acquired Resistance to Tarlatamab in Human SCLC: Selection for NEUROD1 Lineage Selection Associated with DLL3 Downregulation, Isolated Antigen Loss, and Immune Exhaustion

We next sought to identify potential mechanisms of acquired resistance to tarlatamab using APEX-derived transcriptional data, comparing plasma cfChIP-seq data at acquired resistance with baseline samples. To do this, we first identified patients in our cohort (**see Figure 1**) who initially responded to tarlatamab and subsequently developed acquired resistance.

### Selection for a NEUROD1-High, DLL3-Low State at Acquired Resistance

Patient 6257’s baseline tumor was classified as SCLC-A, characterized by strong *ASCL1* and *DLL3* signal and minimal *NEUROD1* expression. This patient initially demonstrated a deep molecular response with tumor fraction becoming undetectable between C1D1 and C1D15. This early molecular response was confirmed radiographically at C3D1 with a partial response by CT scan. However, despite stable disease on imaging, tumor fraction rose to 9.3% at C5D15, reaching 18.4% at C7D15 and 57.4% at C9D1, when radiographic progression was finally confirmed suggesting molecular progression preceded radiographic progression by at least 2 cycles (**Figure 3A**), highlighting the potential of liquid biopsy as a bellwether for radiographic progression.

**Figure 3.**
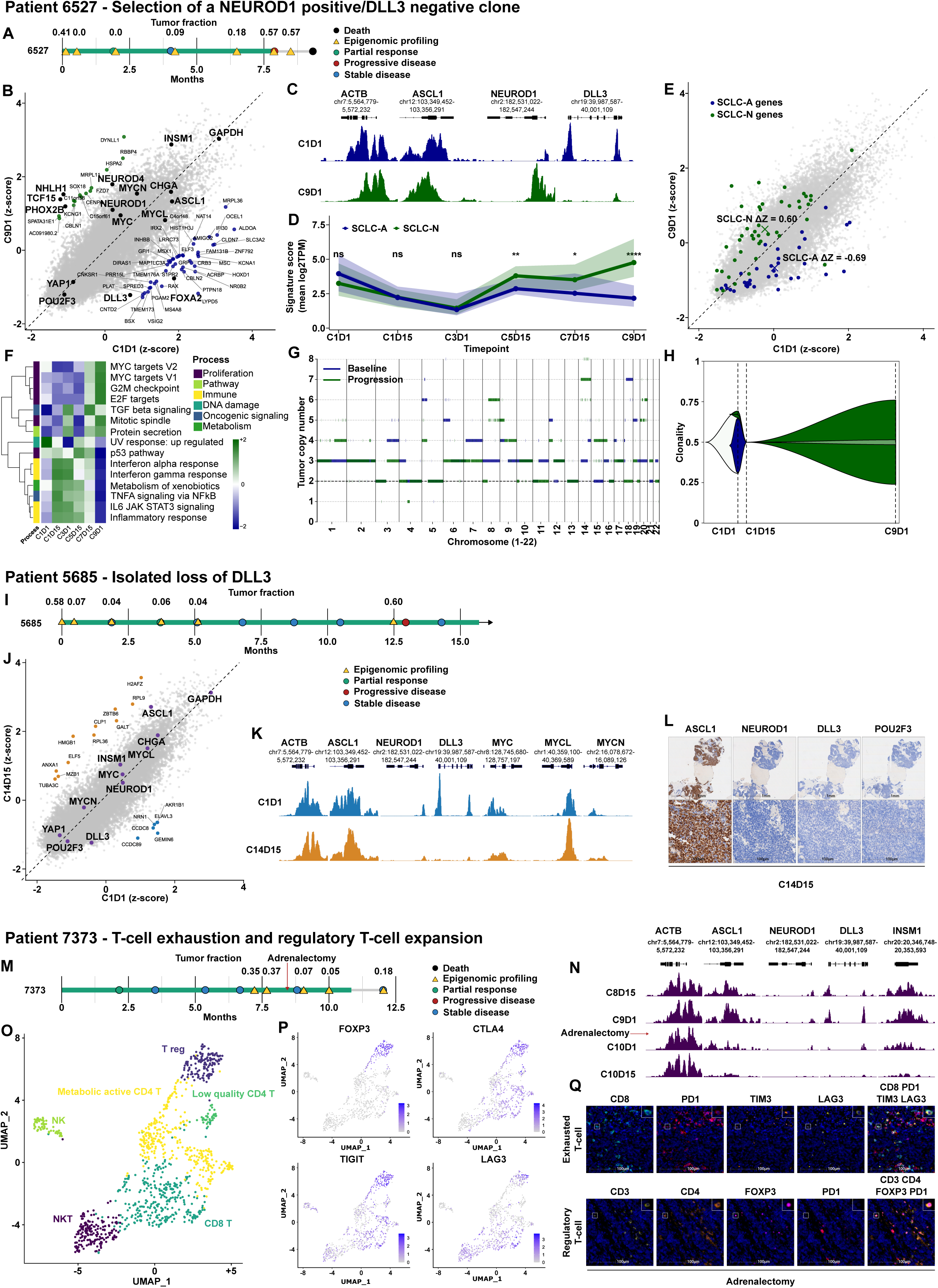
Mechanisms of Acquired Resistance to Tarlatamab in Human SCLC. **(A)**, **(I)**, **(M)** Swimmer plot showing the clinical course, treatment outcomes, and tumor fraction for patients 6527 (**A**), 5685 (**I**), and 7373 (**M**). (**B**) Scatterplot showing the Z-score of gene expression at C1D1 vs. at C9D1 for patient 6527. Each point represents a single gene. Genes (black) represent key genes of interest in SCLC. Genes classified as higher in C1D1 (dark blue) show a ≥2-unit increase in Z-score at C1D1 compared to C9D1, whereas genes higher in C9D1 (dark green) show a ≥2-unit increase in Z-score at C9D1 compared to C1D1. Other genes (grey) do not exhibit a substantial change between timepoints. The dashed diagonal line represents no change between timepoints. (**C**) Genome browser view of H3K4me3 cfChIP-seq signal at C1D1 (dark blue) and at C9D1 (dark green). (**D**) Line plot showing the dynamics of SCLC-N and SCLC-A transcriptional signatures (**Methods**) over serial timepoints for patient 6527. The thick colored lines represent the median expression per timepoint. Shaded ribbons indicate the interquartile range (25th–75th percentile). At each timepoint, the statistical difference between SCLC-N and SCLC-A signatures was assessed using a two-sided Wilcoxon rank-sum test, with significance indicated above the timepoints: * (p < 0.05), ** (p < 0.01), **** (p < 0.0001), **** (p < 0.0001); ns = not significant. SCLC-N is shown in dark green and SCLC-A in dark blue. (**E**) Scatter plot showing gene-level Z-scores at baseline vs progression for patient 6527. Each point represents one gene. Genes annotated as SCLC-A or SCLC-N according to predefined subtype gene lists^31^ are highlighted (SCLC-A, dark blue; SCLC-N, dark green), while all other genes are shown in grey. Subtype centroids (mean Z-score across genes within each subtype) are shown as crosses. The annotated ΔZ values indicate the mean change in Z-score (C9D1 − C1D1) for each subtype. (**F**) Heatmap showing the top 15 most variable MSigDB Hallmark pathways based on mean Z-scores across serial timepoints for patient 6527. Pathways are sorted by the absolute difference between C1D1 and C9D1. (**G**), Genome-wide copy number variation profiles for patient 6527 at C1D1 and at C9D1. (**H**) Hypothetical clonal evolution (“fish plot”) for patient 6527 across longitudinal sampling. Colored areas represent the estimated fraction of each clone at each timepoint (light blue, clone 1/ancestral; dark blue, clone 2; green, clone 3; light green, clone 4). Clonal relationships are depicted by nesting according to the specified parent structure (clone 1 is the founder clone; clones 2 and 3 arise from clone 1; clone 4 arises from clone 3). (**J**) Scatterplot showing the Z-score of gene expression at C1D1 vs. at C14D15 for patient 5685. Each point represents a single gene. Genes (black) represent key genes of interest in SCLC. Genes classified as higher in C1D1 (light blue) show a ≥2-unit increase in Z-score at C1D1 compared to C14D15, whereas genes higher in C14D15 (orange) show a ≥2-unit increase in Z-score at C14D15 compared to C1D1. Other genes (grey) do not exhibit a substantial change between timepoints. The dashed diagonal line represents no change between timepoints. (**K**) Genome browser view of H3K4me3 cfChIP-seq signal at C1D1 (light blue) and at C14D15 (orange). (**L**) Representative IHC for ASCL1, NEUROD1, POU2F3, and DLL3 from patient 5685 on C14D15 showing loss of DLL3 protein expression. (**N**) Genome browser view of cfChIP–seq H3K4me3 signal across representative loci for patient 7373. Of note, there was no baseline plasma sample available for this patient. (**O**) scRNAseq UMAP visualization of all CD3+ T cells (n=1040 CD3+ T cells) from adrenalectomy. NKT: Natural Killer T-cells, NK: Natural Killer, T reg: Regulatory T cells. (**P**) scRNAseq UMAP visualization of all CD3+ T cells showing expression of regulatory T cell markers and markers of T cell exhaustion: FOXP3, CTLA4, TIGIT, and LAG3. (**Q**) Representative multiplex immunofluorescence (mIF) staining of the adrenalectomy specimen of patient 7373.

To identify transcriptional changes associated with acquired resistance, we performed APEX-based differential expression analysis on plasma samples comparing acquired resistance (C9D1) with baseline (C1D1), revealing many genes that were differentially expressed at acquired resistance relative to baseline (**Figure 3B**). Notably, several NEUROD1 target genes that are highly expressed in the NEUROD1 subtype—including NEUROD4, TCF15, and NHLH1^9^—were significantly enriched at acquired resistance compared with baseline (**Figure 3B**). NEUROD1 itself also showed enrichment at acquired resistance (0.9 standard deviations relative to baseline), with IGV tracks for H3K4me3 revealing a prominently gained NEUROD1 promoter peak at progression (**Figure 3B-3C**). While ASCL1 expression was only modestly reduced, some of the most significantly and robustly downregulated genes at acquired resistance were those associated with the *ASCL1* subtype, including *DLL3* and *FOXA2*^9^, with loss of H3K4me3 peaks at the DLL3 locus at progression (**Figure 3B-3C**). These differences were unlikely related to tumor fraction which were comparable at baseline and progression (41.4% versus 57.4%) (**Figure 3A**). Consistent with these gene-level changes, lineage-centric analysis revealed coordinated transcriptional reprogramming at progression. SCLC-A–associated genes were globally downregulated, whereas SCLC-N–associated genes were concomitantly upregulated, indicating a shift toward a *NEUROD1*-enriched cell state at progression (**Figure 3D-3E**). Pathway analysis further demonstrated enrichment of MYC target programs, G2M checkpoint, and mitotic spindle signatures at progression, all characteristic of SCLC-N relative to SCLC-A^33^ (**Figure 3F**). Decreased DLL3 expression was not related to a genetic mutation in *DLL3* as whole genome sequencing (WGS) at progression did not reveal a *DLL3* mutation. A similar pattern of enriched NEUROD1 signatures at acquired resistance was observed from an independent patient (6940), who developed isolated progression after seven treatment cycles and exhibited enrichment of a NEUROD1 transcriptional program accompanied by near-complete loss of *DLL3* expression (**Figure S3A-S3B**).

To better characterize the tumor evolution on treatment, we performed copy-number profiling comparing C9D1 to C1D1 (see **METHODS**). Haplotype analysis of chromosome 15 at C1D1 supported a dominant clone with two B-alleles and no A-allele, and a minor subclone with an A-allele. At progression, the dominant clone contained four copies of chromosome 15 with three B-alleles and one A-allele. Thus, the dominant clone at progression contained an A-allele that was not present in the dominant clone at baseline. This observation suggests that the dominant clone at resistance did not evolve from the clone that predominated at treatment initiation. Rather, it may have been present before therapy and become dominant due to the selective pressure of treatment (**Figure 3G-3H & Figure S3C-S3E**).

Together, these findings support a transition from an ASCL1-driven, DLL3-high SCLC state to a NEUROD1-enriched transcriptional state associated with reduced DLL3 expression at acquired resistance. This is consistent with human SCLCs of the NEUROD1 subtype expressing lower levels of *DLL3* relative to *ASCL1*^13,33^, and is also in line with our findings (**see Figure 1**) showing that *ASCL1* is strongly correlated with response to tarlatamab, whereas *NEUROD1* is not.

### Isolated Loss of DLL3 Expression at Acquired Resistance

Patient 5685, who received tarlatamab for nearly one year, initially exhibited a precipitous decline in tumor fraction (from 57.8% at C1D1 to 6.8% at C1D15), followed by a durable response with sustained low tumor fraction (**Figure 1B**, **Figure 3I**). However, after 11 months of treatment, a liver metastasis was detected on CT imaging and the tumor fraction rebounded concordantly with this isolated progression (**Figure 3I**). In contrast to patient 6527, APEX-inferred transcriptome-wide analysis comparing the acquired resistance sample (C14D15) to the baseline sample (C1D1) revealed very few genes that were differentially expressed at acquired resistance, with no significant changes in expression of lineage transcription factors, including *ASCL1, NEUROD1*, and *POU2F3*. We noticed that DLL3 was relatively decreased (0.81 standard deviations relative to baseline) at acquired resistance relative to baseline with IGV traces revealing near-complete absence of *DLL3* at progression (**Figure 3J-3K**). These results were corroborated by IHC from a tissue biopsy at progression, where DLL3 was completely absent (**Figure 3L**). WGS did not identify acquired on-target mutations in DLL3 at progression. Haplotype-based BAF analysis from WGS revealed that, at C1D1, few arm-level copy-number alterations (CNAs) were explained by a single clone, whereas at C14D15, all arm-level CNAs were explained by a single clone (**Figure S3F**). This observation suggests that multiple clones contributed to the tumor burden at baseline, whereas a single clone predominated at acquired resistance. Together, these findings strongly suggest that isolated loss of *DLL3*, the tumor cell antigen targeted by tarlatamab, promotes acquired resistance to tarlatamab in this patient.

### Enrichment of Regulatory and Exhausted T Cells at Acquired Resistance

Patient 7373 initially achieved a partial response and developed isolated progression of an adrenal metastatic lesion after nine months of treatment with tarlatamab while other lesions continued to show a durable response. Given the isolated adrenal recurrence with overall continued systemic control of disease, the patient underwent an adrenalectomy to remove the adrenal metastasis and then continued on tarlatamab. Consistent with their clinical course described above, ctDNA dynamics tracked with adrenalectomy: tumor fraction decreased from 36.9% to 4.6% after adrenalectomy (**Figure 3M**). H3K4me3 cfChIP-seq prior to adrenalectomy showed high H3K4me3 promoter signal of *ASCL1, INSM1*, and *DLL3*, and lower *NEUROD1* signal. Following surgery, *ASCL1, INSM1*, and *DLL3* H3K4me3 signals from plasma cfChIP-seq were decreased, consistent with removal of the isolated site of progression contributing tumor-derived cell-free DNA to the plasma (**Figure 3N**).

Single-cell RNA-seq (scRNA-seq) was performed on the adrenalectomy surgical specimen. Consistent with our plasma cfChIP-seq findings, tumor cells uniformly expressed *ASCL1* and INSM1, heterogeneously expressed *DLL3*, and did not express *NEUROD1* (**Figure S3G-S3H**). Moreover, IHC confirmed protein expression of ASCL1 and DLL3 (**Figure S3I**). Strikingly, scRNA-seq of the adrenal metastasis showed that the majority of the cells were T cells (1040 cells, 57.4% of the total population) (**Figure S3G-S3H**). Notably, the number of T cells exceeded the number of tumor cells (431 cells, 23.8%), which is unusual for neuroendocrine SCLC, which typically is composed of sheets of tumor cells with a very low proportion of T cells^13,38,41^. Next, we subclassified the T-cell subsets from the adrenalectomy scRNA-seq data. Interestingly, this revealed a high proportion of FOXP3+ regulatory cells as well as T cells with an exhausted phenotype (CTLA4, TIGIT, and LAG3) (**Figure 3O-3P**). These findings were confirmed by multiplex IF (**Figure 3Q**). Given that this isolated site of adrenal progression showed an ASCL1⁺, DLL3⁺ tumor with marked enrichment of regulatory and exhausted T-cell phenotypes, these findings suggest that regulatory T cells and/or T-cell exhaustion are associated with acquired resistance to tarlatamab in this patient.

### Conserved Acquired Resistance to Tarlatamab in Mice Through NEUROD1-Driven DLL3 Loss and Immune Dysfunction

We next sought to develop mouse models of acquired resistance to tarlatamab using our immunocompetent system, enabling direct comparison with resistance mechanisms identified in human plasma and deeper mechanistic dissection. During extended *in vivo* treatment studies in which tarlatamab was administered weekly over several months (**Figure 4A, also see Figure 4N**), the majority of tumors exhibited durable responses; however, a single tumor treated with 0.3 mg/kg tarlatamab eventually progressed despite continued therapy, indicating the emergence of acquired resistance. Tumor tissue from this resistant lesion was harvested, and a cell line (1014P2-TR#2) was established for downstream analyses. Flow cytometric analysis demonstrated a marked loss of DLL3 surface expression in the resistant cell line relative to the parental 1014P2 cells (**Figure 4B, Figure S4A**). Immunoblotting further confirmed near complete loss of DLL3 protein levels (**Figure 4C**). Strikingly and consistent with our findings from human plasma (**see Figure 3**), this tumor and cell line at acquired resistance to tarlatamab exhibited robust upregulation of NEUROD1 with loss of DLL3, while maintaining ASCL1 expression (**Figure 4C-4D, Figure S4B**).

**Figure 4.**
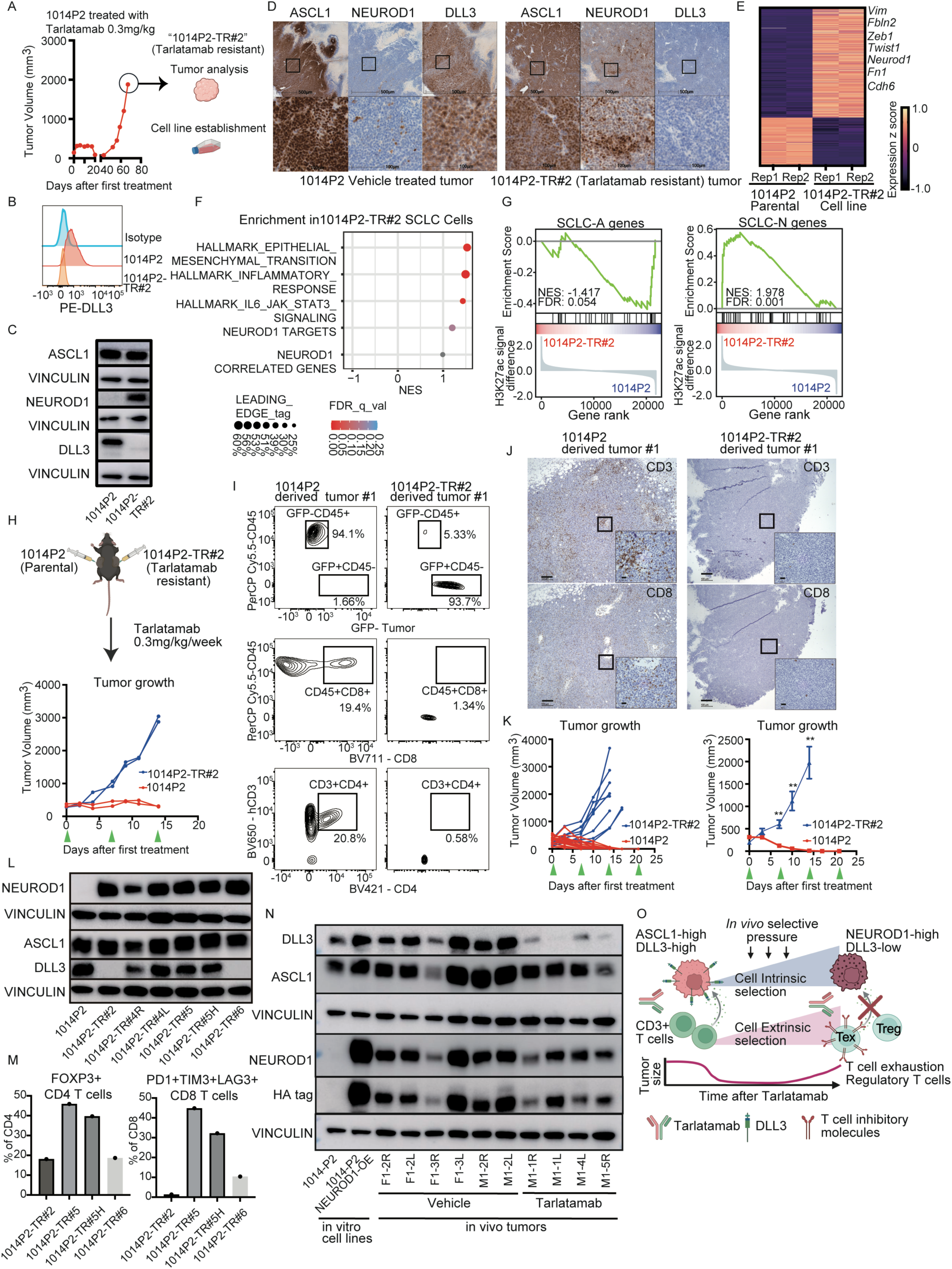
Conserved Acquired Resistance to Tarlatamab in Mice Through NEUROD1-Driven DLL3 Loss and Immune Dysfunction. (**A**) An acquired Tarlatamab-resistant tumor (1014P2-TR#2) was harvested for downstream tumor analyses, and a stable cell line was established for subsequent experiments. (**B**) Representative flow cytometric histogram for cell-surface DLL3 expression in the parental 1014P2 and 1014P2-TR cell line from three independant experiments. For (b), y-axis is normalized to mode. (**C**) Immunoblot analysis of parental 1014P2 cells and 1014P2-TR#2 cells with the antibodies indicated. (**D**) IHC staining for ASCL1, NEUROD1, and DLL3 in a vehicle-treated 1014P2 tumor (left) and the 1014P2-TR#2 tumor (right). Black box on top row figures indicate the area magnified on the bottom row. Original magnification 5x for top row and 40x for bottom row. (**E**) Heatmap of top differentially expressed genes (p value<0.05) from RNA-seq comparing 1014P2-TR#2 cells (2 biological replicates) vs. parental 1014P2 cells (2 biological replicates) (total 714 genes). Neurod1 and EMT genes are labeled. (**F**) Hallmarks gene set enrichment analysis of RNA-seq data from Figure 4F of top enriched gene sets in 1014P2-TR#2 cells vs. parental 1014P2 cells. NES = Normalized Enrichment Score. (**G**) Gene set enrichment plot of ASCL1 (left) and NEUROD1 (right) gene signatures (see **Method**) using H3K27ac ChIP-seq data comparing 1014P2-TR#2 vs. 1014P2. (**H**) *In vivo* tarlatamab competition experiments where 1014P2 and 1014P2-TR#2 cells were subcutaneously injected into opposite bilateral flanks of the same humanized CD3 BL6 mouse (one cell line per flank; n=2 independent mice, one genOway and one Biocytogen mouse) and treated with tarlatamab (0.3 mg/kg) by IP injection on days 0, 7, and 14 (green arrow head). (I) Tumors from Figure 4H were harvested at experimental endpoint (>2000 mm^3^) and analyzed by flow cytometry for GFP (for tumor cells), and CD45+, CD3+, CD4+, CD8+ for immune cell infiltration. (**J**) IHC for CD3 and CD8 of 1014P2 parental tumor and 1014P2-TR#2 tumor from experiment in Figure 4H. Black box indicates the area magnified on bottom right corner of each image. Black bar is 100 mm for weak magnification (original maginification 5x) and 25 mm for magnified image (original magnification 40x). (**K**) Parental 1014P2 and 1014P2-TR#2 cells were injected subcutaneously into individual humanized CD3 mice and treated with tarlatamab (0.3 mg/kg) administered weekly on days 0, 7, 14, and 21 (green arrow head). Left panel shows individual tumor growth and right panel shows mean +/- SEM. Two-way ANOVA followed by Sidak’s multiple-comparisons test was conducted ((**p < 0.01, n=16 tumors (eight mice) for parental 1014P2 and n=8 tumors (four mice) for 1014P2-TR#2)).(**L**) Immunoblot analysis for ASCL1, NEUROD1, and DLL3 from 1014P2 and 1014P2 tarlatamab resistant cell lines (TR#2, 4R, 4L, 5, 5H, and 6) established from tarlatamab resistant tumors. Details of each resistant tumor cell line are in the methods section. (**M**) Tumors from 1014P2 resistant tumors (TR#2, 5, 5H, and 6) were harvested and dissociated into single cell to assess FOXP3 expression on CD4+ T cells (left panel) and PD1/TIM3/LAG3 expression on CD8+ T cells (right panel). (**N**), Immunoblot analysis of parental 1014P2 cells, 1014P2 cells transduced with NEUROD1 overexpression (OE) vector, and mouse tumors made from 1014P2 NEUROD1 overexpression cell line treated with either vehicle or tarlatamab. Of note, the **Figure S5H** immunoblot contains the same 1014P2 NEUROD1 overexpression mouse tumors with additional controls. (**O**) Schematic of the proposed model in which ASCL1-driven, DLL3-high SCLCs are highly sensitive to tarlatamab. This response requires tumor cell expression of DLL3 driven by ASCL1 and functional CD3⁺ T cells. Tarlatamab exerts both tumor-intrinsic and tumor-extrinsic selective pressure to overcome its mechanism of action. Tumor cells undergo subtype selection toward NEUROD1, which promotes loss of DLL3 and can ultimately lead to acquired resistance. In parallel, tarlatamab promotes enrichment of regulatory and exhausted T cells, compromising CD3 effector function and thereby limiting therapeutic efficacy. Although both mechanisms can be selected under therapeutic pressure, one mechanism of acquired resistance ultimately predominates within a given tumor to sustain tumor growth in the presence of tarlatamab. Figure made using BioRender.

To define transcriptional changes associated with resistance, we performed RNA sequencing on: 1) the tarlatamab acquired resistant tumor versus vehicle-treated parental tumors; 2) the tarlatamab acquired resistant cell line versus parental 1014P2 cells. In both analyses, Neurod1 emerged as one of the most significantly upregulated genes (**Figure 4E-4F, Figure S4C-S4D**). In addition, multiple EMT-associated transcriptional regulators, including VIM, ZEB1, and TWIST1 (**Figure 4E-4F, Figure S4C-S4D**), were strongly induced, indicating acquisition of a mesenchymal-like state that is enriched in the NEUROD1 subtype of human SCLC^42^. Although changes in ASCL1 and NEUROD1 mRNA expression in tarlatamab-resistant samples relative to their tarlatamab-sensitive counterparts were concordant with the observed protein changes, DLL3 mRNA expression was surprisingly significantly higher, rather than lower, in the tarlatamab-resistant tumors and cell line compared with their tarlatamab-sensitive counterparts, suggesting a mechanism of post-transcriptional downregulation of DLL3 in mouse tumors (**Figure S4E-S4G**). To determine whether the observed mRNA expression changes were associated with epigenomic activation, and whether increased NEUROD1 expression promoted epigenomic changes consistent with a subtype switch toward the NEUROD1 SCLC subtype, we performed ChIP-seq for H3K4me3, H3K27ac, and H3K36me3 in 1014P2-TR#2 and 1014P2 cells. Consistent with our RNA-seq data, H3K4me3, H3K27ac, and H3K36me3 peaks at the NEUROD1 locus were increased in 1014P2-TR#2 cells relative to 1014P2 cells, with no change at the ASCL1 locus (**Figure S4H**). Gene set enrichment analysis of H3K27ac ChIP-seq data showed enrichment of NEUROD1 signature genes and depletion of ASCL1 signature genes in 1014P2-TR#2 cells relative to 1014P2 cells, supporting an ASCL1-to-NEUROD1 subtype switch at acquired resistance to tarlatamab (**Figure 4G**). Deep amplicon sequencing of the DLL3 gene revealed no genetic mutations in DLL3 itself in the tarlatamab resistant cell line. We next integrated these murine resistance-associated gene expression changes with APEX-derived inferred gene expression data from human plasma samples from patient 6527 obtained at acquired resistance found to have a gain in NEUROD1 signatures. This cross-species comparison identified a limited but highly concordant set of genes upregulated in across species, including genes associated with the NEUROD1 subtype and EMT (**Figure S4I**). These findings show that cell-state changes characterized by NEUROD1 and EMT programs and downregulation of DLL3 protein expression are associated with acquired resistance to tarlatamab in both humans and mice.

To determine whether this resistance mechanism was indeed tumor cell-intrinsic and maintained upon reimplantation *in vivo*, we performed tarlatamab *in vivo* competition experiments. Parental 1014P2 cells were injected into the left flank of human transgenic CD3 BL6 mice, while 1014P2-TR#2 acquired resistant cells were injected into the contralateral flank of the same animals (**Figure 4H**). Once tumors were established, mice were treated with tarlatamab at 0.3 mg/kg weekly. 1014P2 tumors exhibited rapid regression, whereas 1014P2-TR#2 tumors failed to respond, directly demonstrating acquired, tumor cell-intrinsic resistance to tarlatamab within the same host environment (**Figure 4H**). At tumor endpoint, flow cytometry and IHC analyses revealed marked immune infiltration in tarlatamab-sensitive tumors, with increased CD3⁺, CD4⁺, and CD8⁺ T cells (**Figure 4I-4J, Figure S4J-S4M**). In contrast, resistant tumors were largely devoid of immune infiltrates (**Figure 4I-4J, Figure S4J-S4M**). To further confirm acquired resistance *in vivo* in a larger number of independent mice, parental 1014P2 tumors and 1014P2-TR#2 resistant tumors were implanted into separate cohorts of humanized transgenic CD3 BL6 mice and treated with tarlatamab. Consistent with prior findings, 1014P2 tumors remained exquisitely sensitive, whereas 1014P2-TR#2 resistant tumors showed no response, establishing a true tumor cell-intrinsic mouse model of acquired resistance to tarlatamab (**Figure 4K**).

To more comprehensively interrogate mechanisms of acquired resistance, we continued weekly treatment with 0.3mg/kg tarlatamab for all parental tumors (8 mice, 16 tumors) in Figure 4K. All tumors underwent complete regression with no recurrence after 90 days, indicating that acquired resistance to tarlatamab in this highly sensitive ASCL1-positive 1014P2 model is rare (**Figure S5A**). To further probe mechanisms of resistance, we rechallenged seven independent mice that had previously achieved durable complete responses and remained tumor-free for more than 150 days after treatment. These mice were again injected bilaterally with 1014P2 cells (14 potential tumors). Ten of fourteen tumors grew following rechallenge and, once established, were treated with tarlatamab (**Figure S5B-S5C**). Six of ten tumors again achieved complete responses, whereas the remaining four developed acquired resistance (**Figure S5C)**. Resistant tumors were harvested and cell lines were established. Strikingly, all resistant tumors demonstrated induction of NEUROD1, whereas only one additional tumor showed DLL3 downregulation, similar to the 1014P2-TR#2 model (**Figure 4L, Figure S5D**). In contrast, tumors that retained DLL3 exhibited enrichment of FOXP3⁺ regulatory T cells and increased expression of exhaustion markers on CD8⁺ T cells, including TIM3 and LAG3 (**Figure 4M, Figure S5E**). The consistent induction of NEUROD1 across models, even in the absence of DLL3 loss, suggests that tarlatamab selects for both antigen escape and immune dysfunction, with one mechanism ultimately predominating within a given tumor under sustained therapeutic pressure.

Lastly, we asked whether forced NEUROD1 expression could promote DLL3 downregulation. 1014P2 cells were transduced to stably overexpress NEUROD1 (**Figure S5F**). NEUROD1 overexpression *in vitro* did not markedly alter DLL3 nor ASCL1 levels (**Figure S5F**). We next asked whether NEUROD1-overexpressing 1014P2 cells would select for DLL3 downregulation under tarlatamab pressure *in vivo*. Humanized CD3 BL6 mice were bilaterally injected subcutaneously with NEUROD1-overexpressing 1014P2 cells and, once tumors were established, randomized to receive tarlatamab or vehicle (three mice/six tumors for vehicle, four mice/seven tumors for tarlatamab). Two of seven NEUROD1-overexpressing tumors were inherently resistant to tarlatamab, whereas four initially achieved complete responses and one achieved a partial response **(Figure S5G).** Notably, two of the five responders rapidly developed acquired resistance. Immunoblot analysis of the four tumors that either failed to respond or developed early resistance revealed loss of DLL3 in all resistant tumors expressing exogenous NEUROD1 **(Figure 4N, Figure S5H)**, whereas none of the vehicle-treated NEUROD1-overexpressing tumors lost DLL3. Although this experiment was underpowered and does not establish that NEUROD1 overexpression alone is sufficient to cause resistance, these data indicate that enforced NEUROD1 expression under tarlatamab pressure promotes DLL3 downregulation. Together, our findings demonstrate convergent mechanisms of acquired resistance in both humans and mice, involving tumor-intrinsic adaptations in which NEUROD1 induction drives DLL3 downregulation, as well as tumor-extrinsic mechanism characterized by enrichment of exhausted and regulatory T-cell phenotypes. Our data suggest that both programs are progressively selected under sustained tarlatamab pressure, with the dominant mechanism ultimately manifesting as the observed mode of acquired resistance (**Figure 4O**).

## DISCUSSION

For decades, small cell lung cancer (SCLC) has lacked meaningful therapeutic breakthroughs. Although platinum-etoposide has a high initial response rate and combination with anti-PD-L1 immunotherapy has led to durable disease control in a small subset of patients^43–45^, the majority of patients experience recurrence within months. The DLL3 × CD3 bispecific T-cell engager tarlatamab represents one of the first clinically meaningful therapeutic breakthroughs in relapsed SCLC, producing objective responses in approximately 35-40% of patients and durable benefit in a substantial subset^2–4^. Understanding the biological determinants of response and resistance is essential for rational patient selection and the development of combinatorial strategies to enhance durability. A major obstacle in studying response and resistance is the limited availability of serial tumor biopsies, especially in SCLC. In this study, we overcame this challenge through prospective longitudinal plasma collection coupled with genome-wide cfChIP-seq profiling integrated with APEX to infer tumor gene expression at multiple timepoints, including acquired resistance. In parallel, we developed the first immunocompetent, ASCL1-high, DLL3-high syngeneic mouse model capable of modeling both exquisite sensitivity and acquired resistance to tarlatamab. Across species, our findings converge on a central and unifying principle: transcription factor subtype governs both primary response and acquired resistance to DLL3-directed T-cell engagement.

Our plasma-based transcriptome-wide analysis demonstrates that SCLC subtype is a major determinant of primary response to tarlatamab. The ASCL1 subtype emerged as one of the most robust predictors of benefit, with 80% of ASCL1-subtype tumors achieving clinical benefit, followed by NEUROD1-subtype tumors (20%), whereas all four POU2F3-subtype tumors were uniformly resistant. These findings are biologically grounded. DLL3 is a direct transcriptional target of ASCL1^9,10^ and is most highly expressed in ASCL1-dominant tumors, expressed at lower levels in NEUROD1 tumors, and minimally expressed in POU2F3 tumors^13^. Although DLL3 is the direct molecular target of tarlatamab and correlates with response in our study and others^2,4,46^, our unbiased genome-wide analysis suggests that response is more broadly linked to the ASCL1 transcriptional program (i.e., the ASCL1 subtype), which includes DLL3 as a direct ASCL1 target gene. Notably, human neuroendocrine SCLCs frequently exhibit intratumoral heterogeneity between ASCL1 and NEUROD1 expression, either within the same tumor cells or across distinct subpopulations^47–49^. In this context, our data suggest that ASCL1-dominant tumors are most sensitive to tarlatamab, while tumors exhibiting intratumoral heterogeneity with both ASCL1 and NEUROD1 expression—even when DLL3 is present—will be less responsive or more prone to develop acquired resistance. This model is consistent with our observation that NEUROD1 dominant tumors show lower response rates as well as increased acquired resistance to tarlatamab. This may also explain why multiple ASCL1-regulated genes, including DLL3, MYCL, FOXA2, and SOX2, were enriched in responders. Because plasma profiling integrates signals across multiple tumor sites, it may more accurately capture subtype heterogeneity than single-lesion biopsies. Clinically, our findings suggest that subtype-enriched enrollment strategies should be further explored. Notably, although the sample size was limited, we did not observe responses in POU2F3-dominant tumors, suggesting that this subtype may represent a subset of SCLC that is less responsive to tarlatamab and for which alternative therapeutic strategies should be prioritized. Future studies in larger cohorts will be needed to evaluate transcription factor subtype and DLL3 expression as predictive biomarkers of response, as well as the role of intratumoral subtype heterogeneity in primary resistance.

Tarlatamab’s mechanism of action requires both DLL3 antigen expression on tumor cells and functional T-cell engagement. Accordingly, we observed that tumors develop acquired resistance to tarlatamab through either decreased DLL3 expression or mechanisms that impair T-cell function. The most striking finding is that resistance to tarlatamab frequently emerges through NEUROD1 lineage selection across species. Under therapeutic pressure, tumors select for a NEUROD1-dominant state accompanied by DLL3 downregulation. This may occur through clonal selection, supported by extensive copy-number alterations found at progression in a human patient, or through lineage plasticity, which remains biologically plausible. Although we observed isolated DLL3 loss without NEUROD1 gain in one human SCLC that developed acquired resistance, we more commonly observed NEUROD1 gain associated with DLL3 loss across both human and mouse SCLCs. We note that the numbers of patients with acquired resistance are limited and this will continue to be an important area of investigation.

We find that NEUROD1 is unlikely to be purely correlative. Under selective pressure from tarlatamab, enforced NEUROD1 expression promoted DLL3 downregulation *in vivo*. Our data suggest that this effect depends on therapeutic pressure exerted by tarlatamab and is therefore likely indirect, as NEUROD1 overexpression does not downregulate DLL3 *in vitro* and NEUROD1 is canonically a transcriptional activator. The mechanisms by which NEUROD1 promotes DLL3 downregulation under tarlatamab selective pressure remain unclear and will be an important focus of future studies. These findings demonstrate that a targeted therapy directed against a lineage-dependent antigen can exert selective pressure against the lineage transcription factor state that drives that antigen. In this case, tarlatamab targets DLL3, an ASCL1-regulated gene^9,10^, and resistance emerges through selection against the ASCL1-dominant state in favor of NEUROD1. Most neuroendocrine SCLC tumors are dominated by either ASCL1 or NEUROD1 but retain minor subpopulations expressing the alternate transcription factor^47–49^. Genetic alterations such as MYC activation^48,50^ and KDM6A loss^51^ have been shown to promote ASCL1-to-NEUROD1 switching, raising the possibility that tumors harboring these alterations may be more permissive to lineage transitions under selective pressure from tarlatamab and therefore more prone to develop resistance through subtype selection. Even rare NEUROD1-positive subclones may be sufficient to seed resistance under sustained therapeutic pressure. Consistent with this model, several patients with NEUROD1-subtype tumors who developed progressive disease at the first radiographic evaluation exhibited early decreases in tumor fraction followed by rapid rebound, suggesting initial elimination of ASCL1-dominant clones with subsequent outgrowth of resistant populations enriched for NEUROD1. Although we did not observe ASCL1-to-POU2F3 transitions in this cohort, preclinical mouse models have demonstrated that such transitions can occur, particularly in the context of ASCL1 and PTEN loss^52^. Given the intrinsic resistance and low DLL3 expression observed in POU2F3 tumors, selection toward a POU2F3 phenotype represents a plausible additional mechanism of acquired resistance.

Consistent with tarlatamab’s mechanism of action, we also observed tumor-extrinsic resistance associated with enrichment of regulatory T cells and exhausted CD8⁺ T cells in both human and mouse models. Immune dysfunction occurred despite preserved DLL3 expression, indicating that immune failure alone can drive resistance. Similar associations between elevated regulatory T cell levels and inferior outcomes have been reported in patients treated with the BCMA-directed bispecific teclistamab^53^, and T-cell dysfunction has also been observed in a recent tarlatamab study^46^. Interestingly, in our mouse model, this resistance mechanism was only observed upon tumor rechallenge in mice that had previously achieved a complete response to tarlatamab and was not observed in naïve mice developing resistance for the first time. Whether this reflects loss of immune control late after a prolonged complete response remains an open question.

Several resistant tumors displayed features of both programs: NEUROD1 induction with immune exhaustion without complete DLL3 downregulation. These observations suggest that tarlatamab exerts parallel selective pressure on ASCL1-driven, DLL3-expressing tumors, promoting both NEUROD1-associated DLL3 downregulation and T-cell dysfunction, with one mechanism ultimately predominating within a given tumor. Whether NEUROD1 induction directly influences immune remodeling remains unknown, but these two dominant resistance mechanisms underscore the multifactorial nature of resistance to this dual-targeting therapeutic. This work suggests future strategies aimed at constraining the ASCL1 lineage, blocking NEUROD1-driven DLL3 downregulation, and restoring T-cell functionality to overcome tarlatamab resistance should be explored.

We find that early tumor fraction kinetics predicted durable clinical benefit from tarlatamab as early as 1 week after the first full dose of tarlatamab (C1D15) and well before standard radiographic evaluation, indicating that durable therapeutic response is largely determined during the initial two weeks of therapy. The magnitude of tumor fraction reduction at C1D15 closely mirrors reductions reported at later timepoints in a recently reported independent tarlatamab-treated cohort^54^, with our data demonstrating that response trajectories are established early and remain stable thereafter. Our findings are in line with prior observations in first-line extensive stage SCLC treated with chemo-immunotherapy^55^ and refine this paradigm to demonstrate that clinical benefit discrimination can occur substantially earlier, shifting the predictive window closer to treatment initiation. These data highlight the translational potential of ctDNA fraction as an early, clinically actionable biomarker that could enable discontinuation of ineffective therapy and reduce unnecessary toxicity or perhaps inform escalation treatment strategies early on for patients that will not have clinical benefit from tarlatamab alone.

Over the past several years, molecular subclassification has revealed that SCLC comprises distinct transcription factor-defined molecular subtypes, providing a framework for more personalized targeted therapeutic approaches for patients with SCLC^13,14^. Several subsequent studies have identified subtype-specific vulnerabilities^13,30,31,56–58^. Other work has focused on subtype plasticity in preclinical genetic models, suggesting that subtype selection could serve as a mechanism of resistance to therapies targeting subtype-specific vulnerabilities^48,51,52^; however, direct evidence of this phenomenon in the context of a clinically effective therapy has been lacking. Our findings demonstrate that subtype identity is not only predictive of response but can also be dynamically selected under therapeutic pressure as a mechanism of escape. To our knowledge, this represents the first example of a highly effective therapy used to treat patients with SCLC that both depends on and reshapes transcription factor lineage state as a mechanism of resistance. Together, these results position SCLC subtype as a dynamic evolutionary axis that both determines therapeutic vulnerability and is reshaped under selective pressure from subtype-directed therapy. More broadly, this work provides a framework for understanding how therapies targeting lineage-associated cell-surface antigens can drive cancer evolution, with implications for response and resistance to cell-surface-targeted therapies across cancer types.

## METHODS

### Human Subjects

#### Ethical considerations

All patients included in this study signed informed consent to Dana-Farber Cancer Institute (DFCI) IRB protocol #02-180 to collect blood and/or tissue for this study. DFCI IRB protocol # 02-180 titled OPTIMAL, is a biobanking study that collects specimen and clinical data for patients with thoracic malignancies across our clinics and satellite sites. Per DF/HCC policy all patients included in research must provide full informed consent in order to collect specimen and data. Consent was obtained both on paper and through electronic consent which was 21 CFR Part 11 compliant.

#### Clinical cohort

##### Inclusion criteria

Patients with small cell lung cancer (SCLC; *de novo* or transformed), poorly differentiated neuroendocrine carcinoma (LCNEC), or metastatic carcinoid histology who were treated with tarlatamab at Dana-Farber Cancer Institute were prospectively enrolled between September 2024 and December 2025. In total, this included 46 patients comprised of 40 patients (87.0%) with *de novo* SCLC, two (4.3%) with poorly differentiated neuroendocrine carcinoma, two (4.3%) with SCLC transformation from *EGFR*-mutant lung adenocarcinoma, and two (4.3%) with carcinoid histology. The median age at diagnosis was 67 years (range, 40-84). Forty-two patients were current or former smokers (91%), and all had received at least one prior line of therapy. Thirty-one patients (67%) had brain metastases at baseline, and 31 (67%) had received prior radiotherapy. Baseline demographic and clinical characteristics are summarized in **Table 1**.

##### Patient Classification Based on Clinical Benefit

Patients were classified as having clinical benefit if the first CT scan demonstrated partial response, or stable disease. Absence of clinical benefit was defined as progressive or oligo-progressive disease. Acquired resistance was defined as patients who initially experienced clinical benefit but subsequently developed progressive disease. Patients who received tarlatamab but had not yet undergone their first CT scan at the time of analysis were categorized as “Other”.

##### Definition of the Evaluable Population

A total of 46 patients with SCLC treated with tarlatamab were included in the study cohort. Among these, 43 patients had a plasma sample available at baseline, and clinical outcome data were available for 40 patients. 37 patients had both baseline plasma samples and clinical outcome data, allowing evaluation of baseline biomarkers in relation to treatment response.

For baseline tumor fraction analyses, 35 patients had evaluable plasma samples, including 19 with clinical benefit and 16 without clinical benefit. For longitudinal tumor fraction analyses, 31 patients had an additional plasma sample collected at cycle 1 day 15 (C1D15), including 16 with clinical benefit and 15 without clinical benefit. Among these, 28 patients had both baseline and C1D15 samples available, enabling assessment of tumor fraction kinetics during treatment (16 with clinical benefit and 12 without clinical benefit). SCLC molecular subtype classification analysis included 34 patients.

Two patients were excluded from the tumor fraction analysis: one patient (7242) had undetectable tumor fraction in all plasma samples, and another patient (7181) was a statistical outlier. An additional patient was excluded from the SCLC subtype analysis because pathology showed poorly differentiated neuroendocrine carcinoma and not SCLC (7505).

##### Sample Collection and Processing

Peripheral blood (3–10 mL) was drawn into K2-EDTA tubes (BD Biosciences, Cat. #366643) and processed within 4 hours of collection. Initial centrifugation was performed at 1,500 × g for 10 minutes at 4 °C to separate the plasma. The supernatant was carefully transferred to a new conical tube and subjected to a second centrifugation at 1,500 × g for 10 minutes at 4 °C to remove any residual cellular debris. Protease inhibitor (Roche, Cat. #11873580001) was added to the clarified plasma. Processed plasma was aliquoted, flash-frozen in liquid nitrogen, and stored at −80 °C until further analysis.

#### Liquid biopsy epigenomic profiling

##### Cell-free chromatin immunoprecipitation and high-throughput sequencing assay (cfChIP-seq)

cfChIP-seq was performed as described^15^. In brief, 5 μg of antibody were covalently conjugated to 1 mg of Dynabeads M-270 Epoxy (Invitrogen, Cat. #14302D) for a minimum of 16 hours at 37°C with rotation, using buffer C1 and C2 from the Dynabeads Antibody Coupling Kit (Invitrogen, Cat. #14311D) according to the manufacturer’s protocol. After coupling, the beads were pre-cleared by incubating with 0.1% BSA in PBS for 5 minutes at 4°C with rotation. The following antibodies were used, all at a dilution of 1 μg per 900 μl: H3K4me3 (Thermo Fisher Scientific, Cat. #711958), H3K27ac (Abcam, Cat. #ab4729), and H3K36me3 (Diagenode, Cat. #C15410192). Thawed plasma was centrifuged at 3,000g for 15 min at 4 °C after adding protease inhibitor (Roche, Cat. #11873580001). The plasma was subjected to antibody-coupled magnetic beads overnight with rotation at 4 °C. The reclaimed magnetic beads were washed with 1 mL of each washing buffer twice. Three washing buffers were used in the following order: low-salt washing buffer (0.1% SDS, 1% Triton X-100, 2 mM EDTA, 150 mM NaCl, 20 mM Tris-HCl, pH 7.5), high-salt washing buffer (0.1% SDS, 1% Triton X-100, 2 mM EDTA, 500 mM NaCl, 20 mM Tris-HCl, pH 7.5) and LiCl washing buffer (250 mM LiCl, 1% NP-40, 1% Na deoxycholate, 1 mM EDTA, 10 mM Tris-HCl, pH 7.5). Samples were incubated at 4 °C for 1 minute for each wash. Subsequently, the beads were rinsed with TE buffer (Thermo Fisher Scientific, Cat. #BP2473500) and resuspended and incubated in 100 μl of DNA extraction buffer containing 0.1 M NaHCO3, 1% SDS and 0.6LJmgLJml−1 Proteinase K (Qiagen, Cat. #19131) and 0.4LJmgLJml−1 RNaseA (Thermo Fisher Scientific, Cat. #12091021) for 10 min at 37 °C, for 1 h at 50 °C and for 90 min at 65 °C. DNA was purified according to the manufacturer protocol using the ChIP DNA Clean & Concentrator kit (Zymo Research, Cat. #D5205). cfChIP-seq libraries were prepared with ThruPLEX® DNA-Seq Kit (Takara Bio, Cat. #R400675) following the manufacturer’s instructions. After library amplification, the DNA was purified by AMPure XP (Beckman Coulter, A63880). The size distribution of the purified libraries was examined using Agilent 2100 Bioanalyzer with a high-sensitivity DNA Chip (Agilent, Cat. #5067-4626). The library was submitted for 150-bp paired-end sequencing on an Illumina NovaSeq 6000 system (Novogene).

##### Cell-free methylated DNA immunoprecipitation and high-throughput sequencing assay (cfMeDIP-seq)

cfMeDIP-seq was performed using a previously published method^59^. cfDNA libraries were prepared using the KAPA HyperPrep Kit (KAPA Biosystems) according to the manufacturer’s protocol. Briefly, cfDNA underwent end-repair, A-tailing, and ligation to NEBNext adaptors (NEBNext Multiplex Oligos for Illumina kit, New England BioLabs). Libraries were digested with USER enzyme (New England BioLabs) and purified using AMPure XP beads (Beckman Coulter). To standardize input and assess immunoprecipitation efficiency, λ DNA (a mixture of unmethylated and in vitro methylated amplicons) was added to each library to achieve a total of 100 ng DNA. Additionally, 0.3 ng of methylated and unmethylated Arabidopsis thaliana DNA (Diagenode) was spiked in as a quality control. Prior to immunoprecipitation, DNA was heat-denatured at 95°C for 10 minutes and snap-cooled on ice for 10 minutes. Libraries were partitioned into 10% input control (7.5 µL) and 90% (75 µL) for immunoprecipitation, which was performed using the MagMeDIP Kit (Diagenode) following the manufacturer’s protocol. Samples were purified using the iPure Kit v2 (Diagenode) and eluted in 50 µL Buffer C. The success of immunoprecipitation was confirmed by qPCR, measuring recovery of spiked-in methylated and unmethylated Arabidopsis thaliana DNA. Libraries failing quality control thresholds (<1% recovery of unmethylated DNA and >99% recovery of methylated DNA) were excluded. Final libraries were assessed for quality using a TapeStation system (Agilent Technologies) and sequenced on an Illumina NovaSeq 6000 system with 150-bp paired-end sequencing (Novogene).

##### cfDNA extraction and low-pass whole genome sequencing (LP-WGS)

Following cfChIP-seq, cfDNA was extracted from 1 mL of plasma using the QIAamp Circulating Nucleic Acid Kit (QIAGEN, Cat. #55114), according to the manufacturer’s instructions. cfDNA concentration was quantified using the Qubit fluorometer (Thermo Fisher Scientific), and samples were stored at −80°C until further analysis. LP-WGS was performed on all cfDNA samples to evaluate genome-wide copy number alterations. Sequencing libraries were prepared and sequenced using 150 bp paired-end reads on an Illumina NovaSeq 6000 platform (Novogene). Copy number profiles and tumor fraction estimates were generated using the ichorCNA algorithm^60^.

##### Whole Genome Sequencing (WGS)

WGS was performed on selected samples to investigate potential acquired resistance mechanisms and to characterize clonal heterogeneity through haplotype-based BAF analysis. Up to 200 ng of cfDNA was used for library preparation with the ThruPLEX® Tag-Seq FLEX Library Preparation Kit (Takara Bio, Cat. #R400735), according to the manufacturer’s instructions. Following library amplification, DNA was purified using AMPure XP beads (Beckman Coulter, Cat. #A63880). Library size distribution was assessed on an Agilent 2100 Bioanalyzer using a High Sensitivity DNA Kit (Agilent, Cat. #5067-4626). Libraries were sequenced using 150-bp paired-end reads on an Illumina NovaSeq 6000 platform (Novogene) targeting a 40x coverage. Depth was calculated from BAM files using samtools depth with all genomic positions retained, no maximum depth cap, and reads spanning deletions included. Mean depth was defined as the average per-base coverage across positions.

#### Sequence data processing

##### cfChIP-seq

Raw sequencing data were demultiplexed using bcl2fastq2 (RRID:SCR_015058; v2.19) to generate sample-specific FASTQ files based on unique barcode indices. Downstream processing was performed using a reproducible Nextflow-based workflow (SNAPIE v1.5.1) optimized for cfChIP-seq analysis (https://github.com/prc992/SNAPIE)^61^. Paired-end reads were adapter-trimmed with Trim Galore (v0.6.10)^62^ (trim_method = TRIM_GALORE), while quality trimming was disabled (--quality 0) to preserve native fragment length distributions and end-motif sequences.

Trimmed reads were aligned to the hg19 reference genome (GCA_000001405.1) using Burrows–Wheeler Aligner (v0.7.18)^63^ with extended insert size parameters (–I 250,200,2000,10) to minimize incorrect classification of cfDNA fragments as improperly paired. Alignments were coordinate-sorted and filtered to retain uniquely mapped reads using samtools (v1.15.1). PCR and optical duplicates were removed with Picard MarkDuplicates (v2.27.4; RRID:SCR_006525). Additional filtering with samtools (–f 3 –F 3844 –q 30) excluded unmapped, secondary, supplementary, QC-failed, and duplicated reads, retaining only properly paired, high-confidence reads (mapping quality >30).

Filtered BAM files were converted to BEDPE format using BEDtools (v2.30.0)^64^ bamtobed with the –bedpe option, retaining fragments with non-negative insert sizes. Fragments overlapping regions in the ENCODE hg19 blacklist (ENCFF001TDO) were removed. Resulting fragment files were imported into R as GRanges objects (GenomicRanges v1.56.2)^65^, and fragments shorter than 20 bp or longer than 500 bp were excluded from downstream analyses. Coverage tracks were generated from aligned BAM files using bamCoverage (deepTools v3.5.6) with 10-bp binning and RPKM normalization, and exported in bigWig format.

##### cfMeDIP-seq and LP-WGS

cfMeDIP-seq and LP-WGS reads were processed following a standardized pipeline. Adapter sequences were trimmed using fastp (v0.23.2). Reads were aligned to the hg19 reference genome using BWA (v0.7.19) with insert size modeling disabled and the same alignment parameters as described above. Read pairs were retained if they were properly paired and had a mapping quality >5. Filtering was performed using samtools (v1.22.1). Duplicate reads were removed using Picard MarkDuplicates (v2.24.1).

##### Whole genome sequencing

WGS data were processed using a similar pipeline with modified parameters to prioritize sequence accuracy for haplotype-based BAF analysis. Adapter trimming and default length and quality filtering were performed using fastp. Reads were aligned to the hg19 reference genome using BWA as described above. A higher mapping quality threshold (≥20) was applied for downstream analyses. Duplicate reads were removed using Picard MarkDuplicates; for libraries containing unique molecular identifiers (UMIs), reads were considered duplicates only if they mapped to the same genomic coordinates and shared the same UMI.

#### Associating Plasma Epigenomics with eXpression (APEX)

APEX (Associating Plasma Epigenomic features with eXpression) is a machine learning framework that noninvasively infers genome-wide tumor gene expression from circulating cell-free chromatin^20^. The method integrates promoter- and gene body–associated histone modification coverage with fragmentomic features to model transcriptional activity at the gene level. For each gene (∼18,000 protein-coding genes), promoter regions (±3 kb around the transcription start site [TSS]) and gene body regions (TSS to transcription termination site [TTS]) were subdivided into equal-sized bins to capture fine-scale positional variation in chromatin signal. Coverage-based and fragmentomic features, including histone mark signal intensity, fragment length distributions, nucleosomal periodicity, and fragment entropy, were computed across these regions.

A total of 3,516 candidate chromatin-derived features were initially evaluated in a training cohort comprising 15 patients across seven cancer types with matched plasma cfChIP-seq and tumor RNA-seq obtained within 24 hours. Pairwise Spearman correlations between cfChIP-seq features and matched tumor RNA-seq expression (log₂[TPM+1]) were used to assess feature–expression associations. To improve promoter signal resolution, sample-specific transcription start sites were dynamically reassigned based on maximal H3K4me3 enrichment among annotated TSSs.

Gene expression inference was performed using an extreme gradient boosting (XGBoost) regression model, selected after benchmarking against alternative regression approaches due to its robustness to multicollinearity among correlated chromatin features. Model training incorporated percentile scaling of input features and cross-validated tuning of hyperparameters to minimize prediction error. Leave-one-out cross-validation (LOOCV) was used to assess performance in the training cohort.

Feature ablation analyses demonstrated that promoter- and gene body–derived features from H3K4me3 and H3K36me3 were the principal contributors to predictive accuracy, whereas enhancer features and H3K27ac provided minimal incremental value. Based on these analyses, the final APEX model was simplified to 1,532 promoter- and gene body–derived coverage and fragmentomic features from H3K4me3 and H3K36me3.

APEX achieved a median per-gene Spearman correlation of 0.80 between inferred plasma expression and matched tumor RNA-seq in the training cohort and demonstrated robust generalization across an independent validation set of 34 matched plasma–tumor samples spanning 18 cancer types. The model maintained predictive performance across a wide range of tumor fractions and outperformed individual histone mark coverage and alternative cfChIP-seq–based expression inference methods. Plasma-inferred expression values were subsequently used for downstream subtype classification and biomarker analyses (also see **Figure S2)**.

#### cfMeDIP-seq analysis

Differentially methylated sites for each SCLC subtype were retrieved from Chemi et al. and Heeke et al. ^17^. In Chemi et al. ^66^, subtype-specific hypermethylated sites were defined using pairwise subtype comparisons; regions with a β-value difference ≥0.4 and adjusted P <0.05 were considered hypermethylated in the corresponding subtype. Sites meeting these criteria in more than one subtype were discarded. From Heeke et al., we used subtype annotations previously assigned to each site, defining sites with higher methylation scores relative to other subtypes as subtype-specific. Hypermethylated sites from the two studies were combined, and sites annotated in more than one subtype were removed. Each remaining site was resized to a 400-bp window for downstream analyses. To remove regions potentially confounded by tumor fraction, we performed correlation filtering using cfMeDIP-seq data from Baca et al.^15^. cfMeDIP peaks previously called with MACS2^67^ were retrieved from that dataset. Peaks detected in more than 250 samples were selected as common peak and resized to 400 bp. Peaks overlapping high-noise genomic regions (https://github.com/Boyle-Lab/Blacklist/blob/master/lists/hg19-blacklist.v2.bed.gz) were excluded. For each sample, cfMeDIP-seq counts at common peaks were normalized by the number of unique fragments to adjust for sequencing depth. Spearman correlations between normalized signal and ctDNA fraction were then computed across 291 samples, excluding samples without ctDNA estimates or with average counts <10 across peaks. Peaks were removed if they met any of the following criteria: absolute Spearman correlation with ctDNA fraction >0.1, coefficient of variation >1, or normalized signal >200. The median count of the remaining peaks was used as a sample-specific normalization factor. Normalized methylation signal was then calculated for each subtype-specific hypermethylated site and correlated with ctDNA fraction. Sites showing strong correlation with tumor fraction (absolute Spearman correlation ≥0.5) were removed. Finally, subtype scores were computed by summing cfMeDIP-seq counts across filtered hypermethylated sites for each subtype. Scores were converted to Z-scores to enable comparison across subtypes, and the four subtype scores for each sample were transformed using a softmax function.

#### Haplotype-based BAF analyses

##### Haplotype phasing

To identify germline single-nucleotide polymorphisms (SNPs) for phasing, variants were called from BAM files using GATK (v4.6.1.0)^68^ HaplotypeCaller in GVCF mode, followed by genotyping with GenotypeGVCFs. Variant normalization was performed using bcftools (v1.21) norm^69^, and only biallelic SNPs with QUAL ≥30 and read depth ≥10 were retained for downstream phasing. Common variants were phased using SHAPEIT5 (v5.1.1) with a genetic map distributed with Eagle^70^ and a phased reference panel from the 1000 Genomes Project Phase 3 release^71^. Read-backed phasing was then refined using WhatsHap (v2.8)^72^. Phase-set blocks were extracted from phased VCF files by grouping variants sharing the same phase-set tag and defining the genomic span of each block from the minimum to maximum variant position. For haplotype-resolved read assignment, BAM files were haplotagged using WhatsHap. Within each phase-set block, haplotagged reads were counted separately for haplotypes 1 and 2, using only reads whose phase-set tag matched the corresponding block.

##### Haplotype-based BAF calculation

Allelic imbalance was assessed for each arm-level segment (≥20 Mb), under the assumption that the segment shared a common copy-number state, by calculating the haplotype 1 fraction for each phase-set block within the segment and examining the distribution of haplotype 1 fractions. For each segment, the frequency of haplotype 1 fractions was computed using bins of width 0.005. An imbalance was considered present when the mode of this distribution was at least 0.05 away from 0.5 and the number of phase-set blocks in the modal bin exceeded the maximum count within the interval 0.48 to 0.52, which was taken to represent the balanced state. When imbalance was detected, the symmetric haplotype 1 fractions were transformed to the range 0.5-1, and the BAF for the segment was defined as the median of the transformed values. Thus, in our analysis, BAF refers to the major allele frequency.

##### Allele-specific copy-number and clonality analysis

For each patient, three plasma samples were analyzed: baseline, undetectable tumor fraction (where ichorCNA predicted ctDNA content was <3%), and progression. Haplotype phasing was performed using the undetectable tumor fraction sample, and the baseline and progression samples were subsequently assigned haplotype tags based on this phasing result.

Genome segmentation for the baseline and progression samples was estimated using TitanCNA^73^. The sample with 0% tumor fraction was used as the matched normal, and TitanCNA was run in tumor–normal paired mode using 10 kb bins with default parameters. The resulting solution was used for initialization. Segments ≤1 Mb in size or supported by fewer than 1,000 SNPs were excluded. Adjacent segments predicted to share the same major and minor allele configuration and separated by ≤5 Mb were subsequently merged.

Next, under the purity (tumor fraction) estimated by the default TitanCNA solution, the major and minor allele assignments were finetuned when the observed BAF was closer to the BAF that would be expected under the alternative major and minor allele configuration with the same total copy number of a single clone. Tumor fraction estimation was then further finetuned by grid search to identify the tumor fraction value that minimized the difference between the observed BAF and the expected BAF under the inferred allelic configuration. The expected BAF was calculated by following equation

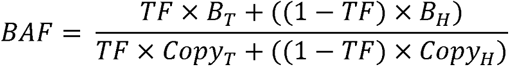

where *TF* denotes tumor fraction; B_T_ denotes the number of B alleles in tumor cells; B_H_ denotes the number of B-alleles in healthy cells, which is set to 1; Copy_T_ denotes the tumor copy number; and Copy_H_ denotes copy number in healthy cells, which is set to 2.

Segments in which the difference between the expected BAF and the observed BAF exceeded 0.03 were considered not to be explainable by a single clonal population. For patient 6527, the subclonal fraction was estimated using the chromosome 15 segment where the observed BAF suggested the presence of a subclone. Because the coverage of chromosome 15 was comparable to that of other segments inferred to have the same total copy number (two copies), the subclone was assumed to also carry two copies. The allelic configuration and subclone fraction that minimized the discrepancy between the observed and expected BAF for the chromosome 15 segment were then determined by grid search.

#### Inference of SCLC Molecular Subtypes from Plasma APEX Profiles

##### General framework

Given the focus of this analysis was on subtyping SCLC, only patients with SCLC were included and patients with poorly differentiated neuroendocrine carcinoma and carcinoid were excluded. Moreover, patient (7242) with undetectable tumor fraction was also excluded as well as the outlier patient 7181. SCLC molecular subtypes were inferred from baseline plasma samples using a hybrid approach that integrates a supervised random forest (RF) classifier trained on tumor RNA-seq data from the IMpower133 tumor cohort with established subtype-specific transcriptional gene signatures^33^. For each sample, the subtype with the highest predicted probability was initially assigned. Predictions were considered confident when the difference between the highest and second-highest predicted probability was ≥0.1, in which case the subtype with the highest probability was retained. For samples in which the difference between the two highest subtype probabilities was <0.1, additional gene signature–based criteria were applied to refine subtype assignment. When the ambiguity involved SCLC-A versus SCLC-N, a subtype score ratio was calculated using previously published gene signatures^33^. Specifically, the mean expression of a 50-gene SCLC-A signature and the mean expression of a 50-gene SCLC-N signature were computed for each sample, and the A/N ratio was derived. Samples with a ratio >1 were classified as likely-A, whereas those with a ratio <1 were classified as likely-N. For cases involving ambiguity between other subtype combinations, subtype assignment was guided by the expression of lineage-defining transcription factors. Log2-transformed TPM expression levels of *POU2F3*, *ASCL1*, *NEUROD1*, and *YAP1* were examined, and samples were classified as likely belonging to the subtype corresponding to the transcription factor with the highest expression.

##### Relative subtype probabilities

Relative subtype probabilities were calculated to facilitate comparison of subtype dominance within individual samples. For each sample, the predicted probability of each subtype was divided by the maximum subtype probability for that sample, resulting in values scaled between 0 and 1, where the dominant subtype equals 1 and other subtype probabilities are expressed relative to it. This normalization was used for visualization of subtype patterns across patients.

##### Random Forest–Based Inference

We trained a supervised machine learning model using tumor RNA-seq data with established subtype annotations, as defined in Gay et al.^13^. Tumor tissue RNA-seq TPM values were retrieved from the IMpower133 dataset, and cfChIP-seq expression like data were extracted from plasma-derived samples profiled using the APEX model. Clinical subtype labels for tissue samples were harmonized with corresponding expression data by matching sample identifiers. All gene expression matrices (tumor tissue and cfChIP-seq) were filtered to retain only genes shared across both platforms. We randomly partitioned the tumor tissue dataset into a training set (80%) and a validation set (20%). Gene-wise variance was calculated from the training set, and a grid search was performed to evaluate classification performance using the top n most variable genes, with n ranging from 100 to 10,000 in increments of 100. For each value of n, a Random Forest model (R package randomForest^74^, ntree = 400) was trained on the training data and evaluated on the held-out validation set. Model performance was assessed using validation accuracy. The optimal mode, yielding the highest validation accuracy used the top 1000 most variable genes. Using these 1000 genes, a final Random Forest model was trained on the full training set (ntree = 500). This model was then applied to predict SCLC subtype probabilities in plasma samples. Subtype assignment for each plasma sample was based on the maximum predicted probability. Final results included per-sample predicted subtype and associated class probabilities (**Figure S2**).

##### Gene signatures

For plasma samples in which the difference between the highest and second-highest subtype probabilities predicted by the random forest model was <0.1, subtype assignment was further refined using transcriptional gene signatures described by Nabet et al.^33^. The top 50 genes most specific to the SCLC-N (NMF1) and SCLC-A (NMF2) subtypes were used to define subtype signatures. For each sample, average signature expression was calculated, and samples were classified as Likely-A or Likely-N based on higher expression of the corresponding subtype signature.

##### WBC and SCLC Gene Lists

The SCLC-specific gene signature was derived from the Curated Cancer Cell Atlas^75^, while the white blood cell gene signature was sourced from the GTEx datasets. The data used for the analyses described in this manuscript were obtained from the GTEx Portal on 10/04/25.

#### Mouse Models

All mouse experiments complied with National Institutes of Health guidelines and were approved by Dana-Farber Cancer Institute Animal Care and Use Committee (DFCI, protocol 19-009). For the cell line subcutaneous syngeneic mouse tumor studies in BL6J mice, 7-9 week-old female albino BL6 mice were purchased from Jackson Laboratories (Jackson Lab# 00058). For the cell line subcutaneous syngeneic mouse tumor studies in BL6J mice with humanized CD3 replacing mouse CD3, 6 week old female were purchased from Biocytogen (Biocytogen #110039 B-hCD3EDG mice, strain C57BL/6-Cd3e<tm1(CD3E)Bcgen>, Cd3d<tm1(CD3D)Bcgen>, Cd3g<tm1(CD3G)Bcgen>/Bcgen.) or mixed gender mice at roughly equal male to female ratios from genOway [(genOway, genO-panhCD3 humanized mouse model, C57BL/6N-Cd3etm2.1(CD3E)Geno;Cd3gtm2.1(CD3G)Geno;Cd3dtm2.1(CD3D)Geno)] as described in each figure legend. Housing conditions for mice at the Dana-Farber Cancer Institute include a 12 h/12 h day-night cycle where temperature is maintained at 72 Fahrenheit.

#### Cell Lines

##### Murine SCLC Cell Lines

All murine neuroendocrine (NE) *RPR2* SCLC cells lines were maintained in RPMI-1640 (Life Technologies, #11875135) media with 10% FBS (Gemini, #100-106) containing HITES [10 nmol/L hydrocortisone (Sigma Aldrich, #H0135), Insulin-Transferrin-Selenium (Gemini; #400-145), and 10 nmol/L β-estradiol (Sigma Aldrich, #E2257)] and with 100 U/mL penicillin, and 100 μg/mL streptomycin (P/S) (Gibco, #15140122) in 37°C in 5% CO_2_ as described previously^35,36,51^. The 1014P2 *RPR2* cell line was generated as described previously^37^. Briefly, an sgControl *RPR2* adenovirus was intratracheally injected into pure congenic Lox-stop-lox (LSL) Cas9 BL6J mice purchased from Jackson Labs (Jackson No. 026175) and an autochthonous SCLC tumor developed from which a cell line was established (1014 cells) as described previously^35,36,51^. 1014P2 cells were then made from 1014 cells by passaging through BL6 mice twice to increase engraftment allowing a near 100% take rate for syngeneic mouse tumor studies in BL6 mice as described previoulsy^37^.

The 1014P2-TR#2 (tarlatamab resistant) cell line was established as follows: 8 million parental 1014P2 cells were subcutaneously injected into the flank of a male genOway (genO-panhCD3 humanized mouse model, C57BL/6N Cd3etm2.1(CD3E)Geno;Cd3gtm2.1(CD3G)Geno;Cd3dtm2.1(CD3D)Geno) mouse with 33% Matrigel (Corning, #356231) diluted in PBS. The mouse was treated with 0.3mg/kg of tarlatamab by intraperitoneal (IP) injection once a week. The mouse initially had a complete response to tarlatamab. After the complete response, the mouse was maintained on tarlatalab (0.3 mg/kg/week by IP injection) and eventually developed acquired resistance with tumor re-growth at which point (66 days after first treatment) the mouse was euthanized and the 1014P2-TR#2 cell line was established.

The 1014P2-TR#4L and 4R (tarlatamab resistant) cell line was established from a female genOway (genO-panhCD3) mouse treated with 3mg/kg of tarlatamab by IP injection once a week for three times and observed complete response. After followup of over 150 days post inoculation and confirming no signs of recurrence, the mouse was rechallenged with bilateral injection of 8 million parental 1014P2 cells. Tumor growth was monitored and after tumors were well established (>1000mm^3^) the mouse was treated with weekly tarlatamab 0.3mg/kg, IP. Both tumors initially showed a response with tumor shrinkage to tarlatamab but regrew and was sacrificed 24 days after retreatment and cell lines from both tumors were established.

The 1014P2-TR#5 and 5H Tarlatamab resistant) cell line was established from a female genOway (genO-panhCD3) mouse treated with 0.3mg/kg of tarlatamab by IP injection once a week for three times and observed complete response. After followup of over 150 days post inoculation and confirming no signs of recurrence, the mouse was rechallenged bilaterally with 8 million parental 1014P2 cells. Only left side tumor slowly established and after tumor size exceeded 500mm^3^, the mouse was treated with weekly tarlatamab 0.3mg/kg, IP. The mouse showed a limited response to tarlatamab and was sacrificed 16 days after retreatment and cell lines were established as mentioned above. Dissection showed two individual tumors in the left flank and established cell lines from each.

The 1014P2-TR#6 Tarlatamab resistant) cell line was established from a male genOway (genO-panhCD3) mouse treated with 0.3mg/kg of tarlatamab by intraperitoneal (IP) injection once a week for three times and observed complete response. After followup of over 150 days post inoculation and confirming no signs of recurrence, the mouse was rechallenged bilaterally with 8 million parental 1014P2 cells. Both tumors slowly established and after the tumors were well established (>1000mm^3^), the mouse was treated with weekly tarlatamab 0.3mg/kg, IP. The mouse showed a remarkable response with both tumors completely rejected. Despite continued tarlatamab treatment, right side tumor showed recurrence and was sacrificed 45 days after retreatment and cell lines were established.

Early passage cells of all cell lines listed above were tested for Mycoplasma (Lonza, #LT07-218) and then were frozen using Bambanker’s freezing media (Bulldog Bio, # BB01). All experiments were performed with cell lines that were maintained in culture for <3 months at which time an early passage cell line was thawed.

#### Murine tumor dissociation

Mice were euthanized with CO_2_ followed by neck dislocation and the tumors were quickly removed under sterile condition onto a 10 cm dish with 1 mL of PBS. A portion of the tumor was taken and minced with sterile razor blades and used for single-cell dissociation using the Tumor Dissociation Kit, mouse (Miltenyi Biotec, #130-096-730), transferred into a gentle MACS C Tube (Miltenyi Biotec, #130-093-237) added with 2.25 mL of RPMI 1640, 100 µL of Enzyme D, 10 µL of Enzyme R, and 12.5 µL of Enzyme A and set on a gentleMACS Dissociator with heater and run under the program “37C_m_TDK_2”. After dissociation, cell suspension was filtered through a 70 µm cell strainer (Thermo Fisher Scientific, #22363548) and washed with 20 mL of RPMI 1640 medium. Following centrifuge at 300 x g for 7 minutes, supernatant was removed and cells were used for flow cytometric analysis or cell line generation.

#### Lentiviral sgRNA Expression Vectors and sgRNA Sequences

sgRNA sequences were designed using the CRISPick sgRNA design tool from BROAD Institute (https://portals.broadinstitute.org/gppx/crispick/public). Oligo DNAs were synthesized from IDT (Integrated DNA Technologies). 11.25 µL of 100mM sense and antisense oligonucleotides were mixed in an eppendorf tube with 2.5 µL of annealing buffer (1x annealing buffer: 100 mM NaCl, 10 mM Tris-HCl, pH 7.4) and heated at 100°C. After slowly cooling to 30 °C by leaving at room temperature for over 3 hours, tubes were briefly spin and diluted 1 : 400 with 0.5x annealing buffer. 1 mL of the diluted annealed oligo was ligated into Esp3I digested pLentiGuide-Puro vector (Addgene; #52963) by adding 1 µL of the digested vector, 1 µL of 10x ligase buffer, 1 µL of 10mM ATP, 1 µL of T4 DNA ligase and 6 µL of DNase/RNase free water and incubating at room temperature for 2 hours. Ligation mixture was then transformed into HB101 competent cells by heat shock method. After recovery of the competent cell with LB broth and shaking for 1 hour at 37°C, HB101 cells were plated onto an ampicillin LB agar plate and incubated at 37°C overnight. Colonies were picked up and expanded for miniprep (QIAprep Spin Plasmid Miniprep Kit, Qiagen, #27106) and validated by DNA sequencing. The following sgRNA oligos were used. DLL3 sgRNA (sg1 sense: 5’ CACCGGTGGCCTCCTGGGAGACTCC 3’, sg1 antisense: 5’ AAACGGAGTCTCCCAGGAGGCCACC 3’, sg2 sense: 5’CACCGCCTGAAGCCCGGAGTCTCCC3’, sg2 antisense: 5’ AAACGGGAGACTCCGGGCTTCAGGC 3’) and Control sgRNA (sg1 sense: TTCGAGATTTTAATAGCGCA, sg1 antisense: TGCGCTATTAAAATCTCGAA, sg2 sense: TGAGGTTACCGCCGGCTTTT, sg2 antisense: AAAAGCCGGCGGTAACCTCA)

#### NEUROD1 over expression vector and sequence

The mouse NEUROD1 over expression vector was ordered through TWIST Bioscience by inserting the Kozak - mouse NEUROD1 – HA tag – HA tag sequence (GCCACCATGACAAAGTCCTATTCAGAATCTGGATTGATGGGCGAACCCCAACCTCAGGGGCCTCCCTCCTGGACCGACGAATGCCTGAGCAGCCAAGATGAAGAGCATGAAGCCGATAAGAAGGAAGATGAACTTGAGGCTATGAACGCGGAAGAAGATAGCCTTCGCAATGGTGGGGAAGAAGAAGAAGAGGACGAAGACCTCGAAGAGGAAGAAGAGGAAGAGGAAGAAGAAGAAGACCAGAAACCAAAACGACGTGGGCCTAAGAAGAAGAAAATGACTAAAGCCCGACTGGAGAGATTCAAGCTTAGACGAATGAAAGCAAATGCACGGGAACGAAATAGAATGCATGGCCTAAATGCTGCTTTGGATAATCTAAGGAAAGTCGTTCCGTGTTATAGTAAAACGCAAAAGCTCTCCAAGATTGAAACCCTACGGCTTGCAAAGAATTATATATGGGCCCTTTCCGAAATTCTTAGGAGCGGAAAGAGTCCGGACCTTGTGAGTTTTGTCCAAACCCTGTGTAAGGGACTGAGCCAACCAACGACAAACCTTGTGGCGGGGTGTTTGCAATTGAATCCCAGGACCTTTCTCCCCGAACAAAATCCAGATATGCCACCTCACTTGCCGACAGCTTCCGCCAGTTTTCCTGTCCACCCGTATTCTTATCAAAGTCCAGGCCTACCAAGTCCACCATATGGTACAATGGATAGTAGTCATGTGTTTCATGTTAAACCACCACCGCATGCTTATTCTGCCGCACTCGAACCGTTCTTCGAGTCACCGCTGACCGATTGTACGTCCCCATCATTCGATGGGCCGCTGTCACCACCTCTGTCAATTAACGGGAATTTCTCATTTAAGCATGAGCCGTCTGCTGAATTCGAGAAGAACTACGCGTTCACGATGCATTATCCGGCTGCTACCCTTGCGGGCCCACAGTCTCATGGGTCCATATTTAGCAGCGGAGCAGCCGCTCCCAGGTGTGAAATTCCTATTGATAATATAATGTCCTTTGACAGTCACTCTCACCACGAACGGGTGATGTCCGCGCAACTGAACGCTATTTTCCATGACTACCCCTACGACGTGCCCGACTACGCCGGCTATCCGTATGATGTCCCGGACTATGCATAA) into the pTwist Lenti SFFV puro WPRE followed by codon optimization. Lentivirus was produced and 1014P2 cells were transduced with the following methods.

#### Lentiviral production

Lentiviruses were produced by transfecting 293T cells with the lentiviral expression vector, packaging plasmids psPAX2, and pMD2.G (Addgene, #12260) and pMD2.G (Addgene, #12259) at a ratio of 4:3:1 and co-transfecting with Lipofectamine 2000 (Thermo Fisher Scientific, #11668019). Virus-containing media was collected 48 and 72 hours post-transfection and passed through a 0.45 SFCA filter (CellTreat, #229766) and used directly or stored at -80°C until later usage.

#### Lentiviral Transduction

2 million cells were resuspended in 1 mL of lentivirus containing media with 8 μg/mL of polybrene and plated into a 12 well plate. After centrifugation at 2000 rpm 30°C for 2 hours, cells were incubated overnight and media was replaced the following day and cells were expanded into a T25 or T75 flask. On day 3, 1 μg/mL of puromycin was added for selection.

#### SCLC Cell Line Syngeneic Mouse Tumor Experiments in BL6 Mice

SCLC syngeneic mouse tumors were established as following. 1014P2 or 1014P2-TR#2 cells were cultured and expanded to desired number of cells in a T225 flask. On day of inoculation, cells were collected into 50mL conical tubes and washed with PBS followed by centrifuge at 1500 rpm for 4 min room temperature for two times. After the second wash, cells were resuspended in 50 mL of PBS and counted in duplicate. Cells were centrifuged one last time at same condition and resuspend in PBS to make a final concentration of 8 million cells/66 μL. Cells were mixed with Matrigel (Corning, #356231) at 2:1 volume/volume ratio and 100 μL of the mixture (= 8 million cells) was injected subcutaneously into the flank of 7-9 week-old female albino BL6 mice (Jackson Lab# 00058), or humanized CD3 mice [(Biocytogen #110039 B-hCD3EDG mice, strain C57BL/6-Cd3e<tm1(CD3E)Bcgen>, Cd3d<tm1(CD3D)Bcgen>, Cd3g<tm1(CD3G)Bcgen>/Bcgen)] or [(genOway, genO-panhCD3 humanized mouse model, C57BL/6N-Cd3etm2.1(CD3E)Geno;Cd3gtm2.1(CD3G)Geno;Cd3dtm2.1(CD3D)Geno mice through a 27G needle.

#### Tarlatamab *In Vivo* Efficacy Experiments

For efficacy experiments in humanized CD3 BL6 mice from Biocytogen or genOway, or in BL6J mice from Jackson Laboratories, 1014P2 cells were subcutaneously injected into the mice as described above and tumors were allowed to establish to (Biocytogen 129 +/- 83 mm^3^, genOway 236+/- 73 mm^3^, BL6J 169 +/- 40 mm^3^) in size before randomization into tarlatamab or vehicle (PBS) by IP injection once a week at the doses indicated. Throughout the experiment, tumor diameters were measured twice a week using calipers and tumor volume was calculated using the following formula; Tumor volume (mm^3^) = width (mm) x length(mm) x height(mm) x π/6. Body weight was measured once a week. The experimental endpoint was when tumors were over 2cm in maximum diameter at which point the mouse was euthanized. For downstream analysis of the tumor, after euthanization, tumors were immediately dissected and ∼1/3 of the tumor was flash frozen on dry ice, ∼1/3 of the tumor was fixed in 10% formalin for 24 hours then stored in 70% ethanol for histological analysis and IHC, and ∼1/3 of the tumor was used for single cell dissociation.

#### Tarlatamab *In Vivo* Competition Experiments

1014P2 and 1014P2-TR#2 cells were subcutaneously injected into opposite bilateral flanks of the same humanized CD3 BL6 mouse (one cell line per flank; one female genOway mouse, one male genOway mouse). Once tumors were established (232 +/- 72 mm^3^), mice were treated with tarlatamab (0.3 mg/kg) or vehicle (PBS). Tumors were harvested for IHC and flow cytometric analysis when the 1014P2-TR#2 tumor exceeded 2,000 mm³.

#### *In Vivo* Rechallenge experiments

Mice which had complete response after three doses of tarlatamab from experiments shown in **Figure 2J-2M** were followed up for over 150 days to confirm no evidence of recurrence. These mice were rechalleged with parental 1014P2 by preparing cells as described in above section “SCLC Cell Line Syngeneic Mouse Tumor Experiments in BL6 Mice”. After injection, tumor size was measured twice a week. When tumors were well established (1579 +/- 733mm^3^), mice were started with weekly tarlatamab 0.3 mg/kg IP injection until endpoint.

#### Neurod1 overexpression *In Vivo* experiments

8 million Neurod1 overexpressed 1014P2 cells (1014P2 NEUROD1-OE cells) were subcutaneously injected into the flanks of humanized CD3 mice with mixed gender from genOway. After tumors were established (256 +/- 128 mm^3^) mice were randomized to receive either vehicle (PBS) or tarlatamab (0.1 mg/kg) weekly IP.

#### RNA-Seq

Tumor and cell line samples were collected for RNA sequencing. Tumors included two independent vehicle (PBS)-treated 1014P2 syngeneic mouse tumor and one tumor that developed acquired resistance to tarlatamab (1014P2-TR#2). Cell line samples included the parental 1014P2 cell line (control) and a cell line derived from 1014P2-TR#2 (two biological replicates). Tumor tissues were snap frozen and stored at −80°C until processing. Cell lines were prepared fresh for RNA isolation. For RNA extraction, snap-frozen tumor tissues were processed directly from −80°C storage. For cell line samples, cells were plated in 6-well plates at 200,000 cells/mL and harvested 24 hours later. Total RNA was extracted using the Quick-RNA Miniprep Kit (Zymo; #R1055). Purified RNA was submitted to Novogene Inc. for sequencing. For non strand specific library, messenger RNA was purified from total RNA using poly-T oligo-attached magnetic beads. After fragmentation, the first strand cDNA was synthesized using random hexamer primers followed by the second strand cDNA synthesis. The library was ready after end repair, A-tailing, adapter ligation, size selection, amplification, and purification. The library was checked with Qubit and real-time PCR for quantification and bioanalyzer for size distribution detection. For strand specific library, messenger RNA was purified from total RNA using poly-T oligo-attached magnetic beads. After fragmentation, the first strand cDNA was synthesized using random hexamer primers. Then the second strand cDNA was synthesized using dUTP, instead of dTTP. The directional library was ready after end repair, A-tailing, adapter ligation, size selection, USER enzyme digestion, amplification, and purification. The library was checked with Qubit and real-time PCR for quantification and bioanalyzer for size distribution detection.Sequencing reads were mapped to the mm39 genome using HISAT2 (2.2.1) and statistics for differentially expressed genes were calculated using DESeq2 (1.42.0).

#### Chromatin Preparation from Mouse SCLC Cell Lines

Approximately 1 × 10LJ cells were crosslinked by adding formaldehyde directly to the culture medium to a final concentration of 1% and incubating for 10 min at room temperature with gentle agitation. Crosslinking was quenched by addition of glycine (final concentration 125 mM) followed by a 5-min incubation at room temperature. Cells were washed twice with ice-cold PBS containing complete protease inhibitors (Sigma-Aldrich, Cat. #11873580001), scraped, and pelleted by centrifugation (625 × g, 4 min, 4 °C). Pellets were resuspended in 1 mL dilution buffer (1% NP-40, 0.5% sodium deoxycholate, 0.1% SDS, and complete protease inhibitors in PBS) and incubated on ice with gentle rocking for 10 min. Chromatin was sheared using a Covaris AFA system (peak incident power 140, duty factor 5%, 200 cycles per burst) for 25 min at 4 °C. Following sonication, lysates were clarified by centrifugation (13,000 × g, 10 min, 4 °C), and the supernatant containing soluble chromatin was collected and stored at −80 °C until further use. ChIP-sequencing for H3K4me3, H3K27Ac, and H3K36me3 was performed as described in the cfChIP-seq method above with the following modification. For each wash with low-salt, high-salt, and LiCl buffers, samples were incubated for 10 minutes at 4 °C rather than for 1 minute. In addition, incubation with the extraction buffer was extended to 30 minutes at 37 °C, 1 hour at 55 °C, and 8 hours at 65 °C.

#### ChIP-seq analysis of mouse cell lines

Adapter sequences were trimmed using fastp (v0.23.2) with default parameters. Reads were aligned to the mm10 reference genome using BWA (v0.7.19) with default parameters. Read pairs were retained if they were properly paired and had a mapping quality greater than 20. Read pairs aligned to the exact same position were considered PCR duplicates and discarded. Each read pair was represented as a single fragment in BEDPE format. Mouse protein-coding genes were retrieved from GENCODE (version M23)^76^. Duplicated genes were discarded, and promoters were defined as ±1 kb around the transcription start site. Fragment midpoint coverage across promoters was calculated for each gene in each sample; these values were used as the ChIP-seq signal. The ChIP-seq signals were normalized as follows: each signal was divided by the corresponding gene-level signal in the input sample; if the input sample did not show any fragment on a gene promoter, the ChIP-seq signal was divided by 1. Subsequently, ChIP-seq signals were log_2_(x + 1)-transformed. The mean signal across genes was then used to scale each sample to make samples comparable. The top 50 genes most specific to the SCLC-N (NMF1) and SCLC-A (NMF2) subtypes were used to define subtype signatures and then converted to mouse orthologs . NEUROD1 and ASCL1 Orthologs of SCLC subtype signature genes were retrieved from the BioMart database^77^ using the pybiomart package. Gene set enrichment analysis (GSEA) was performed using GSEApy^78^, using the difference in ChIP-seq signal between 1014P2 and 1014P2-TR#2 to rank genes for each signature gene set.

#### Bulk RNA-Sequencing Analysis

Bulk RNA-sequencing heatmaps in **Figure 4E** and **Figure S4C** were generated by calculating z score from log transformed FPKM values of differentially expressed genes (p value<0.05). To calculate z score, a data point was subtracted the mean and then divide the result by the standard deviation.

#### Gene Set Enrichment Analysis

GSEA software was downloaded from the Gene Set Enrichment Analysis website [http://www.broad.mit.edu/gsea/downloads.jsp]^79^. Pre-ranked GSEA was performed using hallmark gene sets from the human MSigDB Collections32 in **Figure 4F** and **Figure S4D.**

#### Single-Cell RNA-Seq

Surgically resected patient tumor (cohort ID 7373) was freshly obtained and dissociated into single cell using the human tumor dissociation kit (Miltenyi Biotech, #130-095-929) within two hours of resection by running on gentleMACS™ Octo Dissociator with the “37C_h_TDK_2” program. After washing and filtering, dead cell removal was performed using the MojoSort™ Human Dead Cell Removal Kit (Biolegend, #480159). Number of live cells were counted and resuspended in PBS + 0.04% BSA at 1000 cells/μL for sequencing.

Single-cell RNAseq (scRNA-seq) gene expression (GEX) libraries were generated using the 10x Chromium GEM-X Single Cell 5’ Reagent Kit v3 (10x Genomics, #PN-1000699) according to the manufacturer’s protocol. Briefly, a suspension targeting 10,000 cell recovery was loaded onto a Chromium GEM-X single cell 5’ Chip (PN-1000698) and processed on a Chromium X Controller. In this step, single cells were partitioned into Gel Bead-in-emulsions (GEMs), where the RNA was reverse transcribed using primers containing a 10x Barcode, a unique molecular identifier (UMI), and a template switch oligo to generate barcoded cDNA. After breaking the emulsion, the pooled cDNA was purified and amplified by PCR. GEX libraries were then constructed from the amplified cDNA using the Library Construction Kit C (PN-1000694). Finally, sample indexing PCR was performed using the Dual Index Kit TT Set A (PN-1000215) to barcode the sequencing libraries. The final libraries were sequenced on a NovaSeq X (Illumina) with paired-end, dual-index reads, using the following parameters: Read 1: 28 cycles, index 1:10 cycles, index 2: 10 cycles, and Read 2: 90 cycles. The scRNA-seq data was processed using the cellranger version 8.0.1 (10x Genomics).

#### Single-Cell RNA Sequencing Data Analysis

Filtered raw counts data was imported into Seurat v4 R package (version 4.1.0) for downstream processing and analysis. Low quality cells were filtered out using following thresholds: total UMI counts < 500, number of transcripts < 200, log10TranscriptsPerUMI <= 0.7, and cells with more than 20% transcripts mapping to mitochondrial genes. In addition, genes expressed in less than three cells were removed. The UMI counts matrices were then natural-log normalized and scaled with Seurat’s ‘NormalizeData’ and ‘ScaleData’ functions.

Dimension Reduction, Cluster Analysis and Visualization of scRNA-Seq

The Seurat v4 R package was used for dimension reduction and clustering. Top 2,000 genes with the highest variance were selected using the ‘FindVariableFeatures’ function with ‘vst’ method to perform linear dimensional reduction (principal component analysis) using the ‘RunPCA’ function, and top 17 principal components were used to perform graph-based unsupervised clustering with the ‘FindNeighbors’ and ‘FindClusters’ functions and Uniform Manifold Approximation and Projection for Dimension Reduction (UMAP) (arXiv:1802.03426) for data visualization in two-dimensional space. Automatic cluster annotation using the R package SingleR (version 1.8.1) with ImmGenData and manual tumor cluster annotation were performed to annotate cell population. Subsequently, the T and NK subpopulations were reanalyzed. Top 2,000 genes were selected using the ‘FindVariableFeatures’ function with ‘vst’ method to perform linear dimensional reduction (principal component analysis) using the ‘RunPCA’ function, and top 14 principal components were used to plot UMAP for data visualization in two-dimensional space. Differential expression profiles were obtained with Seurat’s ‘FindMarkers’ function to annotate subpopulations of T and NK cells. Seurat’s ‘FeaturePlot’ function was used to visualize single cell level expression in **Figure 3O-3P** and **Figure S3G-S3H**.

#### Reverse-Transcriptase Quantitative PCR (RT-qPCR)

RNA was extracted using Quick-RNA Miniprep Kit (Zymo; #R1055) according to the manufacturer’s instructions. cDNA was synthesized using iScript Reverse Transcription Supermix for RT-qPCR (Bio-Rad; #1708841). qPCR was performed using Taqman (Thermo Fisher Scientific) pre-designed probes with indicated primer oligos according to the manufacturer’s instructions. The ΔΔCT method was used to analyze the data. The CT values for each probe were normalized to the CT value of ACTB for that sample. The data from each experiment was then normalized to the control (1014P2) to determine the relative fold change in mRNA expression. Primers; Ascl1 (Mm0358063_m1), Neurod1 (Mm01280117_m1), Dll3 (Mm00432854_m1), ActinB (Mm00607939_s1).

#### Amplicon sequencing for Dll3 in 1014P2 and 1014P2-TR#2 cell line

Genomic DNA was extracted from 1014P2 and 1014P2-TR cell line using the QIAmp DNA Blood Mini Kit (QIAGEN, #51104). Target site on Dll3 was amplified by PCR using the KOD XTREME HOT START DNA POLYMERASE (Sigma Aldrich, #71975) with the following primers; Dll3 Fw primer 5’ GCTGATCCTGGCTTTTCTTCT 3’, Dll3 Rv primer 5’ CCTGGAGATGGAGCCTGAAT 3’,which covers the Dll3 genomic DNA region from 75 – 7923 with a PCR product of 7849bp. The PCR product was run on an agarose gel and column purified using QIAquick Gel Extraction kit (Qiagen, #28704) and sent to the MGH DNA Core Facility for amplicon sequencing using next generation sequencing. Data was analyzed using MGH DNA Core’s complete amplicon sequencing pipeline.

#### Flow cytometry

Single-cell suspensions were washed once with 5 mL of PBS and centrifuged at 1,500 rpm for 3 min. After removal of the supernatant, cells were resuspended in PBS containing Zombie Violet Fixable Viability Kit (Biolegend, #423113) or Zombie NIR Fixable Viability Dye (BioLegend, #423106) at a 1:500 dilution and incubated for 10 min at room temperature in the dark. Samples were then washed once with 2 mL of FACS buffer (PBS supplemented with 2% FBS). Following centrifugation at 1,500 rpm for 3 min, cells were resuspended in FACS buffer and incubated with anti-mouse CD16/32 antibody (Fc block; BioLegend, #101320) at a 1:100 dilution for 30 min at 4°C in the dark. Surface staining was performed by adding fluorochrome-conjugated antibodies directly to the samples and incubating for additional 30 min at 4°C in the dark. After surface staining, samples were washed twice with FACS buffer, resuspended in an appropriate volume of buffer, and passed through a cell strainer prior to flow cytometric analysis.

For intracellular staining, cells were fixed and permeabilized using the eBioscience™ Foxp3/ Transcription Factor Staining Buffer Set (Thermo Fisher Scientific, #00-5523-00) according to the manufacturer’s instructions. Briefly, after the final surface-staining wash, cells were resuspended in 1 mL of Foxp3 Fixation/Permeabilization working solution and incubated overnight at 4°C protected from light. The following day, samples were washed twice with 2 mL of 1× Permeabilization Buffer, with centrifugation at 1,500 rpm for 3 min at room temperature between washes. Cells were then resuspended in residual 1× Permeabilization Buffer, incubated with intracellular antibodies for 60 min at room temperature in the dark, and washed twice with 1× Permeabilization Buffer. Finally, cells were resuspended in an appropriate volume of FACS buffer and passed through a cell strainer prior to flow cytometric analysis.

For surface staining from cell lines, 1-2 million cells were collected, centrifuged, and washed with PBS. After removal of the supernatant, cells were resuspended in PBS containing Zombie Violet Fixable Viability Kit (Biolegend, #423113) or Zombie NIR™ Fixable Viability Dye (BioLegend, #423106) at a 1:500 dilution and incubated for 10 min at room temperature in the dark. Samples were then washed once with 2 mL of FACS buffer (PBS supplemented with 2% FBS). Surface staining was performed by adding fluorochrome-conjugated antibodies to the samples and incubating for 30 min at 4°C in the dark. After staining, samples were washed twice with FACS buffer, resuspended in an appropriate volume of buffer, and passed through a cell strainer prior to flow cytometric analysis.

Antibodies used were Zombie Violet Fixable Viability Kit (Biolegend, #423113), Zombie NIR Fixable Viability Kit (BioLegend, #423105), PerCP/Cyanine5.5 anti-mouse CD45 (BioLegend, (#103132), Brilliant Violet 650 anti-human CD3 (BioLegend, #300467), Brilliant Violet 421 anti-mouse CD4 (BioLegend, #100437), Brilliant Violet 711 anti-mouse CD8a (BioLegend, #100747), PE anti-mouse DLL3 (BioLegend, #154003), PE Rat IgG2b kappa, Isotype Ctrl (BioLegend, #400607), Alexa Fluor 647 anti-mouse FOXP3 (Biolegend, #126407), Brilliant Violet 785 anti-mouse PD-1 (Biolegend, #135225), Brilliant Violet 605 anti-mouse Lag3 (Biolegend, #125257), APC anti-mouse Tim3 (Biolegend, #119705). BD LSRFortessa™ Cell Analyzer and MACSQuant® Analyzer 10 Flow Cytometer were used for flow cytometery and data was analyzed using FlowJo ver10 software (https://www.flowjo.com).

#### Tarlatamab binding assay

Cells (either mouse SCLC cells or T cells) were collected and washed twice with PBS, and incubated with Tarlatamab (MedChemExpress, #HY-P99575) at a final concentration of 200 nM for 1 hour on ice. Samples were washed twice with FACS buffer and subsequently incubated with anti-Fc antibody (Jackson ImmunoResearch, #109-605-008, Alexa Fluor® 647 AffiniPure® Goat Anti-Human IgG, Fcγ fragment specific) at 1:400 volume/volume ratio on ice for 1 hour. After incubation, cells were washed with FACS buffer twice and resuspended in appropriate volume of FACS buffer for flow cytometric analysis.

#### Human T cell isolation

Human T cells were isolated fresh from buffy coats prepared from whole blood collected by the Kraft Blood Center at Dana-Farber Cancer Institute Boston MA, in accordance with Institutional Review Board (IRB) protocol # T0761 by utilizing the human T cell enrichment kit Rosettesep (STEMCELL technologies, #15021) following the manufacturer’s instructions. Briefly, the ends of the collar were trimmed and drained into a 50 mL conical tube. Human T cell isolation was performed from healthy donor collar by utilizing the human T cell enrichment kit Rosettesep (STEMCELL technologies, #15021) following the manufacturer’s instructions. Briefly, the ends of the collar were trimmed and drained into a 50 mL conical tube. 50 μL of Rosettesep per 1 mL of blood was added and mixed well. After incubation at room temperature for 10 min, blood was diluted with an equal amount of RPMI + 2%FBS and layered on top of 15 mL of Ficoll (MilliPore Sigma, #GE17-1440-02) in a sepmate tube (Stemcell, # 85450) via pipetting down the wall of the tube, with the tube angled at 45 degrees. Sepmate tube was centrifuged at 1200 x g for 10 minutes and the top layer from spin was poured into a new 50 mL conical tube and added another 1:1 dilution with 2% RPMI for wash. After centrifuge 300 x g for 10 minutes, the T-cells were resuspended in 1 mL of T-cell growth medium and counted for downstream applications.

#### Mouse T cell isolation

EasySep Mouse T-cell isolation kit (StemCell Technologies, # 19851) was used to isolate T cells following the manufacturer’s instructions. Briefly, after harvesting the spleen and mechanically dissociating into single cells, 150 million splenocytes were resuspended in 1.5 mL of isolation buffer (PBS + 2%FBS + 1mM EDTA). 30 μL of FcR blocker was added to the sample and mixed gently. 75μl of Isolation Cocktail was added to sample and mixed gently followed by incubation for 10 minutes at room temperature. 112.5 μL of RapidSpheres was added to the sample and incubated for 2.5 minutes at room temperature. 1 mL of isolation buffer was added and the tube was incubated in a magnetic stand for 2.5 minutes at room temperature. After incubation, the magnet and tube were inverted, and the enriched cell suspension was poured into a new tube and used for downstream assays.

#### *In vitro* coculture of murine SCLC cells and human T cells

A polyclonal population of murine the 1014P2 SCLC cell line was transduced with the sgControl#2 sgRNA (for DLL3 WT) or sgDll3#2 sgRNA [for DLL3 knockout (KO)], selected with puromycin, and then immunoblotted to confirm DLL3 KO. These DLL3 isogenic 1014P2 cells were then cocultured with isolated human T cells for 48 hours with Tarlatamab at concentrations of 0nM, 0.001nM, 0.01nM, 0.1nM, 1nM, 10nM, and 100nM. Human T cells were thawed and rested in T cell medium [(RPMI + 10% FBS + 1% PS supplemented with 200U/mL of human IL-2 (Miltenyi Biotec, #130-097-743)] and incubated at 37 °C one day before coculture. The next day T cells were counted on Vicell and resuspended in RPMI at 1 million/mL (for E:T ration of 2:1) and 2.5 million/mL (for E:T ratio of 5:1). Mouse SCLC cells were counted and resuspended in T cell medium at 250,000 cells/mL . Tarlatamab was prepared by serial dilution with RPMI from 400nM to make 400nM, 40nM, 4nM, 0.4nM, 0.04nM, 0.004nM. 100 μL of mouse SCLC cells (25,000 cells), 50 μL of Tarlatamab (4x) and 50 μL of T cells were combined and added into a 96 well flat bottom tissue culture treated plate in triplicates. After 48 hours of incubation at 37°C, cells were analyzed for cell viability by flow cytometry. Briefly, cells were collected into 96 well round bottom plates and centrifuge was performed at 1600 rpm for 4 min at room temperature. After discarding supernatant cells were washed with 200 μL of PBS and spin down at 1600 rpm for 4 min at room temperature. Following removal of supernatant, cells were stained with Zombie Violet Fixable Viability Kit (BioLegend, #423113) by resuspending in 50 μL of PBS added with Zombie Violet (1:500) and incubating for 5 minutes in the dark. After washing with FACS buffer (PBS + 2%FBS), cells were stained for human CD45 and incubated at 4°C for 20 minutes. After washing twice with FACS buffer, cells were fixed with 2% PFA and analyzed on MACSQuant Analyzer 10 (Miltenyi Biotec).

#### Immunoblot Analysis

Murine SCLC cells were collected and centrifuged at 1500 rpm for 3 minutes and supernatant was removed. Cell pellet was washed with ice cold PBS and transferred to an Eppendorf tube and centrifuged at 4000 rpm for 4 minutes at 4°C. After carefully removing supernatant, cell pellet was lysed by resuspending in EBC lysis buffer (50 mM Tris-Cl pH 7.5, 250 mM NaCl, 0.5% NP-40, 10% glycerol) supplemented with a protease inhibitor (Complete, Roche Applied Science, #11836153001) and phosphatase inhibitors (PhosSTOP, Sigma, #04906837001) and incubated on a rotator for 40 minutes at 4°C. For murine tumor samples, snap frozen samples were mechanically minced and lysed with EBC lysis buffer similarly. After lysing, tubes were centrifuged at 13200 rpm for 10 minutes at 4°C and the cell extracts were quantified using the Bradford Protein Assay (Bio-Rad Laboratories; #5000006). 20 µg of protein per sample was boiled in sample buffer and resolved by SDS-polyacrylamide gel electrophoresis (SDS-PAGE) (10% SDS-PAGE was used for all proteins) and transferred onto nitrocellulose membranes. The membranes were blocked in 5% milk in Tris-Buffered Saline with 0.1% Tween 20 (TBS-T) for 1 hour at room temperature, and probed with the indicated primary antibodies overnight at 4°C. Membranes were then washed 3 times in TBS-T, probed with the indicated horseradish peroxidase-conjugated (HRP) secondary antibodies for 1 hour at room temperature followed by 3 washes in TBS-T. Bound antibodies were detected with enhanced chemiluminescence (ECL) western blotting detection reagents [Immobilon (Thermo Fisher Scientific, #WBKLS0500) or Supersignal West Pico (Thermo Fisher Scientific, #PI34078)]. Primary antibodies used are, MASH1/Achaete-scute homolog 1 antibody [EPR19840] (Abcam, #ab211327-100ul), DLL3 antibody [EPR22592-18] (Abcam, #ab229902), NEUROD1 antibody [EPR4008] (Abcam, #ab109224), HA tag antibody (Cell Signaling Technology, #3724S), VINCULIN antibody (Sigma Aldrich, #V9131-100UL), and β-Actin−Peroxidase antibody (AC-15, Sigma Aldrich, #A3854-200UL)

#### Pharmacological inhibitors

Tarlatamab was purchased from MedChemExpress (MedChemExpress, #HY-P99575) and stored at -80°C. Pavurutamab was purchased from MedChemExpress (MedChemExpress, AMG-701; # HY-P99814,) and stored at -80 °C.

#### Immunohistochemistry

##### Mouse Tumors

Following fixation, samples were processed on the Leica ASP300S tissue processor and embedded in paraffin (Paraplast X-tra Leica, #39603002). Tissues were sectioned at 5µm and placed on positively charged glass slides (Fisherbrand Superfrost Plus, #22-037-246). H&E staining was performed on the Leica Spectra ST through a standard regressive staining protocol using Harris Hematoxylin (Avantik, #RS4020-8) and Eosin-Y (Anatech, #832). Immunohistochemistry was performed on the Leica Bond III automated staining platform using the Leica Biosystems Refine Detection Kit (Leica, #DS9800). FFPE tissue sections were baked for 30 minutes at 60°C and deparaffinized (Leica, #AR9222) prior to staining. Primary antibodies were visualized via DAB, and counterstained with hematoxylin (Leica, #DS9800). The slides were dehydrated in graded alcohol and coverslipped using the HistoreCore Spectra CV mounting medium (Leica, #3801733). Following antibodies were used at indicated dilutions. Human CD3 (CST, #85061 clone D7A6E Rabbit, 1:800), Mouse CD4 (CST, #25229 clone D7D2Z Rabbit, 1:50), Mouse CD8 (CST, #98941 D4W2Z Rabbit, 1:200).

##### Human Tumors

Immunohistochemistry (IHC) was performed on 4 μm thick sections cut from formalin-fixed paraffin-embedded (FFPE) mouse and human tumor samples. The staining was conducted on the Bond Rx Autostainer (Leica Biosystems) using the Bond Polymer Refine Detection Kit (Leica Biosystems).

Single-marker IHC was performed using antibodies against: ASCL1 (rabbit monoclonal antibody, clone: EPR19840, Abcam; 1:100 concentration, citrate-based antigen retrieval for 30 min.); NEUROD1 (rabbit monoclonal antibody, clone: EPR20766, Abcam; 1:100 concentration, citrate-based antigen retrieval for 30 min.); POU2F3 (rabbit polyclonal antibody, Atlas Antibodies; 1:200 concentration, EDTA-based antigen retrieval for 20 min.); DLL3 (rabbit monoclonal antibody, clone: EPR22592-18, Abcam; 1:100 concentration, EDTA-based antigen retrieval for 20 min.). The slides were counterstained with hematoxylin, dehydrated using graded ethanol and xylene, and mounted with coverslips. The percentage of tumor cells positive for each marker was independently evaluated by two pathologists (BS & YNL), followed by a joint review to reach a final consensus percentage.

#### Multiplex Immunofluorescence Assays

Two optimized multiplex immunofluorescence (mIF) panels were performed on one post-tarlatamab human FFPE tumor tissue sample (Patient 7373). Automated staining was performed on a BOND Rx Autostainer (Leica Biosystems) using the Opal Tyramide Signal Amplification system (Akoya Biosciences).

Following staining, slides were scanned using the PhenoImager HT system (Akoya Biosciences, v2.0.0). We first acquired whole-slide multispectral images, from which specific regions of interest (ROIs) were selected using Phenochart software (v2.0.0, Akoya Biosciences). Then, spectral unmixing was performed in InForm software (v2.6.0, Akoya Biosciences) using spectral libraries generated from monoplex-stained controls.

#### Statistical Analysis

All t-tests were unpaired two-sided, and p values < 0.05 were considered statistically significant unless otherwise specified. No statistical methods were used to predetermine sample size.

Violin plots with box-and-whisker overlays display the median and interquartile range (IQR). For comparisons between two groups, p values were calculated using a two-sided Wilcoxon rank-sum test.

Kaplan-Meier survival curves were generated to estimate survival distributions, and p values were calculated using the two-sided log-rank test.

In all figures, significance levels are denoted as follows: * p < 0.05; ** p < 0.01; *** p < 0.001; **** p < 0.0001. Error bars represent SD unless otherwise indicated in figure legends. The number of independent biological experiments is indicated in each figure legend.

For *in vivo* syngeneic mouse tumor experiments in **Figure 2K-2L** and **Figure 4K**, 2-way ANOVA was used to compare drug treatment vs. vehicle. For *in vivo* syngeneic mouse tumor experiment in **Figure 2J**, statistical significance on Day 18 after initial treatment was calculated using unpaired, two-tailed Students t-test.

For flow cytometry analysis in **Figure. 2C**, **2D**, **2H**, **4B**, **Figure S4A**, **S5D**, at least two biological experiments were performed and representative data are shown. For *in vitro* coculture assay in **Figure 2FG**, representative data from three biological independent experiments are shown. For all *in vivo* experiments, the number of independent biological experiments are indicated in each figure legend. For immunoblots in **Figure 2B**, **4C**, **4L**, **Figure S5F** at least two biological independent experiments were performed, and representative blots are shown. For immunoblots in **Figure 4N** and **Figure S5H**, representative blots from at least two technical replicates are shown.

Analyses were performed in R (v4.5.1) using the following packages: tidyverse (v2.0.0), dplyr (v1.2.0), tidyr (v1.3.2), tibble (v3.3.1), ggplot2 (v4.0.2), ggrepel (v0.9.7), ggpubr (v0.6.3), scales (v1.4.0), viridis (v0.6.5), broom (v1.0.12), randomForest (v4.7-1.2), survival (v3.8-6), survminer (v0.5.2), data.table (v1.18.2.1), ggrastr (v1.0.2), msigdbr (v26.1.0), pheatmap (v1.0.13), scico (v1.5.0), FactoMineR (v2.13), factoextra (v2.0.0), ComplexHeatmap (v2.26.1), circlize (v0.4.17), readxl (v1.4.5), gridExtra (v2.3), BiocManager (v1.30.27), and rlang (v1.1.7).

## ACKNOWLEDGEMENTS

This work was supported by a Novartis DDTRP Award (M.G.O.), Supplemental Novartis DDTRP Award (M.G.O., M.L.F, S.C.B.) award. D.V. is supported by awards from Gustave Roussy, Institut Servier, and Philippe Foundation. This work and S.S. was supported by the Deborah A. Connolly Small Cell Lung Cancer Research Fund. G.S.G. is supported by T32CA009172 from the National Cancer Institute. Y.L. is supported by an NCI K99/R00 Pathway to Independence Award (1K99CA304214-01). We thank members of the Oser, Freedman, and Baca labs for their critical feedback. We are grateful to members of the Translational Immunogenomics Lab at DFCI for scRNA-seq. We thank Dana-Farber/Harvard Cancer Center in Boston, MA, for the use of the Specialized Histopathology Core, which provided histology and immunohistochemistry service. Dana-Farber/Harvard Cancer Center is supported in part by an NCI Cancer Center Support Grant # NIH 5 P30 CA06516.

## AUTHOR CONTRIBUTIONS

D.V. and S.Saito. contributed to conceptualization, methodology, performed and analyzed experiments, data curation, visualization, and writing of manuscript. G.L. and G.G. contributed to conceptualization, methodology, analysis, and data curation. Y.N.L and B.S. optimized and performed all IHC and multiplexed IF. M.L and H.C. coordinated and collected patient samples. Y.L. contributed to analysis. T.W. contributed to validation. J.H.S., H.S., B.J., W.D.K, G.N., and R.N. contributed to experiments. Z.Z., K.S., Z.J., contributed to computational analysis. A.C. contributed to clinical data analysis, K.M. contributed to supervising clinical sample collection and protocol adherence, M.D.S contributed to IHC sample preparation, E.B. and E.S. contributed to experiment design, J.C., M.P, C.C. contributed to project administration, T.K.C. reviewed and edited the manuscript, S.Signoretti. contributed to supervision, resources, J.S. contributed to conceptualization, resources, supervision, project administration, and funding acquisition. S.C.B., M.L.F, and M.G.O. contributed to conceptualization, methodology, investigation, formal analysis, validation, resources, data curation, visualization, supervision, overall project administration, writing manuscript, and funding acquisition.

## DECLARATION OF INTERESTS

M.G.O. reports grants from Eli Lilly, Takeda, Novartis, BMS, Auron Therapeutics, Amgen, and Circle Pharma. G.S.G., D.V., M.G.O., S.C.B., and M.L.F. have patent applications related to methods and for inferring cancer gene expression and their applications (US 63/902,270; US 63/984,890). M.L.F. and S.B.C are co-founders and shareholders of Precede Biosciences. This work was funded by a Novartis DDTRP award and Novartis DDTRP supplemental award. G.S.G. is a co-founder and shareholder of CytoTRACE Biosciences. A.J.C has received honoraria from MJH Life Sciences, Ideology Health, Intellisphere LLC, MedStar Health, PeerDirect and CancerGRACE, and consulting fees from Gilead Sciences, Inc, Daiichi/Astra Zeneca, Novartis, BI, Amgen, and Regeneron. She reports prior research funding to institution from Merck, Monte Rosa, AbbVie, Roche, Daiichi Sankyo, and Amgen (end 08/2025). E.L.S and E.J.B are inventors on a pending patent for CARs targeting TROP2 outside of this submitted work. E.J.B. reports personal fees from ImmuneBridge outside the submitted work. C.C., J.C. are shareholders of Novartis. T.K.C. reports institutional and/or personal, paid and/or unpaid support for research, advisory boards, consultancy, and/or honoraria past 10 years, ongoing or not, from: Alkermes, Arcus Bio, AstraZeneca, Aravive, Aveo, Bayer, Bristol Myers-Squibb, Bicycle Therapeutics, Calithera, Caris, Circle Pharma, Deciphera Pharmaceuticals, Eisai, EMD Serono, Exelixis, GlaxoSmithKline, Gilead, HiberCell, IQVA, Infinity, Institut Servier, Ipsen, Jansen, Kanaph, Lilly, Merck, Nikang, Neomorph, Nuscan/PrecedeBio, Novartis, Oncohost, Pfizer, Roche, Sanofi/Aventis, Scholar Rock, Surface Oncology, Takeda, Tempest, Up-To-Date, CME and non-CME events (Mashup Media, Elite Elements, Peerview, OncLive, MJH, CCO and others), Xencor, outside the submitted work. Institutional patents filed on molecular alterations and immunotherapy response/toxicity, rare genitourinary cancers, and ctDNA/liquids biopsies. Equity: Tempest, Osel, Precede Bio, CureResponse, Primium, Abalytics, Faron Pharma, Open Medicine. Committees: NCCN, GU Steering Committee, ASCO (BOD 6/2024-), ESMO, ACCRU, KidneyCan. Medical writing and editorial assistance support may have been funded by Communications companies in part. No speaker’s bureau. Mentored several non-US citizens on research projects with potential funding (in part) from non-US sources/Foreign Components. The institution (Dana-Farber Cancer Institute) may have received additional independent funding of drug companies or/and royalties potentially involved in research around the subject matter. JS reports consulting from Abbvie, Amgen, AstraZeneca, Boehringer Ingelheim, Curadev, Daiichi Sankyo, Fosun, Genentech, Gilead, GSK, Jazz Pharmaceuticals, Lilly, Merck, Novartis, Pfizer, Pharma Mar, Sanofi; grants from Amgen and Novartis. All other authors declare no competing interests.

## SUPPLEMENTAL FIGURE LEGENDS

**Figure S1. Early Tumor Fraction Dynamics and APEX Transcriptional Profiling in SCLC Plasma Samples.** (**A**) Waterfall plot showing percent change in tumor fraction from baseline to C1D15 (n = 28 patients with paired C1D1 and C1D15 samples; 12 with clinical benefit and 16 without). Only patients with paired baseline and C1D15 plasma samples were included. Bars represent 100 × (C1D15 − Baseline)/Baseline for each patient. Dashed horizontal lines indicate group medians. (**B**) Genome browser view of cfChIP–seq H3K4me3 signal across representative loci (ACTB, SPI1, CHGA, DLL3) in a baseline SCLC patient (6709) and a healthy control. (**C**) Scatter plot showing mean Log_2_TPM APEX expression in 10 healthy controls versus 34 baseline SCLC samples. Grey points indicate genes not included in SCLC or WBC signatures (“Other”), colored points represent genes from the WBC (lavender) or SCLC (purple) gene sets, and black points highlight selected labeled genes. Values are averaged across samples within each group. (**D**) Principal component analysis (PCA) of Log_2_TPM APEX gene expression in 10 healthy controls and 34 baseline SCLC samples. PCA was performed on the merged APEX expression matrix (genes × samples) after scaling. Each point represents a sample (squares, healthy; triangles, SCLC). Points are colored according to tumor fraction using a viridis gradient. Ellipses represent normal data ellipses for each group. Axes indicate the percentage of variance explained by PC1 and PC2. (**E**) Scatter plot showing the association between baseline tumor fraction and mean APEX Log_2_TPM expression of the WBC gene signature in 34 SCLC baseline samples. Dashed line indicates linear regression across all samples. Spearman’s rank correlation coefficient (ρ) and two-sided P value are reported. (**F**) Violin plots showing mean APEX Log_2_TPM expression of the WBC gene signature in 10 healthy controls and 34 baseline SCLC samples ns (non significant). (**G**) Scatter plot showing the association between baseline tumor fraction and mean APEX Log_2_TPM expression of the SCLC gene signature. Spearman’s ρ and two-sided P value are shown. (**H**) Violin plots showing mean APEX Log_2_TPM expression of SCLC gene signatures in 10 healthy controls and 34 baseline SCLC samples **** (p < 0.0001). Plot structure and statistical testing are as described in panel (**F**). (**I**) Volcano plot showing baseline APEX Log2TPM gene expression differences between responders (partial response) (n = 16 patients) and non responders (oligo-progression or progressive disease) (n = 16 patients). The x-axis represents log₂ fold change (responders vs. non responders), and the y-axis represents –log₁₀(p-value). Dashed vertical lines indicate ±1 log₂ fold change, and the dashed horizontal line represents the significance threshold (p = 0.01). Gene labels highlight selected genes of interest in black. Genes enriched in patients with clinical benefit (p < 0.01, unadjusted; log₂ fold change > 1) are shown in green, whereas genes enriched in patients without clinical benefit (p < 0.01, unadjusted; log₂ fold change < −1) are shown in orange.

**Figure S2. Tarlatamab Response is Correlated with SCLC Transcription Factor Subtype.** (**A**) Heatmap of baseline transcriptional subtype probabilities and DLL3 expression using APEX. Columns represent individual patients and rows represent SCLC transcriptional subtype probabilities (SCLC-A, SCLC-I, SCLC-N, and SCLC-P) and DLL3 expression in Log_2_TPM. Subtype probabilities are displayed as relative subtype probabilities, calculated by dividing each subtype probability by the maximum subtype probability within the same sample (range 0–1), highlighting the dominant subtype per patientacross 34 patients. (**B**) Scatter plot showing the correlation between subtype probabilities derived from cfChIP–seq using a Random Forest classifier and probabilities derived from cfMeDIP–seq for each subtype (n = 167 samples) (see **Methods**). The black line represents the linear regression fit. Pearson’s correlation coefficient (R) and two-sided P value are shown on each plot. (**C**) Representative IHC for ASCL1, NEUROD1, POU2F3, and DLL3 from 4 patients. (**D**) IHC quantification for all 6 patients and subtype prediction using cfChIP-seq. Please note all IHC and cfChIP-seq subtype prediction are concordant except for case 6917, which may be due to tissue being obtained six months prior to the plasma sample as this patient received a line of therapy prior to tarlatamab, or to intra-tumor subtype heterogeneity being captured by APEX-derived RNA data but not by the tumor biopsy. (**E**) Performance of DLL3 expression and SCLC-A subtype as predictors of clinical benefit. DLL3 expression was dichotomized at the cohort median Log2TPM value, and subtype classification was derived from the random forest model (SCLC-A vs non-A). Left : 2 × 2 contingency tables were constructed to compare patients with and without clinical benefit between subtypes, and corresponding Fisher’s exact test P-values are shown. Subtypes were grouped as SCLC-A (including Likely-A) vs Others (SCLC-N, Likely-N, and SCLC-P). Right : Bars plots showing positive predictive value (PPV), negative predictive value (NPV), sensitivity, and specificity across baseline samples (n = 34). Proportions are shown within each bar.

**Figure S3. Mechanisms of Acquired Resistance to Tarlatamab in Human SCLC.** (**A**) Longitudinal plot showing mean APEX Log_2_TPM expression of SCLC and WBC signature scores across serial timepoints for patient 6940, alongside tumor fraction (black dotted line; dual y-axis). Dashed vertical lines indicate radiographic assessments with clinical outcomes annotated as: PR (partial response), SD (stable disease), PD (progressive disease), and Oligo-PD (oligoprogression). (**B**) Longitudinal dynamics of SCLC-N and SCLC-A transcriptional signatures for patient 6940. Thick lines represent median expression per timepoint. Shaded ribbons indicate IQR. Differences between signatures at each timepoint were assessed using two-sided Wilcoxon rank-sum tests and annotated as * (p < 0.05), ** (p < 0.01), or ns (not significant). (**C**) Haplotype-based BAF analysis framework. (1) Germline SNPs are phased and haplotype-specific read counts are used to estimate allele fractions across phased blocks. (2) Observed BAF is calculated across genomic segments by aggregating haplotype fractions. (3) Expected BAF values are derived from allele-specific copy-number configurations and tumor fraction. (4) Deviations between observed and expected BAF indicate clonal heterogeneity, whereas concordance supports a single dominant clone. (**D**) Haplotype-based BAF analysis in patient 6527. Observed versus expected BAF across genomic segments at baseline and progression. Black points indicate segments consistent with a single-clone model; red points indicate deviations. Chromosome 15 deviates from the model at baseline (a dominant clone with two B-alleles and no A-Alleles, and a minor subclone with an A-allele) but is consistent with a dominant 3:1 allelic configuration at progression. Lower panels show copy-number state and haplotype-based BAF distributions for chromosome 15, with broader and mismatched distributions at baseline and tighter agreement at progression, consistent with selection of a pre-existing subclone. Average depth of sequencing was 46x and 28x for C1D1 vs C9D1 for patient 6527. (**E**) Clonal evolution model for chromosome 15. Baseline tumor is composed of a dominant B-allele–only clone and a minor A-containing subclone. At progression, the dominant clone carries an A allele. Scenarios involving de novo re-emergence of the A allele are unlikely, whereas expansion of a pre-existing A-containing subclone is consistent with the observed data, supporting clonal selection under treatment. (**F**) Same as (**D**), but for patient 5685. Average depth of sequencing was 49x and 45x for C1D1 and C14D15 for patient 5685. (**G**) scRNAseq UMAP visualization from adrenalectomy for patient 7373. n=1813 cells. (**H**) scRNAseq UMAP visualization of tumor cells markers (ASCL1, NEUROD1, DLL3, and INSM1) from adrenalectomy for patient 7373. (**I**) Representative immunohistochemistry (IHC) images from the adrenalectomy specimen of patient 7373.

**Figure S4. Conserved Acquired Resistance to Tarlatamab in Mice Through NEUROD1-Driven DLL3 Loss.** (**A**) Quantitation of mean fluorescence intensity (MFI) from flow cytometry analysis for cell-surface DLL3 expression in the parental 1014P2 and 1014P2-TR#2 cell lines. N= 3 independant experiments. *p < 0.01, Student’s unpaired two-tailed t test. (**B**) Full image of IHC staining for ASCL1, NEUROD1, and DLL3 in 1014P2 vehicle treated tumor (top row) and 1014P2-TR#2 tumor (bottom row) shown in **Figure 4D**. (**C**) Heatmap of top differentially expressed genes (p value<0.05) from RNA-seq data comparing 1014P2-TR#2 tumor vs. 2 independent vehicle-treated 1014P2 tumors (total 1676 genes). Neurod1 and EMT genes are labeled. (**D**) Hallmark gene set enrichment analysis of RNA-seq data from **Figure S4C** of top enriched gene sets in the 1014P2-TR#2 tumor vs. 2 independent vehicle-treated 1014P2 tumors. NES = Normalized Enrichment Score. (**E-G**) FPKM value of Neurod1 (**E**), Ascl1 (**F**), and Dll3 (**G**) acquired from RNA-seq data on left and RT-qPCR validation normalized to 1014P2 on right. (**H**) IGV tracks for each indicated gene obtained from H3K4me3 (green), H3K27ac (maroon), H3K36me3 (blue) ChIP-seq from 1014P2 and 1014P2-TR#2 cells. Bottom gray shows input. Scale is shown on left of each track. Right panel shows zoomed scale for Neurod1. (**I**) Venn diagram showing the overlap of upregulated genes in tarlatamab acquired-resistance tumor that exhibited gain of NEUROD1 as a resistance mechanism in mouse (1014P2-TR#2) and human sample (patient 6527). Differentially upregulated genes in 1014P2-TR#2 tumors vs. vehicle-treated 1014P2 tumors (adjusted p value under 0.05, total 498 genes, green) are compared with genes upregulated (>1.5-fold) at progression (1 month after last C8D15 tarlatamab dose) relative to baseline in patient 6527 (total 1167 genes, blue). Genes in solid black box highlight genes found in HALLMARK_EPITHELIAL_MESENCHYMAL_TRANSITION human gene set (M5930). Broken black box highlight genes found in the NEUROD1 targeted genes list reported by Borromeo et al.^9^. (**J**) Flow cytometric plots of tumors harvested at the experimental endpoint from the study shown in **Figure 4H** and not shown in **Figure 4I.** GFP+ tumor cells and infiltrating immune cells (CD45⁺, CD3⁺, CD4⁺, and CD8⁺) are shown. 1014P2 tumors on left and 1014P2-TR#2 tumors on right. Top two rows are gated of live cells, and bottom row is gated on live/GFP-CD45+ cells. (**K**) CD45, CD8, CD4 infiltration in 1014P2 derived and 1014P2-TR#2 derived tumor from flow cytometry data obtained in **Figure 4I** and **Figure 4J.** n=2 for each condition. (**L**) H&E and CD4 IHC staining of the tumor shown in **Figure 4K**. The black box indicates the region magnified in the lower right corner of each image. Scale bars: 100 μm (low magnification, original 5x) and 25 μm (higher magnification inset, original 40x). (**M**) Histological analysis of the other tumor from **Figure 4H** showing the entire H&E and IHC panel of CD3, CD4, CD8. The black box indicates the region magnified in the lower right corner of each image. Scale bars: 100 μm (low magnification, original 5x) and 25 μm (higher magnification inset, original 40x).

**Figure S5. NEUROD1 Promotes DLL3 Downregulation Under Tarlatamab Selective Pressure *In Vivo.*** (**A**) Long term treatment with tarlatamab 0.3mg/kg (green arrowhead) and follow up of 1014P2 parental tumors shown in **Figure 4K**. n=8 mice 16 tumors show no evidence of recurrence after 100 days post first tarlatamab treatment. (**B**) Humanized CD3 mice which rejected 1014P2 after 3 doses of tarlatamab treatment from **Figure 2J-2M** and followed up for over 150 days to confirm no evidence of recurrence were rechallenged with 1014P2 (8million cells, subcutaneous injection) on both flanks and tumor growth was monitored. n = 7 mice, 14 tumors. 4 tumors were rejected and 10 tumors from five mice were established. (**C**) Humanized CD3 mice rechallenged with 1014P2 and established tumors in **Figure S5B** were treated with weekly tarlatamab 0.3mg/kg (green arrowhead) and tumor growth was monitored. n = 10 tumors total. 7 tumors were completely rejected and 3 tumors showed progression after partial response. 1 tumor showed recurrence after complete rejection. Details of recurrence tumors are described in methods. (**D**) Representative flow cytometric histogram for cell-surface DLL3 expression in the parental 1014P2 and tarlatamab resistant cell lines compared to isotype control. y-axis is normalized to mode. (**E**) Percentage of TIM3 positive CD8 T cells, LAG3 positive CD8 T cells and FOXP3 positive CD4 T cell infiltration in 1014P2 derived and 1014P2-TR#2 derived tumor from flow cytometry analysis. (**F**) 1014P2 cells were transuduced with NEUROD1 over expression vector (1014P2 NEUROD1-OE) and immunoblot analysis was performed for each indicated antibody. (**G**) 1014P2 cells transduced with NEUROD1 over expression vector were subcutaneously inoculated into both flanks of humanized CD3 mice (genOway). After at least one side of the tumor exceeded 200mm^3^, mice were randomized to either vehicle (PBS) treatment or weekly tarlatamab 0.1mg/kg treatment (green arrowhead) and tumor size was monitored. Tumors were excluded if there was no measurable tumor on the day of treatment initiation. Vehicle n = 6 tumors (three mice), tarlatamab n = 7 tumors (four mice). (**H**) Immunoblot analysis of 1014P2 cells transduced with NEUROD1 overexpression vector and then inoculated into humanized CD3 mice, treated with either vehicle or tarlatamab, and finally harvested at endpoint for protein extraction (1014P2 NEUROD1-OE tumors). PBS treated 1014P2 tumors (PBS) and BCMA targeting bispecific T cell engager Pavurutamab treated 1014P2 tumors (BCMA) are shown as controls.

## TABLE LEGEND

**Table 1.** Summary of Baseline Demographic and Clinical Characteristics of Patient Cohort.

## References

1. Rudin, C.M., Brambilla, E., Faivre-Finn, C., and Sage, J. (2021). Small-cell lung cancer. Nat Rev Dis Primers 7, 3. 10.1038/s41572-020-00235-0.

2. Ahn, M.J., Cho, B.C., Felip, E., Korantzis, I., Ohashi, K., Majem, M., Juan-Vidal, O., Handzhiev, S., Izumi, H., Lee, J.S., et al. (2023). Tarlatamab for Patients with Previously Treated Small-Cell Lung Cancer. N Engl J Med 389, 2063–2075. 10.1056/NEJMoa2307980.

3. Mountzios, G., Sun, L., Cho, B.C., Demirci, U., Baka, S., Gumus, M., Lugini, A., Zhu, B., Yu, Y., Korantzis, I., et al. (2025). Tarlatamab in Small-Cell Lung Cancer after Platinum-Based Chemotherapy. N Engl J Med 393, 349–361. 10.1056/NEJMoa2502099.

4. Paz-Ares, L., Champiat, S., Lai, W.V., Izumi, H., Govindan, R., Boyer, M., Hummel, H.D., Borghaei, H., Johnson, M.L., Steeghs, N., et al. (2023). Tarlatamab, a First-in-Class DLL3-Targeted Bispecific T-Cell Engager, in Recurrent Small-Cell Lung Cancer: An Open-Label, Phase I Study. J Clin Oncol 41, 2893–2903. 10.1200/JCO.22.02823.

5. Dunwoodie, S.L., Henrique, D., Harrison, S.M., and Beddington, R.S. (1997). Mouse Dll3: a novel divergent Delta gene which may complement the function of other Delta homologues during early pattern formation in the mouse embryo. Development 124, 3065–3076. 10.1242/dev.124.16.3065.

6. Geffers, I., Serth, K., Chapman, G., Jaekel, R., Schuster-Gossler, K., Cordes, R., Sparrow, D.B., Kremmer, E., Dunwoodie, S.L., Klein, T., and Gossler, A. (2007). Divergent functions and distinct localization of the Notch ligands DLL1 and DLL3 in vivo. J Cell Biol 178, 465–476. 10.1083/jcb.200702009.

7. Ladi, E., Nichols, J.T., Ge, W., Miyamoto, A., Yao, C., Yang, L.T., Boulter, J., Sun, Y.E., Kintner, C., and Weinmaster, G. (2005). The divergent DSL ligand Dll3 does not activate Notch signaling but cell autonomously attenuates signaling induced by other DSL ligands. J Cell Biol 170, 983–992. 10.1083/jcb.200503113.

8. Kim, J.W., Ko, J.H., and Sage, J. (2022). DLL3 regulates Notch signaling in small cell lung cancer. iScience 25, 105603. 10.1016/j.isci.2022.105603.

9. Borromeo, M.D., Savage, T.K., Kollipara, R.K., He, M., Augustyn, A., Osborne, J.K., Girard, L., Minna, J.D., Gazdar, A.F., Cobb, M.H., and Johnson, J.E. (2016). ASCL1 and NEUROD1 Reveal Heterogeneity in Pulmonary Neuroendocrine Tumors and Regulate Distinct Genetic Programs. Cell Rep 16, 1259–1272. 10.1016/j.celrep.2016.06.081.

10. Henke, R.M., Meredith, D.M., Borromeo, M.D., Savage, T.K., and Johnson, J.E. (2009). Ascl1 and Neurog2 form novel complexes and regulate Delta-like3 (Dll3) expression in the neural tube. Dev Biol 328, 529–540. 10.1016/j.ydbio.2009.01.007.

11. Saunders, L.R., Bankovich, A.J., Anderson, W.C., Aujay, M.A., Bheddah, S., Black, K., Desai, R., Escarpe, P.A., Hampl, J., Laysang, A., et al. (2015). A DLL3-targeted antibody-drug conjugate eradicates high-grade pulmonary neuroendocrine tumor-initiating cells in vivo. Sci Transl Med 7, 302ra136. 10.1126/scitranslmed.aac9459.

12. Rudin, C.M., Reck, M., Johnson, M.L., Blackhall, F., Hann, C.L., Yang, J.C., Bailis, J.M., Bebb, G., Goldrick, A., Umejiego, J., and Paz-Ares, L. (2023). Emerging therapies targeting the delta-like ligand 3 (DLL3) in small cell lung cancer. J Hematol Oncol 16, 66. 10.1186/s13045-023-01464-y.

13. Gay, C.M., Stewart, C.A., Park, E.M., Diao, L., Groves, S.M., Heeke, S., Nabet, B.Y., Fujimoto, J., Solis, L.M., Lu, W., et al. (2021). Patterns of transcription factor programs and immune pathway activation define four major subtypes of SCLC with distinct therapeutic vulnerabilities. Cancer Cell 39, 346–360 e347. 10.1016/j.ccell.2020.12.014.

14. Rudin, C.M., Poirier, J.T., Byers, L.A., Dive, C., Dowlati, A., George, J., Heymach, J.V., Johnson, J.E., Lehman, J.M., MacPherson, D., et al. (2019). Molecular subtypes of small cell lung cancer: a synthesis of human and mouse model data. Nat Rev Cancer 19, 289–297. 10.1038/s41568-019-0133-9.

15. Baca, S.C., Seo, J.H., Davidsohn, M.P., Fortunato, B., Semaan, K., Sotudian, S., Lakshminarayanan, G., Diossy, M., Qiu, X., El Zarif, T., et al. (2023). Liquid biopsy epigenomic profiling for cancer subtyping. Nat Med 29, 2737–2741. 10.1038/s41591-023-02605-z.

16. El Zarif, T., Meador, C.B., Qiu, X., Seo, J.H., Davidsohn, M.P., Savignano, H., Lakshminarayanan, G., McClure, H.M., Canniff, J., Fortunato, B., et al. (2024). Detecting Small Cell Transformation in Patients with Advanced EGFR Mutant Lung Adenocarcinoma through Epigenomic cfDNA Profiling. Clin Cancer Res 30, 3798–3811. 10.1158/1078-0432.CCR-24-0466.

17. Heeke, S., Gay, C.M., Estecio, M.R., Tran, H., Morris, B.B., Zhang, B., Tang, X., Raso, M.G., Rocha, P., Lai, S., et al. (2024). Tumor- and circulating-free DNA methylation identifies clinically relevant small cell lung cancer subtypes. Cancer Cell 42, 225–237 e225. 10.1016/j.ccell.2024.01.001.

18. Hiatt, J.B., Doebley, A.L., Arnold, H.U., Adil, M., Sandborg, H., Persse, T.W., Ko, M., Wu, F., Quintanal Villalonga, A., Santana-Davila, R., et al. (2024). Molecular phenotyping of small cell lung cancer using targeted cfDNA profiling of transcriptional regulatory regions. Sci Adv 10, eadk2082. 10.1126/sciadv.adk2082.

19. Takahashi, N., Pongor, L., Agrawal, S.P., Shtumpf, M., Gurjar, A., Rajapakse, V.N., Shafiei, A., Schultz, C.W., Kim, S., Roame, D., et al. (2025). Genomic alterations and transcriptional phenotypes in circulating free DNA and matched metastatic tumor. Genome Med 17, 15. 10.1186/s13073-025-01438-4.

20. Gulati, G.S., Vasseur, D., Nawfal, R., Sotudian, S., Semaan, K., Eid, M., Seo, J.-H., Phillips, N., Canniff, J.J., Savignano, H., et al. (2026). Integrated inference of cancer gene expression from cell-free plasma chromatin. bioRxiv, 2026.2002.2018.706026. 10.64898/2026.02.18.706026.

21. Maansson, C.T., Thomsen, L.S., Meldgaard, P., Nielsen, A.L., and Sorensen, B.S. (2024). Integration of Cell-Free DNA End Motifs and Fragment Lengths Can Identify Active Genes in Liquid Biopsies. Int J Mol Sci 25. 10.3390/ijms25021243.

22. Trier Maansson, C., Meldgaard, P., Stougaard, M., Nielsen, A.L., and Sorensen, B.S. (2023). Cell-free chromatin immunoprecipitation can determine tumor gene expression in lung cancer patients. Mol Oncol 17, 722–736. 10.1002/1878-0261.13394.

23. Sadeh, R., Sharkia, I., Fialkoff, G., Rahat, A., Gutin, J., Chappleboim, A., Nitzan, M., Fox-Fisher, I., Neiman, D., Meler, G., et al. (2021). ChIP-seq of plasma cell-free nucleosomes identifies gene expression programs of the cells of origin. Nat Biotechnol 39, 586–598. 10.1038/s41587-020-00775-6.

24. Snyder, M.W., Kircher, M., Hill, A.J., Daza, R.M., and Shendure, J. (2016). Cell-free DNA Comprises an In Vivo Nucleosome Footprint that Informs Its Tissues-Of-Origin. Cell 164, 57–68. 10.1016/j.cell.2015.11.050.

25. Ulz, P., Thallinger, G.G., Auer, M., Graf, R., Kashofer, K., Jahn, S.W., Abete, L., Pristauz, G., Petru, E., Geigl, J.B., et al. (2016). Inferring expressed genes by whole-genome sequencing of plasma DNA. Nat Genet 48, 1273–1278. 10.1038/ng.3648.

26. Esfahani, M.S., Hamilton, E.G., Mehrmohamadi, M., Nabet, B.Y., Alig, S.K., King, D.A., Steen, C.B., Macaulay, C.W., Schultz, A., Nesselbush, M.C., et al. (2022). Inferring gene expression from cell-free DNA fragmentation profiles. Nat Biotechnol 40, 585–597. 10.1038/s41587-022-01222-4.

27. Zhao, Z., Szczepanski, A.P., Tsuboyama, N., Abdala-Valencia, H., Goo, Y.A., Singer, B.D., Bartom, E.T., Yue, F., and Wang, L. (2021). PAX9 Determines Epigenetic State Transition and Cell Fate in Cancer. Cancer Res 81, 4696–4708. 10.1158/0008-5472.CAN-21-1114.

28. Ajay, A., Wang, H., Rezvani, A., Savari, O., Grubb, B.J., McColl, K.S., Yoon, S., Joseph, P.L., Kopp, S.R., Kresak, A.M., et al. (2025). Assessment of targets of antibody drug conjugates in SCLC. NPJ Precis Oncol 9, 1. 10.1038/s41698-024-00784-7.

29. Gezelius, E., Rekhtman, N., Baine, M.K., Rudin, C.M., Drilon, A., and Cooper, A.J. (2025). Seizure-Related Homolog Protein 6 (SEZ6): Biology and Therapeutic Target in Neuroendocrine Carcinomas. Clin Cancer Res 31, 4419–4428. 10.1158/1078-0432.CCR-25-2090.

30. Duplaquet, L., So, K., Ying, A.W., Pal Choudhuri, S., Li, X., Xu, G.D., Li, Y., Qiu, X., Li, R., Singh, S., et al. (2024). Mammalian SWI/SNF complex activity regulates POU2F3 and constitutes a targetable dependency in small cell lung cancer. Cancer Cell 42, 1352–1369 e1313. 10.1016/j.ccell.2024.06.012.

31. Huang, Y.H., Klingbeil, O., He, X.Y., Wu, X.S., Arun, G., Lu, B., Somerville, T.D.D., Milazzo, J.P., Wilkinson, J.E., Demerdash, O.E., et al. (2018). POU2F3 is a master regulator of a tuft cell-like variant of small cell lung cancer. Genes Dev 32, 915–928. 10.1101/gad.314815.118.

32. Lang, C., Megyesfalvi, Z., Lantos, A., Oberndorfer, F., Hoda, M.A., Solta, A., Ferencz, B., Fillinger, J., Solyom-Tisza, A., Querner, A.S., et al. (2024). C-Myc protein expression indicates unfavorable clinical outcome in surgically resected small cell lung cancer. World J Surg Oncol 22, 57. 10.1186/s12957-024-03315-7.

33. Nabet, B.Y., Hamidi, H., Lee, M.C., Banchereau, R., Morris, S., Adler, L., Gayevskiy, V., Elhossiny, A.M., Srivastava, M.K., Patil, N.S., et al. (2024). Immune heterogeneity in small-cell lung cancer and vulnerability to immune checkpoint blockade. Cancer Cell 42, 429–443 e424. 10.1016/j.ccell.2024.01.010.

34. Oser, M.G., MacPherson, D., Oliver, T.G., Sage, J., and Park, K.S. (2024). Genetically-engineered mouse models of small cell lung cancer: the next generation. Oncogene 43, 457–469. 10.1038/s41388-023-02929-7.

35. Hong, D., Knelson, E.H., Li, Y., Durmaz, Y.T., Gao, W., Walton, E., Vajdi, A., Thai, T., Sticco-Ivins, M., Sabet, A.H., et al. (2021). Plasticity in the Absence of NOTCH Uncovers a RUNX2-Dependent Pathway in Small Cell Lung Cancer. Cancer Res. 10.1158/0008-5472.CAN-21-1991.

36. Oser, M.G., Sabet, A.H., Gao, W., Chakraborty, A.A., Schinzel, A.C., Jennings, R.B., Fonseca, R., Bonal, D.M., Booker, M.A., Flaifel, A., et al. (2019). The KDM5A/RBP2 histone demethylase represses NOTCH signaling to sustain neuroendocrine differentiation and promote small cell lung cancer tumorigenesis. Genes Dev 33, 1718–1738. 10.1101/gad.328336.119.

37. Li, Y., Mahadevan, N.R., Duplaquet, L., Hong, D., Durmaz, Y.T., Jones, K.L., Cho, H., Morrow, M., Protti, A., Poitras, M.J., et al. (2023). Aurora A kinase inhibition induces accumulation of SCLC tumor cells in mitosis with restored interferon signaling to increase response to PD-L1. Cell Rep Med 4, 101282. 10.1016/j.xcrm.2023.101282.

38. Mahadevan, N.R., Knelson, E.H., Wolff, J.O., Vajdi, A., Saigi, M., Campisi, M., Hong, D., Thai, T.C., Piel, B., Han, S., et al. (2021). Intrinsic immunogenicity of small cell lung carcinoma revealed by its cellular plasticity. Cancer Discov. 10.1158/2159-8290.CD-20-0913.

39. Zhang, H., Christensen, C.L., Dries, R., Oser, M.G., Deng, J., Diskin, B., Li, F., Pan, Y., Zhang, X., Yin, Y., et al. (2019). CDK7 Inhibition Potentiates Genome Instability Triggering Anti-tumor Immunity in Small Cell Lung Cancer. Cancer Cell. 10.1016/j.ccell.2019.11.003.

40. Giffin, M.J., Cooke, K., Lobenhofer, E.K., Estrada, J., Zhan, J., Deegen, P., Thomas, M., Murawsky, C.M., Werner, J., Liu, S., et al. (2021). AMG 757, a Half-Life Extended, DLL3-Targeted Bispecific T-Cell Engager, Shows High Potency and Sensitivity in Preclinical Models of Small-Cell Lung Cancer. Clin Cancer Res 27, 1526–1537. 10.1158/1078-0432.CCR-20-2845.

41. Cai, L., Liu, H., Huang, F., Fujimoto, J., Girard, L., Chen, J., Li, Y., Zhang, Y.A., Deb, D., Stastny, V., et al. (2021). Cell-autonomous immune gene expression is repressed in pulmonary neuroendocrine cells and small cell lung cancer. Commun Biol 4, 314. 10.1038/s42003-021-01842-7.

42. Chan, J.M., Quintanal-Villalonga, A., Gao, V.R., Xie, Y., Allaj, V., Chaudhary, O., Masilionis, I., Egger, J., Chow, A., Walle, T., et al. (2021). Signatures of plasticity, metastasis, and immunosuppression in an atlas of human small cell lung cancer. Cancer Cell 39, 1479–1496 e1418. 10.1016/j.ccell.2021.09.008.

43. Horn, L., Mansfield, A.S., Szczesna, A., Havel, L., Krzakowski, M., Hochmair, M.J., Huemer, F., Losonczy, G., Johnson, M.L., Nishio, M., et al. (2018). First-Line Atezolizumab plus Chemotherapy in Extensive-Stage Small-Cell Lung Cancer. N Engl J Med. 10.1056/NEJMoa1809064.

44. Paz-Ares, L., Dvorkin, M., Chen, Y., Reinmuth, N., Hotta, K., Trukhin, D., Statsenko, G., Hochmair, M.J., Ozguroglu, M., Ji, J.H., et al. (2019). Durvalumab plus platinum-etoposide versus platinum-etoposide in first-line treatment of extensive-stage small-cell lung cancer (CASPIAN): a randomised, controlled, open-label, phase 3 trial. Lancet 394, 1929–1939. 10.1016/S0140-6736(19)32222-6.

45. Reck, M., Dziadziuszko, R., Sugawara, S., Kao, S., Hochmair, M., Huemer, F., de Castro, G., Jr., Havel, L., Bernabe Caro, R., Losonczy, G., et al. (2024). Five-year survival in patients with extensive-stage small cell lung cancer treated with atezolizumab in the Phase III IMpower133 study and the Phase III IMbrella A extension study. Lung Cancer 196, 107924. 10.1016/j.lungcan.2024.107924.

46. Mishra, A., Meador, C.B., Kikkeri, K., Cunneely, Q., Lin, M., Carmona-LaSalle, T.J., Huang, S.B., Bell, R., Putaturo, V., Xia, W., et al. (2026). Circulating Tumor Cells Predict Response to the DLL3-targeting Bispecific Antibody Tarlatamab. Cancer Discov. 10.1158/2159-8290.CD-25-1483.

47. Baine, M.K., Hsieh, M.S., Lai, W.V., Egger, J.V., Jungbluth, A.A., Daneshbod, Y., Beras, A., Spencer, R., Lopardo, J., Bodd, F., et al. (2020). SCLC Subtypes Defined by ASCL1, NEUROD1, POU2F3, and YAP1: A Comprehensive Immunohistochemical and Histopathologic Characterization. J Thorac Oncol 15, 1823–1835. 10.1016/j.jtho.2020.09.009.

48. Ireland, A.S., Micinski, A.M., Kastner, D.W., Guo, B., Wait, S.J., Spainhower, K.B., Conley, C.C., Chen, O.S., Guthrie, M.R., Soltero, D., et al. (2020). MYC Drives Temporal Evolution of Small Cell Lung Cancer Subtypes by Reprogramming Neuroendocrine Fate. Cancer Cell 38, 60–78 e12. 10.1016/j.ccell.2020.05.001.

49. Lissa, D., Takahashi, N., Desai, P., Manukyan, I., Schultz, C.W., Rajapakse, V., Velez, M.J., Mulford, D., Roper, N., Nichols, S., et al. (2022). Heterogeneity of neuroendocrine transcriptional states in metastatic small cell lung cancers and patient-derived models. Nat Commun 13, 2023. 10.1038/s41467-022-29517-9.

50. Mollaoglu, G., Guthrie, M.R., Bohm, S., Bragelmann, J., Can, I., Ballieu, P.M., Marx, A., George, J., Heinen, C., Chalishazar, M.D., et al. (2017). MYC Drives Progression of Small Cell Lung Cancer to a Variant Neuroendocrine Subtype with Vulnerability to Aurora Kinase Inhibition. Cancer Cell 31, 270–285. 10.1016/j.ccell.2016.12.005.

51. Duplaquet, L., Li, Y., Booker, M.A., Xie, Y., Olsen, S.N., Patel, R.A., Hong, D., Hatton, C., Denize, T., Walton, E., et al. (2023). KDM6A epigenetically regulates subtype plasticity in small cell lung cancer. Nat Cell Biol. 10.1038/s41556-023-01210-z.

52. Ireland, A.S., Xie, D.A., Hawgood, S.B., Barbier, M.W., Zuo, L.Y., Hanna, B.E., Lucas-Randolph, S., Tyson, D.R., Witt, B.L., Govindan, R., et al. (2025). Basal cell of origin resolves neuroendocrine-tuft lineage plasticity in cancer. Nature. 10.1038/s41586-025-09503-z.

53. Cortes-Selva, D., Perova, T., Skerget, S., Vishwamitra, D., Stein, S., Boominathan, R., Lau, O., Calara-Nielsen, K., Davis, C., Patel, J., et al. (2024). Correlation of immune fitness with response to teclistamab in relapsed/refractory multiple myeloma in the MajesTEC-1 study. Blood 144, 615–628. 10.1182/blood.2023022823.

54. Graeme Fenton 1, A.L.Z.,, S.Z.,, 4, C.T.,, 4, M.S.,, 4, A.S.,, 4, R.M., et al. (2025). Brief Report: Multi-institution real-world analysis evaluating safety, efficacy and ctDNA dynamics following tarlatamab in patients with previously treated SCLC and LCNEC. JTO 10.1016/j.jtocrr.2025.100933).

55. Ciardullo, C., Tobalina, L., Carr, T.H., Szekeres, P., Kraljevic, S., Byers, L.A., and Fabbri, G. (2025). Early ctDNA Dynamics Inform First-Line Therapy in Patients with Extensive-Stage Small Cell Lung Cancer. Clin Cancer Res 31, 4457–4462. 10.1158/1078-0432.CCR-25-2011.

56. He, T., Xiao, L., Qiao, Y., Klingbeil, O., Young, E., Wu, X.S., Mannan, R., Mahapatra, S., Redin, E., Cho, H., et al. (2024). Targeting the mSWI/SNF complex in POU2F-POU2AF transcription factor-driven malignancies. Cancer Cell 42, 1336–1351 e1339. 10.1016/j.ccell.2024.06.006.

57. Szczepanski, A., Tsuboyama, N., Lyu, H., Wang, P., Beytullahoglu, O., Zhang, T., Singer, B.D., Yue, F., Zhao, Z., and Wang, L. (2024). A SWI/SNF-dependent transcriptional regulation mediated by POU2AF2/C11orf53 at enhancer. Nat Commun 15, 2067. 10.1038/s41467-024-46492-5.

58. Wu, X.S., He, X.Y., Ipsaro, J.J., Huang, Y.H., Preall, J.B., Ng, D., Shue, Y.T., Sage, J., Egeblad, M., Joshua-Tor, L., and Vakoc, C.R. (2022). OCA-T1 and OCA-T2 are coactivators of POU2F3 in the tuft cell lineage. Nature 607, 169–175. 10.1038/s41586-022-04842-7.

59. Shen, S.Y., Burgener, J.M., Bratman, S.V., and De Carvalho, D.D. (2019). Preparation of cfMeDIP-seq libraries for methylome profiling of plasma cell-free DNA. Nat Protoc 14, 2749–2780. 10.1038/s41596-019-0202-2.

60. Adalsteinsson, V.A., Ha, G., Freeman, S.S., Choudhury, A.D., Stover, D.G., Parsons, H.A., Gydush, G., Reed, S.C., Rotem, D., Rhoades, J., et al. (2017). Scalable whole-exome sequencing of cell-free DNA reveals high concordance with metastatic tumors. Nat Commun 8, 1324. 10.1038/s41467-017-00965-y.

61. Zhang, Z., Da Silva Cordeiro, P., Chhetri, S.B., Fortunato, B., Jin, Z., Chehade, R.E.H., Semaan, K., Gulati, G., Lee, G.G., Hemauer, C., et al. (2025). SNAP: Streamlined Nextflow Analysis Pipeline for Immunoprecipitation-Based Epigenomic Profiling of Circulating Chromatin. bioRxiv. 10.64898/2025.12.30.694452.

62. Martin, M. (2011). Cutadapt removes adapter sequences from high-throughput sequencing reads. *EMBnet.journal, [S.l.], v. 17, n. 1, p. pp. 10-12, may 2011. ISSN 2226-6089*. .

63. Li, H., and Durbin, R. (2009). Fast and accurate short read alignment with Burrows-Wheeler transform. Bioinformatics 25, 1754–1760. 10.1093/bioinformatics/btp324.

64. Quinlan, A.R., and Hall, I.M. (2010). BEDTools: a flexible suite of utilities for comparing genomic features. Bioinformatics 26, 841–842. 10.1093/bioinformatics/btq033.

65. Lawrence, M., Huber, W., Pages, H., Aboyoun, P., Carlson, M., Gentleman, R., Morgan, M.T., and Carey, V.J. (2013). Software for computing and annotating genomic ranges. PLoS Comput Biol 9, e1003118. 10.1371/journal.pcbi.1003118.

66. Chemi, F., Pearce, S.P., Clipson, A., Hill, S.M., Conway, A.M., Richardson, S.A., Kamieniecka, K., Caeser, R., White, D.J., Mohan, S., et al. (2022). cfDNA methylome profiling for detection and subtyping of small cell lung cancers. Nat Cancer 3, 1260–1270. 10.1038/s43018-022-00415-9.

67. Zhang, Y., Liu, T., Meyer, C.A., Eeckhoute, J., Johnson, D.S., Bernstein, B.E., Nusbaum, C., Myers, R.M., Brown, M., Li, W., and Liu, X.S. (2008). Model-based analysis of ChIP-Seq (MACS). Genome Biol 9, R137. 10.1186/gb-2008-9-9-r137.

68. Auwera, G.v.d., and O’Connor, B.D. (2020). Genomics in the cloud : using Docker, GATK, and WDL in Terra, First edition. Edition (O’Reilly Media).

69. Danecek, P., Bonfield, J.K., Liddle, J., Marshall, J., Ohan, V., Pollard, M.O., Whitwham, A., Keane, T., McCarthy, S.A., Davies, R.M., and Li, H. (2021). Twelve years of SAMtools and BCFtools. Gigascience 10. 10.1093/gigascience/giab008.

70. Loh, P.R., Danecek, P., Palamara, P.F., Fuchsberger, C., Y, A.R., H, K.F., Schoenherr, S., Forer, L., McCarthy, S., Abecasis, G.R., et al. (2016). Reference-based phasing using the Haplotype Reference Consortium panel. Nat Genet 48, 1443–1448. 10.1038/ng.3679.

71. Genomes Project, C., Auton, A., Brooks, L.D., Durbin, R.M., Garrison, E.P., Kang, H.M., Korbel, J.O., Marchini, J.L., McCarthy, S., McVean, G.A., and Abecasis, G.R. (2015). A global reference for human genetic variation. Nature 526, 68–74. 10.1038/nature15393.

72. Martin, M., Ebert, P., and Marschall, T. (2023). Read-Based Phasing and Analysis of Phased Variants with WhatsHap. Methods Mol Biol 2590, 127–138. 10.1007/978-1-0716-2819-5_8.

73. Ha, G., Roth, A., Khattra, J., Ho, J., Yap, D., Prentice, L.M., Melnyk, N., McPherson, A., Bashashati, A., Laks, E., et al. (2014). TITAN: inference of copy number architectures in clonal cell populations from tumor whole-genome sequence data. Genome Res 24, 1881–1893. 10.1101/gr.180281.114.

74. Breiman, L. (2001). Random Forests. Machine Learning 45, 5–32. 10.1023/A:1010933404324.

75. Gavish, A., Tyler, M., Greenwald, A.C., Hoefflin, R., Simkin, D., Tschernichovsky, R., Galili Darnell, N., Somech, E., Barbolin, C., Antman, T., et al. (2023). Hallmarks of transcriptional intratumour heterogeneity across a thousand tumours. Nature 618, 598–606. 10.1038/s41586-023-06130-4.

76. Harrow, J., Frankish, A., Gonzalez, J.M., Tapanari, E., Diekhans, M., Kokocinski, F., Aken, B.L., Barrell, D., Zadissa, A., Searle, S., et al. (2012). GENCODE: the reference human genome annotation for The ENCODE Project. Genome Res 22, 1760–1774. 10.1101/gr.135350.111.

77. Durinck, S., Spellman, P.T., Birney, E., and Huber, W. (2009). Mapping identifiers for the integration of genomic datasets with the R/Bioconductor package biomaRt. Nat Protoc 4, 1184–1191. 10.1038/nprot.2009.97.

78. Fang, Z., Liu, X., and Peltz, G. (2023). GSEApy: a comprehensive package for performing gene set enrichment analysis in Python. Bioinformatics 39. 10.1093/bioinformatics/btac757.

79. Liberzon, A., Birger, C., Thorvaldsdottir, H., Ghandi, M., Mesirov, J.P., and Tamayo, P. (2015). The Molecular Signatures Database (MSigDB) hallmark gene set collection. Cell Syst 1, 417–425. 10.1016/j.cels.2015.12.004.

