## Supplementary Figure 1 for "Transcription Factor Subtype Governs Response and Resistance to DLL3-Directed T-Cell Engagement in Small Cell Lung Cancer"

### Figure S1. Early Tumor Fraction Dynamics and APEX Transcriptional Profiling in SCLC Plasma Samples

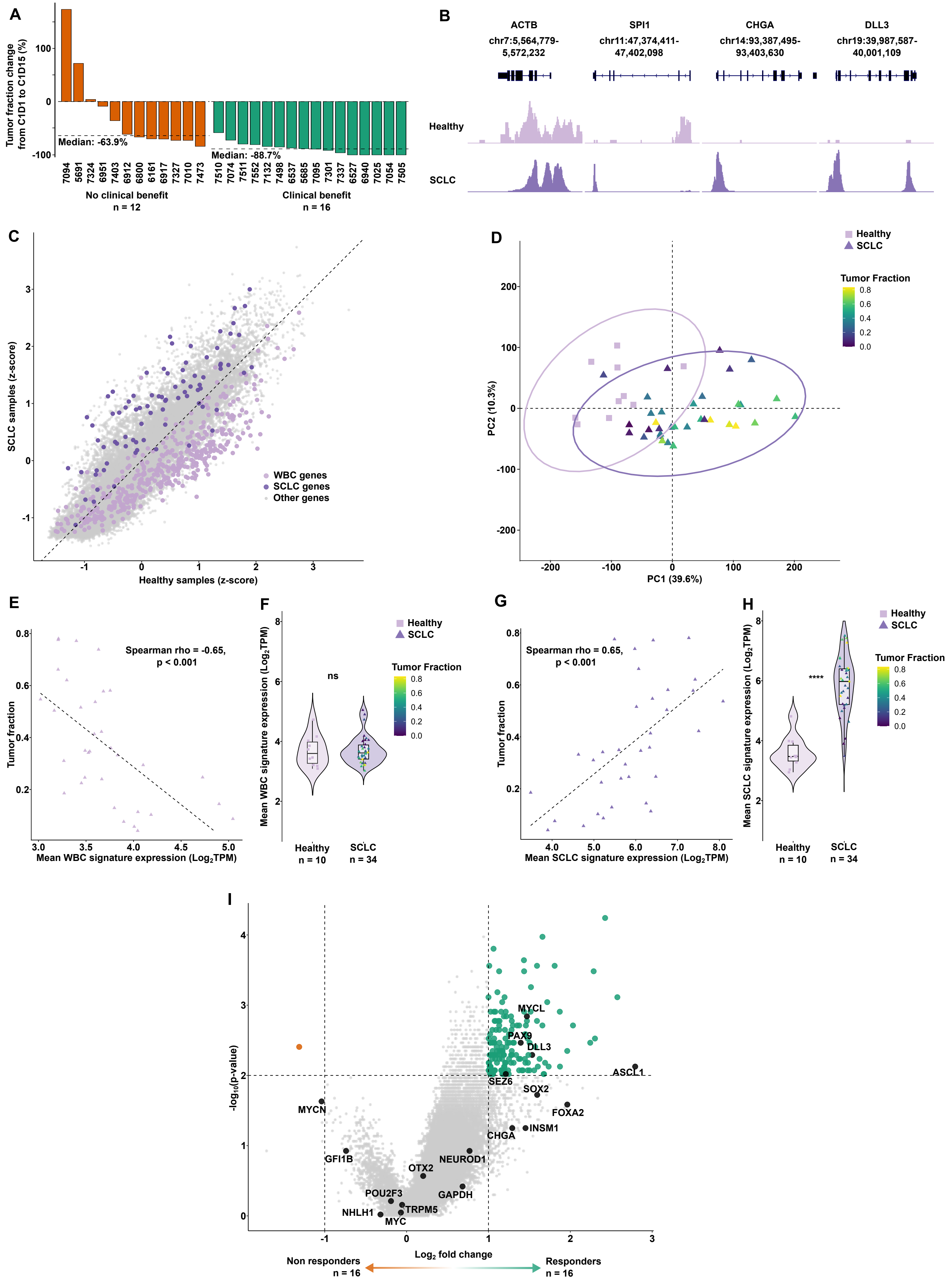
