## Supplementary Figure 2 for "Transcription Factor Subtype Governs Response and Resistance to DLL3-Directed T-Cell Engagement in Small Cell Lung Cancer"

### Figure S2. Tarlatamab Response is Correlated with SCLC Transcription Factor Subtype

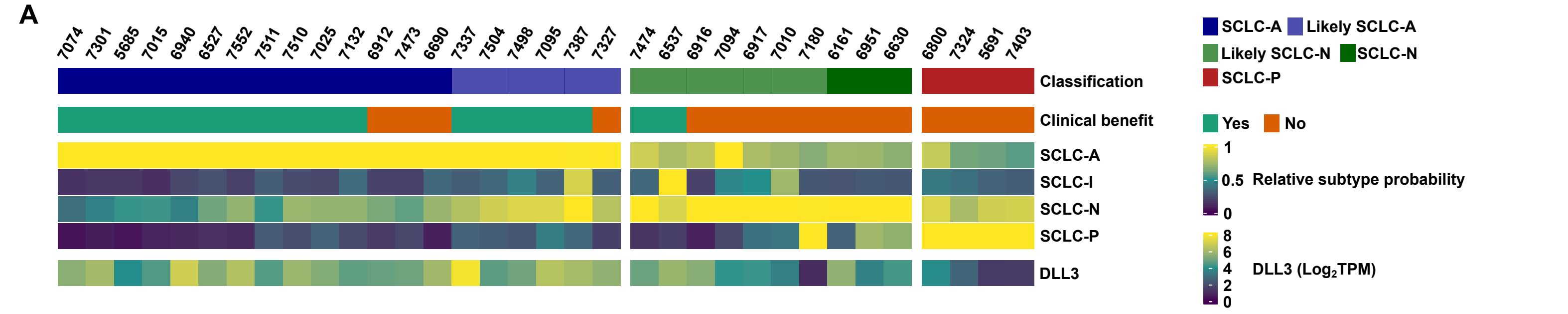


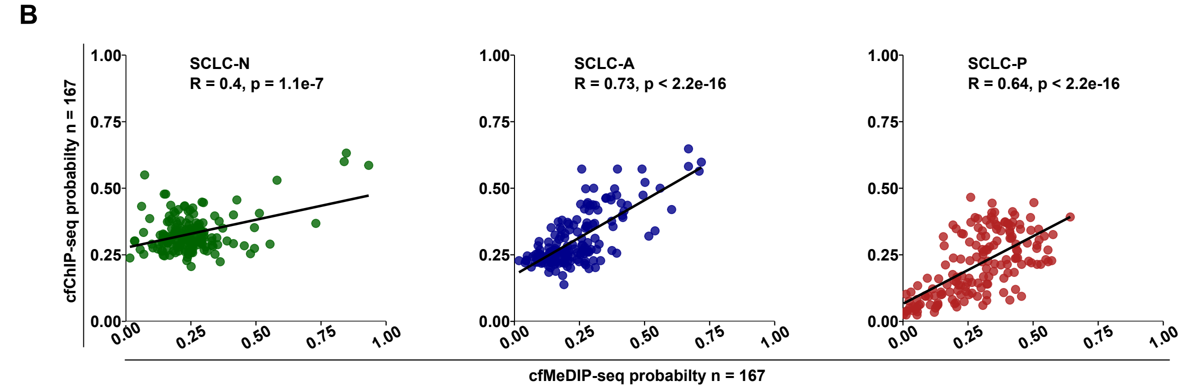


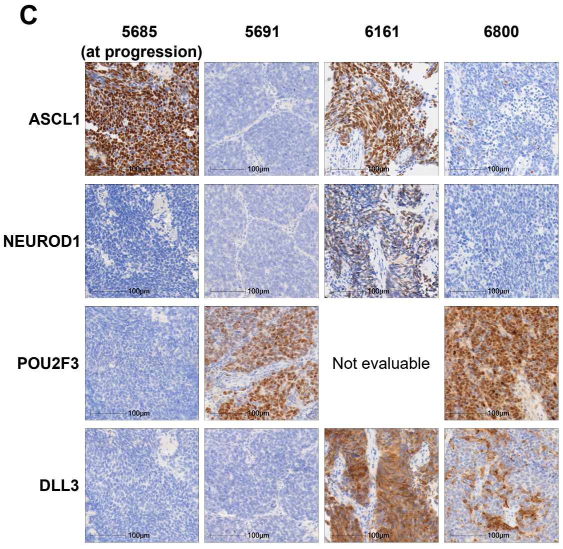


D

| Patient | ASCL1 (%) | NEUROD1 (%) | POU2F3 (%) | DLL3 (%) | cfChIP-seq Classification |
|---|---|---|---|---|---|
| 5685 (progression) | 95 | 0 | 0 | 0 | SCLC-A |
| 5691 | 0 | 0 | 90 | 0 | SCLC-P |
| 6161 | 90 | 60 | Not evaluable | 95 | SCLC-N |
| 6800 | 5 | 0 | 85 | 15 | SCLC-P |
| 6912 | 90 | <1 | 1 | 65 | SCLC-A |
| 6917 | 95 | 1 | 0 | 10 | Likely-N |

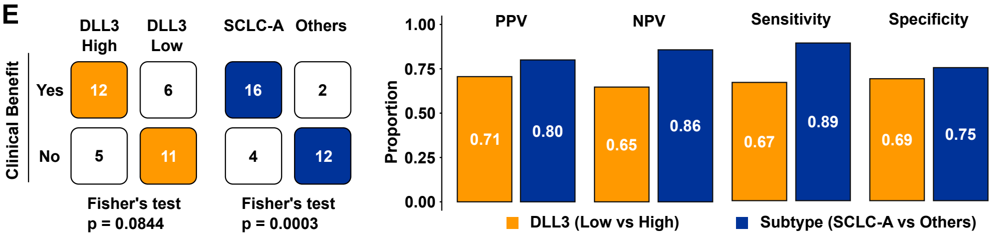
