## Supplementary figures and images for "Transcription Factor Subtype Governs Response and Resistance to DLL3-Directed T-Cell Engagement in Small Cell Lung Cancer"

### Supplementary Figure 3

#### Figure S3. Mechanisms of Acquired Resistance to Tarlatamab in Human SCLC

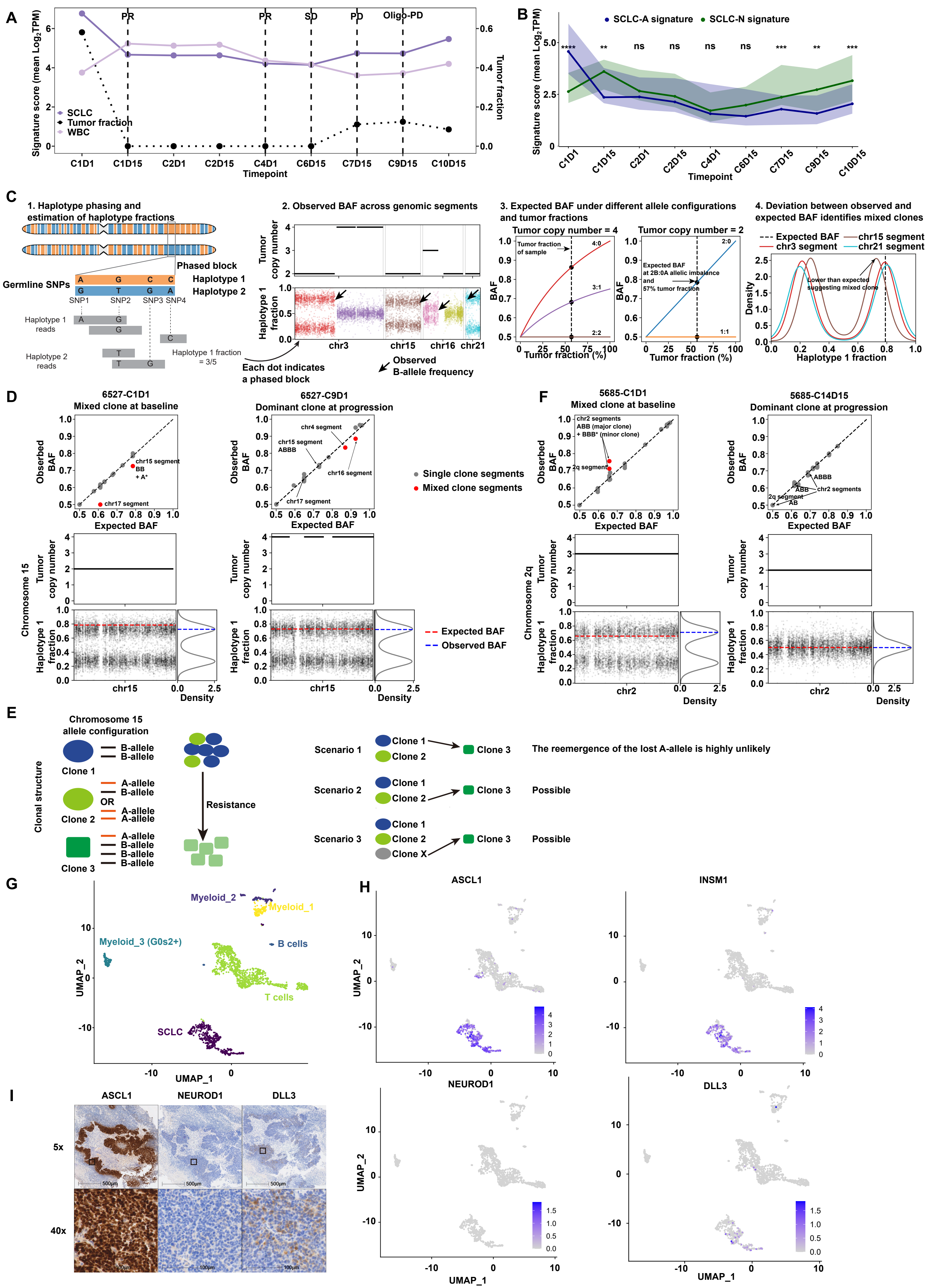

### Supplementary Figure 5

**Figure S5. NEUROD1 Promotes DLL3 Downregulation Under Tarlatamab Selective Pressure *In Vivo***

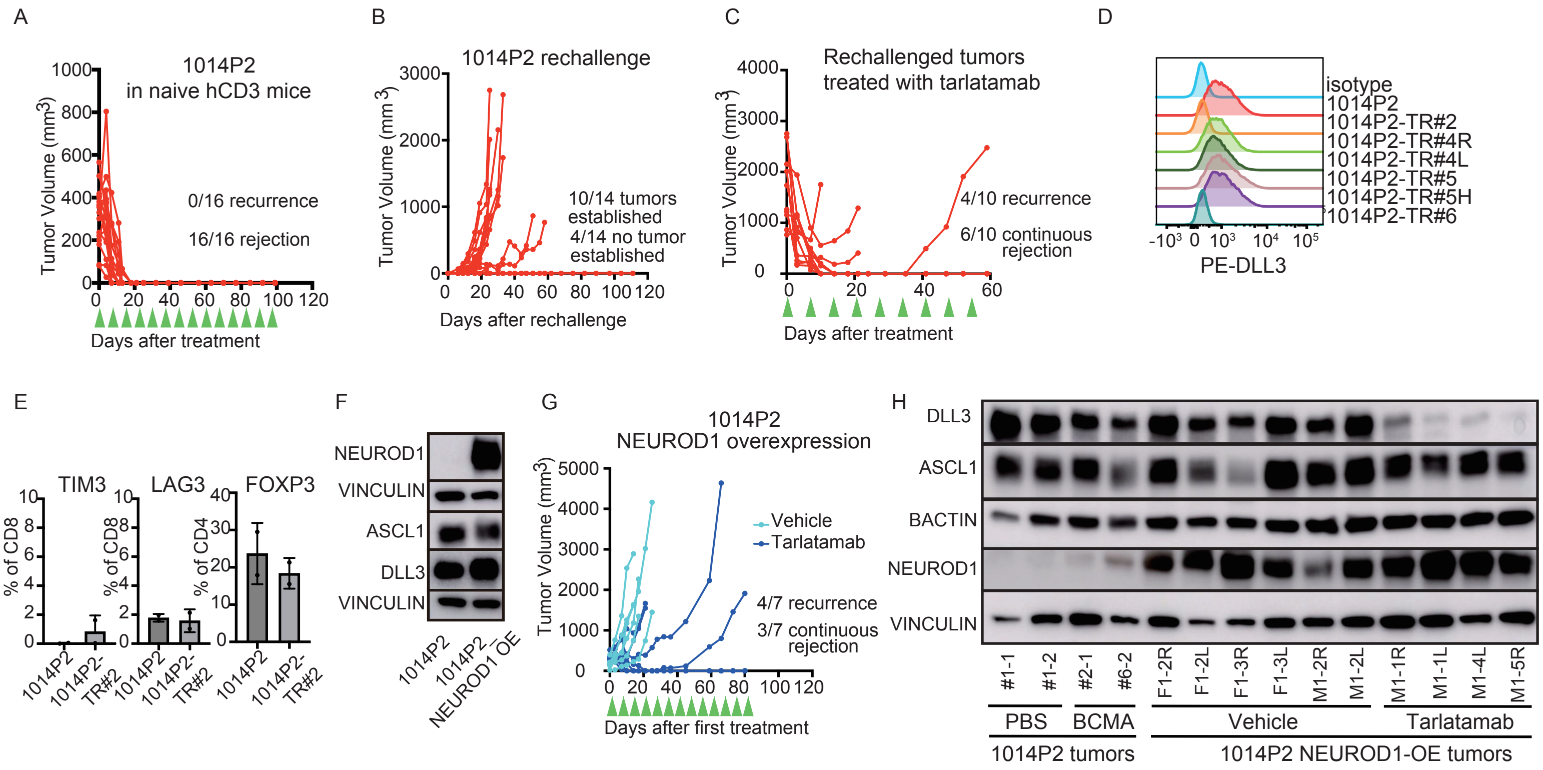
