## Supplementary Figure 4 for "Transcription Factor Subtype Governs Response and Resistance to DLL3-Directed T-Cell Engagement in Small Cell Lung Cancer"

**Figure S4. Conserved Acquired Resistance to Tarlatamab in Mice Through NEUROD1-Driven DLL3 Loss and Immune Dysfunction**

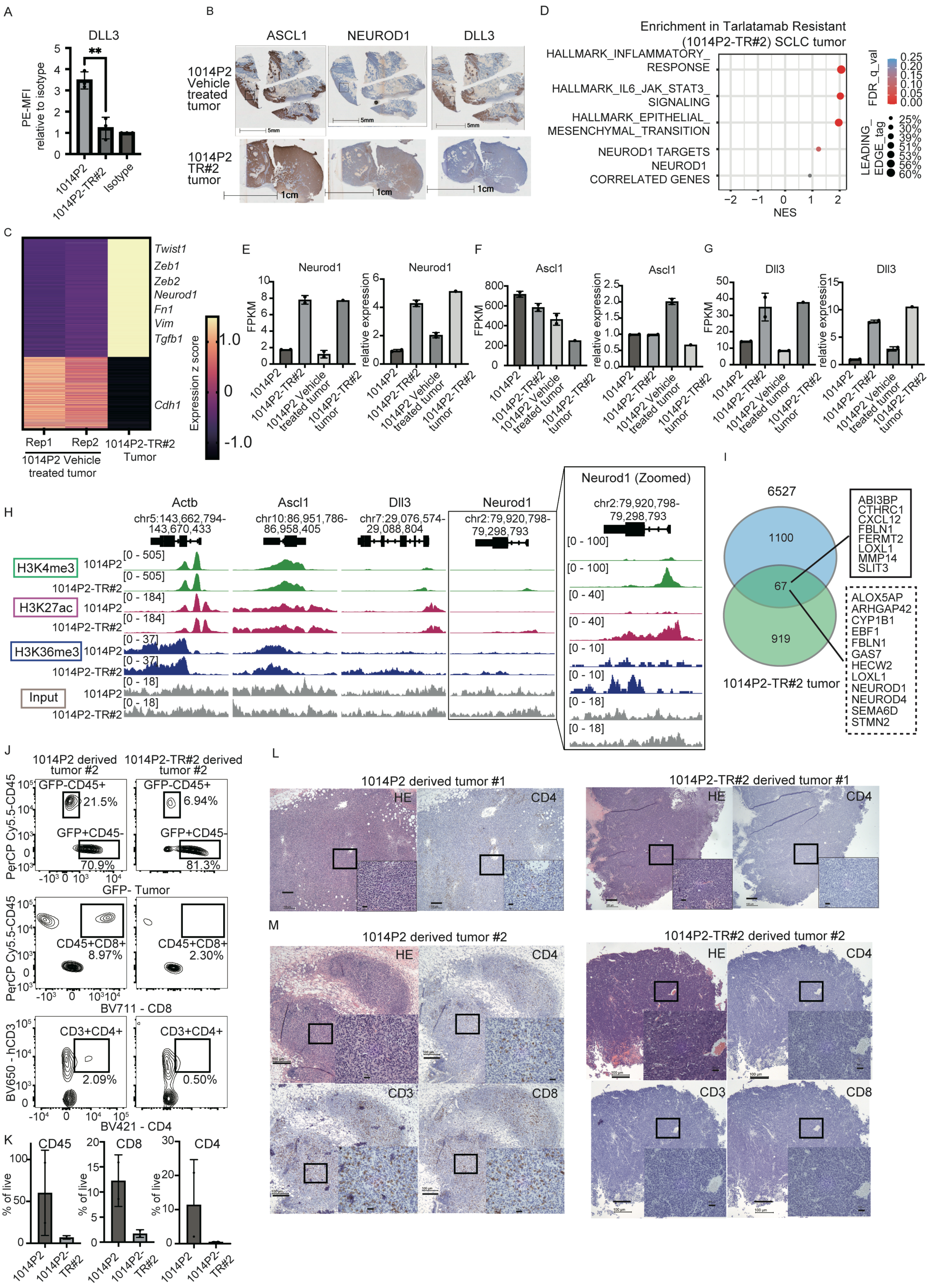
